# Ether Rhodamines with Enhanced Hydrophilicity, Fluorogenicity, and Brightness for Super-Resolution Imaging

**DOI:** 10.1101/2025.01.02.631033

**Authors:** Xiangning Fang, Qinglong Qiao, Zhifeng Li, Hao-Kai Li, Jie Chen, Ning Xu, Kai An, Wenchao Jiang, Yi Tao, Pengjun Bao, Yinchan Zhang, Zhimin Wu, Xiaogang Liu, Zhaochao Xu

## Abstract

Rhodamine dyes are widely used fluorophores in super-resolution fluorescence imaging due to their exceptional optical properties and “aggregation-disaggregation” induced fluorogenic activation. However, their excessive lipophilicity often reduces brightness in aqueous environments and causes off-target staining, limiting their effectiveness in high-resolution imaging. To address these challenges, we introduce an ether-decorated N-terminal modification strategy for rhodamine and silicon-rhodamine (Si-rhodamine), replacing conventional N-alkyl groups. The ether chains enhance water solubility, decrease aggregate size, and improve fluorogenicity across a wide concentration range. Their flexible, hydrophilic structure forms a protective shield around the xanthene core, minimizing dye-water interactions and reducing fluorescence quenching. Additionally, the inductive effect of the ether chains decreases the electron-donating strength of the amino groups, suppressing quenching caused by twisted intramolecular charge transfer (TICT). These modifications collectively increase the quantum yields of **ER** and **ESiR** in water from 0.35 and 0.19 (for tetraethyl-substituted analogs) to 0.70 and 0.41, respectively. Probes derived from **ER** and **ESiR** exhibit outstanding fluorogenicity, enhanced signal-to-noise ratios, and improved resolution in complex aqueous environments, demonstrating superior performance in advanced super-resolution imaging techniques such as structured illumination microscopy (SIM), stimulated emission depletion (STED) microscopy, and single-molecule localization microscopy (SMLM). This work introduces an innovative strategy for fluorophore design, offering significant advancements for super-resolution imaging applications.

## Introduction

Advancements in super-resolution imaging have propelled significant progress in dye chemistry, necessitating stringent optimization of fluorescent dyes’ photophysical properties.^1-9^ Techniques such as stimulated emission depletion (STED) microscopy demand dyes capable of withstanding high-intensity depletion lasers without rapid photobleaching,^10-12^ while single-molecule localization microscopy (SMLM) requires dyes that are bright, photostable, and capable of fluorescence switching.^13-19^ Addressing these stringent requirements has led to significant innovations in dye design.^20-25^ These advancements have also catalyzed progress in fluorogenic sensing^26-28^ and multicolor imaging,^29-33^ paving the way for detailed exploration of cellular structures and dynamics at the nanoscale.^4-5, 28, 34-40^

Among the many fluorescent dyes available, rhodamine derivatives are particularly notable due to their tunable spectral properties,^4, 41-43^ high brightness,^13, 44^ and remarkable photostability.^35^ A key feature of rhodamine dyes is their dynamic equilibrium between the non-fluorescent spirolactone and fluorescent zwitterion forms,^21, 45-46^ which enables controlled fluorescence turn-on and blinking. This property makes rhodamine dyes highly suitable for applications like fluorogenic labeling and SMLM. However, the hydrophobic nature of rhodamine often leads to aggregation in its closed-ring form within the cytoplasm.^47-48^ When functional groups on rhodamine, such as HaloTag substrates, engage in molecular recognition with targets, these aggregates may dissociate, driven by electrostatic interactions with the protein binding sites.^49-51^ These interactions could stabilize rhodamine in its fluorescent zwitterion state and enable fluorogenic imaging (Scheme 1a).^19, 23, 25-26, 52-53^ While a small degree of aggregation facilitates disaggregation-induced fluorogenicity, excessive aggregation can cause non-specific binding and lead to background fluorescence. These issues are particularly problematic in sparse labeling scenarios, where background signals can overwhelm the fluorescence from labeled targets. Furthermore, most biomolecular processes occur in aqueous environments, and the attachment to a long linker often causes the probe to remain outside narrow hydrophobic cavities post-labeling. ^49, 54-56^ These challenges underscore the importance of maintaining high fluorescence brightness in aqueous environments and controlling rhodamine aggregation behavior to preserve its fluorogenic characteristics while minimizing background noise. Addressing these limitations necessitates the development of novel rhodamine derivatives optimized for super-resolution imaging.

Fluorescence quenching of dyes in aqueous environments arises from multiple mechanisms, often limiting their performance in biological imaging (Scheme 1a). Poor water solubility leads to aggregation, resulting in strong intermolecular interactions that quench fluorescence.^57-61^ One common approach to enhance the water solubility of rhodamine dyes and reduce non-specific cellular binding is incorporating sulfonate groups into their structures.^62-64^ However, the negatively charged sulfonate groups can cause electrostatic repulsion with the negatively charged cell membrane, hindering the penetration of these dyes into live cells.^65^ Furthermore, sulfonate-modified rhodamine dyes that remain in their open-ring form disrupt the aggregation-disaggregation fluorogenicity. Another major quenching mechanism is external conversion, where energy from the excited fluorophore is transferred to surrounding water molecules upon collisions.^66-67^ Using deuterated water has been shown to effectively increase fluorescence intensity by reducing collisions between water molecules and the fluorophore.^68-70^ However, this expensive approach raises concerns about its potential effects on cellular physiology, limiting its practical use. Twisted intramolecular charge transfer (TICT) is another significant quenching pathway, particularly in high-polarity environments.^71^ Strategies to mitigate TICT include increasing the TICT energy barrier^13, 44^ by rigidifying the N-alkyl group through structural modifications, such as incorporating azetidine rings to minimize C-N bond rotation.^13^ Alternatively, substituting the N-alkyl group with quaternary piperazine or sulfone-functionalized piperidine reduces the electron-donating ability of the N-substituent, further enhancing brightness and photostability^44, 72^ (Scheme 1a). While these strategies improve rhodamine’s performance in polar environments, they often exacerbate its tendency to aggregate in water, leading to non-specific background signals. To avoid such aggregation-induced artifacts, probes typically require careful concentration control, generally below 50-200 nM,^25^ during super-resolution imaging. However, this limitation significantly hampers their application, particularly for dynamic imaging or targeting regions with high intracellular density. Thus, there is a pressing need for innovative strategies that can simultaneously address these three fluorescence quenching pathways while retaining the benefits of aggregation-disaggregation-induced fluorogenicity. Such approaches would enable super-resolution imaging with reduced dependency on dye concentration, enhancing versatility and applicability.

In this study, we report the development of rhodamine dyes with tailored hydrophilicity and enhanced brightness, specifically N-ether-substituted rhodamine and silicon rhodamine (Si-rhodamine), designated as **ER** and **ESiR**, respectively. Incorporating ether chains significantly improves the quantum yields of these dyes in aqueous environments while substantially enhancing their solubility and reducing their tendency to aggregate. Importantly, these modifications preserve the aggregation-disaggregation-induced fluorogenic sensing mechanism, resulting in concentration-independent fluorogenicity that outperforms hydrophobic rhodamine derivatives. Accordingly, the newly developed probes exhibit nearly double the brightness and signal-to-noise ratio (SNR) in live-cell imaging compared to the conventional N-alkyl rhodamine-based dyes, enabling their effective use in dynamic live-cell imaging and super-resolution techniques such as structured illumination microscopy (SIM) and STED microscopy. Furthermore, by combining this approach with spirolactam modifications, we successfully created a lysosomal probe capable of monitoring long-term lysosomal dynamics with exceptional precision via SMLM.

**Scheme 1.**
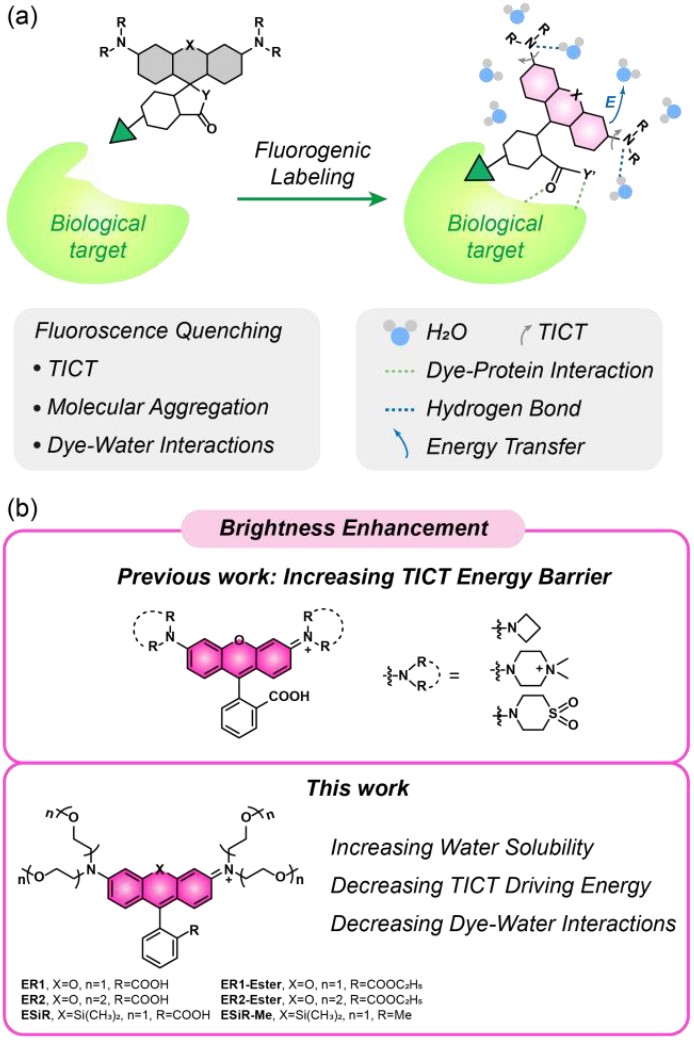
(a) Factors that affect rhodamine probes in fluorogenic labeling and imaging. (b) Design and structures of rhodamines to enhance brightness.

## RESULTS AND DISCUSSION

### Molecular design and photophysical properties of ether rhodamines

Ether linkages, known for their neutral and hydrophilic nature, provide excellent biocompatibility and exhibit low non-specific interactions within cells. These characteristics make them particularly suitable for applications in biomaterials and drug delivery systems.^73-74^ By substituting the traditional N-alkyl group with an ether linkage at the nitrogen position of the rhodamine chromophore, we could effectively enhance the water solubility of rhodamine. The ether group could also act as a “protective barrier,” wrapping around the rhodamine molecule to reduce interactions with water molecules. This mechanism minimizes external conversion, a key fluorescence quenching pathway (Scheme 1b). Importantly, ether linkages exert minimal impact on the electronic delocalization of the rhodamine chromophore without direct conjugation. As a result, the critical dynamic equilibrium between the non-fluorescent spirolactone form and the fluorescent zwitterionic form is preserved. This balance is essential for maintaining the fluorogenic properties of rhodamine derivatives, ensuring their effectiveness in advanced imaging applications.

Building on this rationale, we incorporated ether linkages containing one or two oxygen atoms into rhodamine and Si-rhodamine derivatives, leading to the successful synthesis of **ER1, ER2**, and **ESiR** (Scheme 1b). However, attempts to synthesize the Si-rhodamine derivative with two oxygen atoms in the ether linkage were unsuccessful.

We subsequently evaluated the photophysical properties of the synthesized compounds (Figure 1a-b, S1, Table S1-3). Compared to N-tetraethyl rhodamines (**RhoB** and **TESiR**), introducing ether linkages resulted in slight hypochromic shifts in both λ_abs_ and λ_em_ values. Notably, these modifications substantially improved quantum yields and brightness in aqueous environments (Table 1). **ER1**, which contains one ether group on each N-substituent, exhibited a significantly higher quantum yield (*Φ* = 0.61) than **RhoB** (*Φ* = 0.35). Extending the length of the ether linkages in **ER2** further enhanced the quantum yield to 0.70, doubling that of **RhoB**. Similarly, **ESiR** displayed a quantum yield (*Φ* = 0.41) more than twice as high as **TESiR**’s (*Φ* = 0.19). Consequently, the brightness of these ether-modified rhodamines in water was markedly higher than that of their N-tetraethyl counterparts, underscoring the effectiveness of ether linkages in enhancing fluorophore performance.

**Table 1.** Photophysical properties of ether rhodamines and N-alkyl rhodamines. (* Data from published reference.^57, 75^)

| Dye | $\lambda_{\text{abs}}$<br>(nm) | $\lambda_{\text{em}}$<br>(nm) | $\Delta\lambda$<br>(nm) | $\epsilon$<br>(M<sup>-1</sup> cm<sup>-1</sup>) | $\Phi$ | Brightness<br>($\epsilon \times \Phi$) |
| --- | --- | --- | --- | --- | --- | --- |
| **ER1** | 545 | 570 | 25 | 69265 | 0.61 | 42252 |
| **ER2** | 545 | 571 | 26 | 65770 | 0.70 | 46039 |
| **ER1-Etster** | 550 | 577 | 27 | 72413 | 0.50 | 36207 |
| **ER2-Etster** | 551 | 577 | 26 | 74565 | 0.56 | 41756 |
| **ESiR** | 643 | 664 | 21 | 28005 | 0.41 | 11482 |
| **ESiR-Me** | 648 | 665 | 17 | 112410 | 0.33 | 37095 |
| **RhoB\*** | 554 | 577 | 23 | 111000 | 0.35 | 38850 |
| **TESiR\*** | 650 | 666 | 16 | 50000 | 0.19 | 9500 |

**Figure 1.**
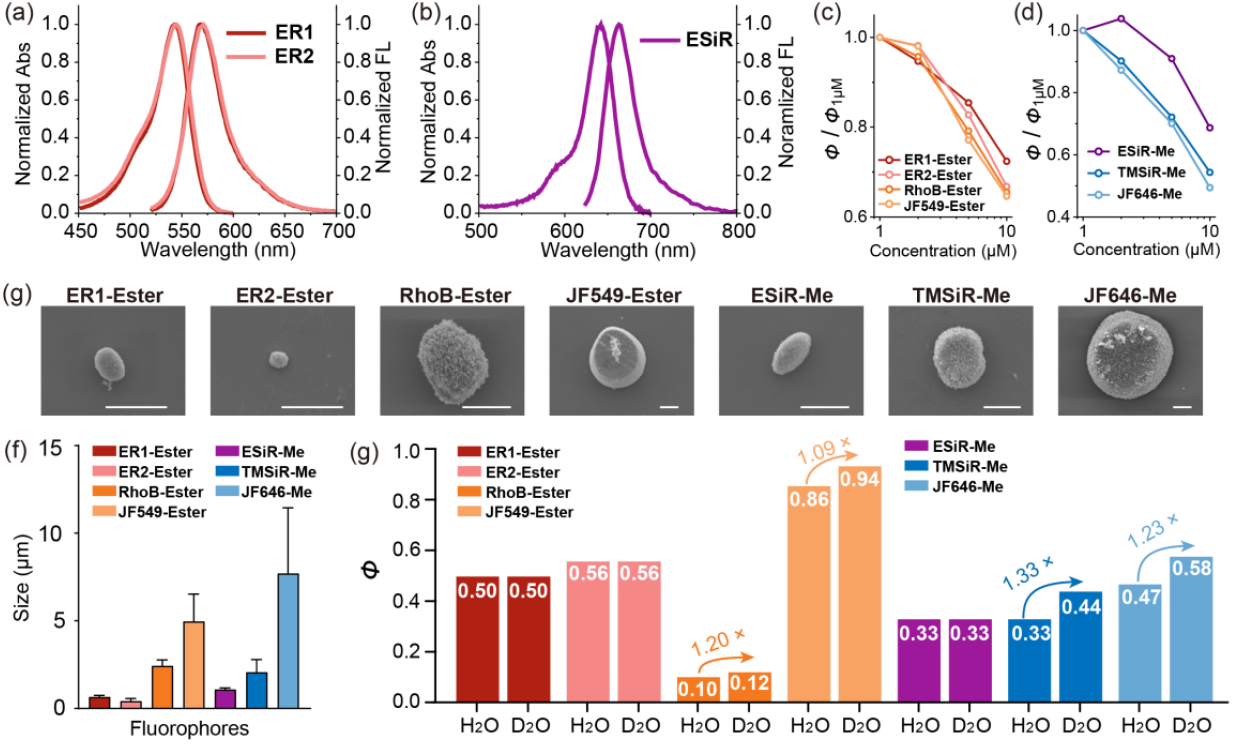
Photophysical properties of ether rhodamines in water. (a) Normalized UV-Vis and fluorescence spectra of **ER1** and **ER2** in H_2_O (2 μM). (b) Normalized UV-Vis and fluorescence spectra of **ESiR** in H_2_O (2 μM). (c, d) The trends of quantum yield changes under different dye concentrations ranging from 1 μmol/L to 10 μmol/L in PBS (10 mM). (e) SEM imaging of aggregates of ether rhodamines, N-alkyl rhodamines, and azetidine rhodamines. Scale bar, 1 μm. (f) Particle sizes of different fluorophores under SEM (*n* = 4). (g) Quantum yields of ether rhodamines, N-alkyl rhodamines, and azetidine rhodamines in H_2_O and deuterated water.

We hypothesized that the hydrophilic ether linkages would improve the solubility of fluorophores in aqueous environments. To test this, we measured the quantum yields of ether rhodamines across a range of concentrations (1 to 10 µM) in phosphate-buffered saline (PBS, 10 mM), as aggregation of rhodamines typically reduces quantum yield. To eliminate the equilibrium between the non-fluorescent lactone and fluorescent zwitterionic forms of rhodamines, we esterified the carboxyl group (**ER1-Ester, ER2-Ester**) or substituted it with a methyl group (**ESiR-Me**) and examined their photophysical properties (Figure S2, Table S4-6). Using the reference quantum yield at 1 µM, we calculated relative quantum yields for other concentrations (Figure 1c-d, Table S7). For comparison, we evaluated open-formed N-alkyl and azetidine rhodamines (**RhoB-Ester, JF549-Ester, TMSiR-Me, JF646-Me**) as shown in Figure S3 to assess the solubility advantage of ether rhodamines. All dyes exhibited a decline in quantum yield at higher concentrations due to aggregation. However, the decrease in quantum yield for **ER1-Ester, ER2-Ester**, and **ESiR-Me** (72%, 67%, 69%) was significantly smaller than that observed for N-alkyl and azetidine rhodamines counterparts (65% of **RhoB-Ester**, 64% of **JF549-Ester**, 54% of **TMSiR-Me**, 49% of **JF646-Me**). This contrast demonstrates the improved solubility of ether rhodamines, attributed to the ability of ether chains to reduce dye aggregation in aqueous solutions.

It is worth noting that the observed decline in quantum yield is also partially influenced by inner filter effects. Therefore, the actual degree of aggregation in ether rhodamines is likely much lower than the calculated 72%, 67% and 69%. This further underscores the effectiveness of ether linkages in mitigating aggregation and maintaining higher quantum yields in aqueous environments.

To directly evaluate the aggregation behavior of these fluorophores in aqueous solution, we used scanning electron microscopy (SEM) to observe the aggregates formed by the fluorophores. Aqueous solutions of the fluorophores (5 μM) were deposited on silicon wafers, and after natural drying, SEM imaging was conducted to measure the sizes of the resulting particles (Figure 1e-f). The fluorophores primarily aggregated into spherical particles. Notably, the spheres formed by ER1-Ester, **ER2-Ester**, and **ESiR-Me** were significantly smaller (651 nm, 425 nm, 1080 nm) compared to those formed by N-alkyl rhodamines and azetidine rhodamines (2430 nm of **RhoB-Ester**, 4974 nm of **JF549-Ester**, 2065 nm of **TMSiR-Me**, 7703 nm of **JF646-Me**). This reduction in aggregate size demonstrates that the ether chains enhance the water solubility of rhodamines, leading to improved dispersion in aqueous environments. These results provide further evidence that ether linkages effectively reduce aggregation, contributing to better photophysical properties in water.

In addition to enhancing water solubility, we hypothesized that ether linkages might minimize interactions between fluorophores and water molecules, thereby suppressing collisional quenching. Collisions with water molecules can facilitate external conversion, where energy from excited-state fluorophores is transferred to water molecules, leading to fluorescence quenching.^66-67^ This effect can be mitigated by substituting H_2_O with D_2_O, as deuterium suppresses vibrational interactions.^76^ To investigate this, we measured the quantum yields of open-formed ether rhodamines in both H_2_O and D_2_O (Figure 1g). In both solvents, the quantum yields of ether rhodamines remained nearly consistent (e.g., *Φ* values in H_2_O and D_2_O: 0.50 for ER1-Ester, 0.56 for **ER2-Ester**, and 0.33 for **ESiR-Me**). In contrast, the quantum yields of N-alkyl and azetidine rhodamines improved significantly in D_2_O. For example, **RhoB-Ester** and **JF549-Ester** showed 1.20-fold and 1.09-fold increases, respectively, while Si-rhodamine derivatives **TMSiR-Me** and **JF646-Me** exhibited even more significant improvements, with 1.33-fold and 1.23-fold increases, respectively. The more pronounced improvement in Si-rhodamine quantum yields is likely due to its smaller S0-S1 energy gap, which is closer to the vibrational energy of the OH group,^67^ making quenching by water more severe.

The consistency of quantum yields for ether rhodamines in H_2_O and D_2_O suggests negligible energy transfer from the excited-state ether rhodamines to water molecules. These results indicate that ether linkages provide a protective effect, shielding the fluorophore from the solvent environment and minimizing external conversion. This protective capability further underscores the advantages of incorporating ether linkages into rhodamine fluorophore.

#### Quantum chemical calculations

Incorporating ether linkages at the N-terminal position of rhodamines significantly enhances their hydrophilicity and brightness. Traditionally, improvements in rhodamine brightness have focused on increasing the TICT energy barrier.^13, 44^ However, other factors, such as driving energy and the local environmental polarity,^66, 71, 77^ also strongly influence TICT but have often been overlooked in previous studies. We performed quantum chemical calculations to better understand the mechanisms behind the enhanced quantum yields of ether rhodamines. These calculations aimed to elucidate the contributions of ether linkages to the photophysical properties of rhodamines.

We evaluated the electron-donating strength of ether substituents relative to conventional amino donors by calculating their ionization potentials (Figure 2a). A phenyl group was introduced to “passivate” the substituents during the calculations to ensure a fair comparison. The results showed that ether substituents exhibit higher ionization potentials than dialkyl-amino groups, indicating reduced electron-donating capacity. These findings were further validated using alternative calculation methods (i.e., passivation using a methyl group) with consistent results (Figure S4).

**Figure 2.**
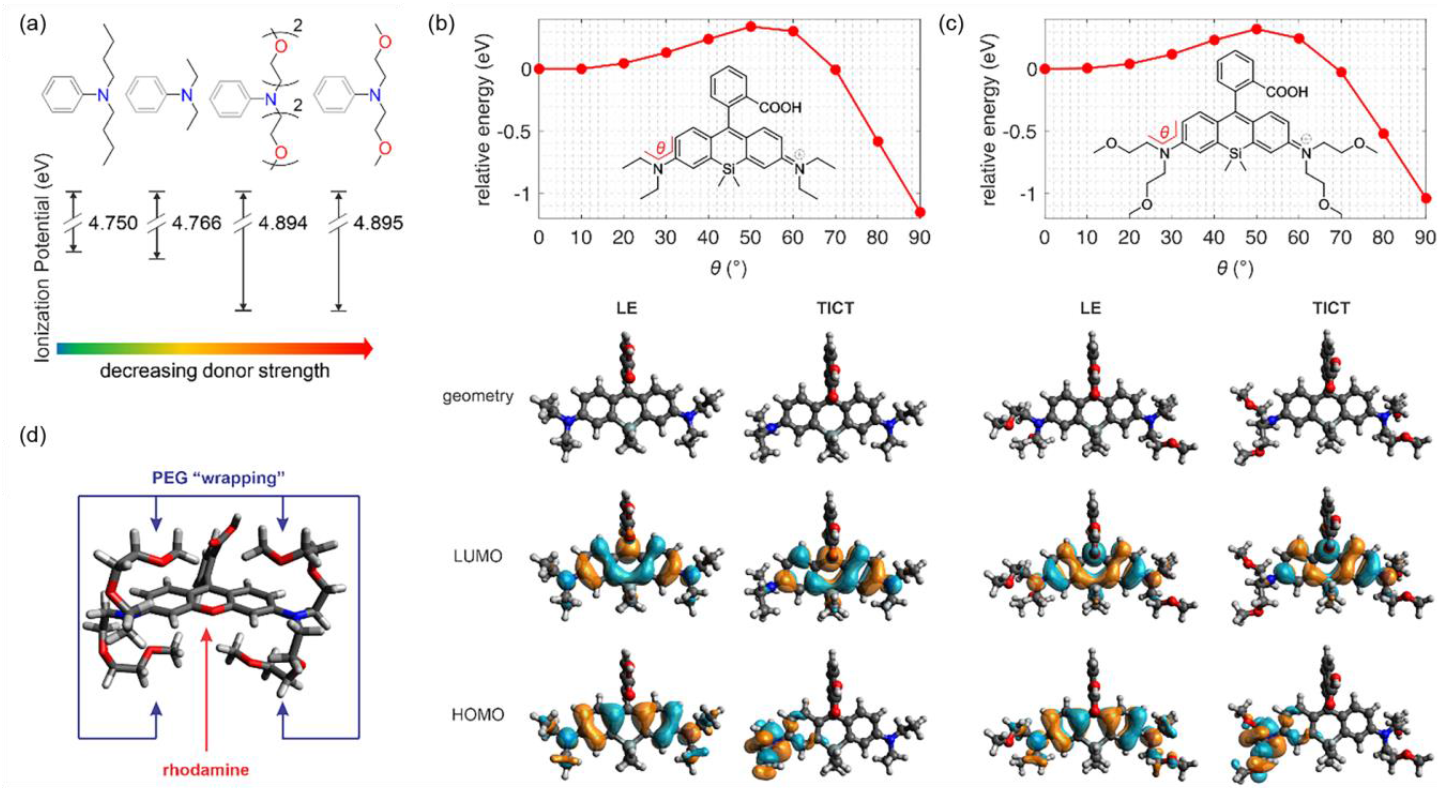
(a) Calculated ionization potential of various amino substituents in water. Relative electronic energy of the S_1_ potential energy surface of (b) **TESiR** and (c) **ESiR** as a function of *θ*. The bottom panel shows the geometries, HOMO, and LUMO of these dyes in the locally excited (LE; *θ* = 0°) and TICT states (*θ* = 90°), respectively. (d) The most stable conformation of **ER2**.

The reduced electron-donating ability of ether substituents, compared to N-alkyl groups, can be attributed to the electron-withdrawing nature of the oxygen atom. The oxygen atom (more electronegative) draws electrons away from adjacent carbon atoms (less electronegative) through inductive effects, thereby weakening their electron-donating strength. This characteristic explains the observed slight blue shift in the absorption maxima of **ER1, ER2** and **ESiR** relative to their N-alkyl-substituted counterparts, with a change of approximately 9 nm in water.

The reduced donor strength of ether-substituted rhodamine dyes increases their resistance to TICT state formation. To investigate this effect, we modeled the potential energy surfaces of these rhodamine derivatives in water to simulate the TICT process. While the energy barriers for TICT rotations were similar across compounds, significant differences were observed in driving energies. For example, the ether rhodamine **ESiR** exhibited a lower driving energy (1.04 eV) than the reference **TESiR** (1.15 eV). Similarly, **ER1** displayed a reduced driving energy (1.14 eV) relative to **RhoB** (1.24 eV) (Figure 2b-c, S5). These reductions in driving energy indicate enhanced resistance to TICT formation, contributing to the higher quantum yields of ether rhodamines.

The conformational analysis further revealed that while ether groups could adopt extended conformations (allowing free rotation of the amino group during TICT), they predominantly stabilize in a “wrapped” configuration around the rhodamine core (Figure 2d; Figure S6). This “wrapping” effect mimics the dissolution of rhodamines in ether molecules and has several key implications: (1) it substantially reduces local polarity, inhibits TICT rotations—which are exacerbated in polar environments like water; (2) the wrapped structure sterically hinders amino group rotations, further suppressing TICT formation; (3) additionally, the wrapped structure minimizes collisions between the dye and water molecules, effectively reducing external conversion. This finding is consistent with the experimentally observed stability of quantum yields in ether rhodamines when measured in both H_2_O and D_2_O.

This discovery also aligns with previous findings, where encapsulating rhodamines within supramolecular cucurbit[7]uril (**CB7**) effectively reduced local polarity and diminished collisions between the dye and the solvent, improving quantum yields and photostability.^78^ Notably, while **CB7** encapsulation has decreased the water solubility of dye-**CB7** complexes, the ether wrapping method enhances water solubility.

In summary, the combined effects of TICT suppression, reduced external conversion, and enhanced solubility— which effectively minimizes aggregation—synergistically contribute to the significantly improved quantum yields observed in ether rhodamines.

#### Concentration-independent Living-cell imaging

Inspired by the exceptional photophysical properties of these novel fluorophores, we utilized them for living-cell fluorogenic imaging. **ER1** and **ESiR** demonstrated excellent cell permeability, while **ER2** was impermeable, likely due to the excessively long ether chains (Figure S7). We thus focused on **ER1** and **ESiR** derivatives for developing fluorescent probes. Considering that rhodamine-based HaloTag probes stabilize in the fluorescent zwitterion state upon binding to HaloTag, thereby providing strong fluorogenic imaging, we synthesized **ER-Halo** and **ESiR-Halo** for HaloTag labeling in living cells (Figure 3a, S8, Table S8).

**Figure 3.**
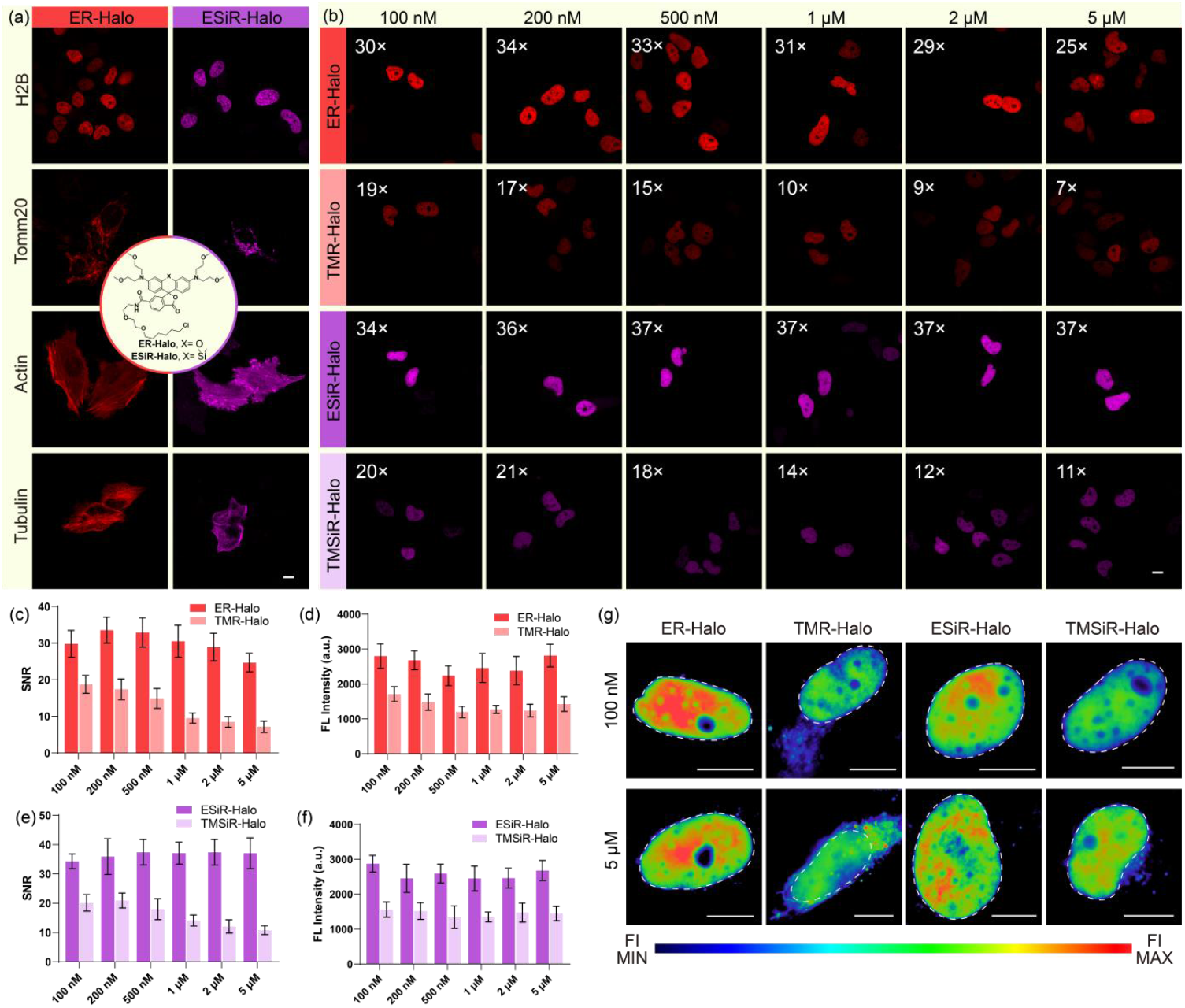
Ether rhodamines-based probes for fluorogenic imaging. (a) Confocal imaging of different organelles (nucleus, mitochondria, actin and tubulin) expressing HaloTag using **ER-Halo** (left) and **ESiR-Halo** (right). HeLa cells transiently expressing Halo-H2B, Halo-Tomm20, Halo-Actin and Halo-Tubulin respectively, were incubated with 100 nM probes for 1 h. (b) Confocal imaging and SNR of nuclei using **ER-Halo, TMR-Halo, ESiR-Halo** and **TMSiR-Halo** under different probe concentration. (c, d) The comparison of SNR (c) and fluorescent intensity (d) between **ER-Halo** and **TMR-Halo** in nuclei imaging under different concentration. (e, f) The comparison of SNR (e) and fluorescent intensity (f) between **ESiR-Halo** and **TMSiR-Halo** in nuclei imaging under different concentration. The SNR data and FL Intensity data of each probe was derived from 15 nuclei. (g) SIM imaging of nuclei using **ER-Halo, TMR-Halo, ESiR-Halo** and **TMSiR-Halo** under different probe concentrations. Scale bar, 10 μm

We first assessed the reactivity of **ER-Halo** and **ESiR-Halo** with purified HaloTag in vitro. The labeling reaction for **ER-Halo** was completed instantaneously, while **ESiR-Halo** required approximately 30 minutes to complete (Figure S9). The quantum yields of both **ER-Halo** and **ESiR-Halo** were higher than those of N-methyl rhodamine-based HaloTag probes **TMR-Halo** and **TMSiR-Halo** (Figure S10), regardless of whether they were bound to HaloTag (Table 2). These results demonstrate that ether rhodamines maintain high quantum yields even after functional modification.

**Table 2.** Quantum yields of HaloTag probes (1 μM) before and after reactions with HaloTag (2 μM).

| Probe | Probe in PBS | Probe + HaloTag |
| --- | --- | --- |
| **ER-Halo** | 0.51 | 0.74 |
| **TMR-Halo** | 0.43 | 0.57 |
| **ESiR-Halo** | 0.27 | 0.29 |
| **TMSiR-Halo** | 0.19 | 0.23 |

Next, we evaluated the biocompatibility of **ER-Halo** and **ESiR-Halo** by incubating them with living cells, where both probes exhibited high compatibility (Figure S11). We transfected commercial plasmids to express HaloTag in various organelles, including the nucleus, mitochondria, actin, and tubulin (Table S9). Following incubation with **ER-Halo** and **ESiR-Halo**, confocal imaging revealed strong fluorescence signals localized to the targeted organelles (Figure 3a, S12). These results highlight the excellent targeting accuracy of **ER-Halo** and **ESiR-Halo** for HaloTag and their universal labeling capability across different organelles.

We next evaluated the living-cell imaging performance of **ER-Halo** and **ESiR-Halo** compared to N-methyl rhodamine-based HaloTag probes (**TMR-Halo** and **TMSiR-Halo**). For this purpose, we transfected HeLa cells with Halo-H2B to assess fluorogenic labeling and imaging results within the nuclei.

First, we incubated transfected HeLa cells with **ER-Halo** and **TMR-Halo** under varying conditions, with probe concentrations ranging from 100 nM to 5 μM (Figure 3b). Across these conditions, **ER-Halo** demonstrated a significantly higher average nuclei-to-cytosol signal-to-noise ratio (SNR), ranging from 33-fold to 25-fold, compared to **TMR-Halo**, which exhibited an SNR of 19-fold to 7-fold (Figure 3c). Notably, as the probe concentration increased, the SNR of **TMR-Halo** decreased markedly by ∼ 63%, whereas the SNR of **ER-Halo** decreased by only ∼17%. Furthermore, the SNR difference between **ER-Halo** and **TMR-Halo** increases significantly, from 1.6-fold at 100 nM to 3.6-fold at 5 μM. This difference highlights **ER-Halo**’s ability to sustain superior SNR even at elevated concentrations, emphasizing its outstanding fluorogenic imaging performance. Fluorescent intensity measurements extracted from nuclei further confirmed that **ER-Halo** was brighter than **TMR-Halo** in living cells (Figure 3d). These results were consistent with the in vitro findings, demonstrating the robust performance of **ER-Halo** in terms of both brightness and SNR, particularly at high concentrations.

Next, we evaluated the SNR and brightness of Si-rhodamine-based probes, comparing **ESiR-Halo** to **TMSiR-Halo**. Across a concentration range of 100 nM to 5 μM, **ESiR-Halo** maintained a stable SNR of 34–37-fold. In contrast, the SNR of **TMSiR-Halo** decreased significantly, from 21-fold to 11-fold, as the dye concentration increased. Similar to the trend observed with **ER-Halo** and **TMR-Halo**, the SNR difference between **ESiR-Halo** and **TMSiR-Halo** increased with concentration, growing from 1.7-fold at lower concentrations to 3.4-fold at higher concentrations (Figure 3e). Fluorescence intensity analysis further confirmed the superior brightness of **ESiR-Halo** compared to **TMSiR-Halo** in living cells (Figure 3f). These results demonstrate that **ESiR-Halo** consistently outperforms **TMSiR-Halo** in both SNR and brightness, even at higher dye concentrations.

To further evaluate the non-specific background fluorescence of the probes in living cells, we used SIM to analyze background fluorescence at both low (100 nM) and high (5 μM) staining concentrations. In Figure 3g, nuclei are outlined with gray dotted circles to facilitate visualization of intracellular probe distribution against background fluorescence (outside the circles). The ether rhodamine probes, **ER-Halo** and **ESiR-Halo**, exhibited negligible intracellular background fluorescence under both low and high staining conditions. In contrast, the N-alkyl rhodamine probe **TMR-Halo** displayed punctate lysosomal background fluorescence even at 100 nM. At the higher concentration of 5 μM, **TMR-Halo** showed pronounced lysosomal and mitochondrial background fluorescence, with background brightness comparable to that of the nucleus. While **TMSiR-Halo** exhibited minimal background fluorescence at 100 nM, it demonstrated noticeable lysosomal fluorescence at 5 μM. Additionally, SIM imaging confirmed that the brightness of ether rhodamine probes consistently surpassed that of N-alkyl rhodamine probes across identical staining concentrations and imaging conditions.

The superior performance of **ER-Halo** and **ESiR-Halo** in living-cell fluorogenic imaging can be attributed to their enhanced solubility and brightness. Increased water solubility improves dispersibility, reducing non-specific binding to cellular components, minimizing background fluorescence, and enhancing SNR. Furthermore, the higher brightness of ether rhodamine probes enhances imaging quality, making them highly suitable for achieving higher resolution and accuracy in super-resolution imaging. These attributes position **ER-Halo** and **ESiR-Halo** as promising tools for advanced cellular imaging applications.

#### Application in SIM and STED super-resolution imaging

The specific labeling ability and exceptional fluorogenicity of **ER-Halo** and **ESiR-Halo** enabled us to achieve long-term super-resolution imaging, allowing the decoding of dynamic cellular processes. MCF cells expressing Halo-TOMM20 were incubated with **ER-Halo** and **ESiR-Halo** and imaged using SIM (Figure 4a). The resulting images revealed clear fluorescent signals outlining the outer mitochondrial membrane (OMM) structure. A gradual decrease in fluorescence intensity was observed inside the mitochondria, enabling the visualization of the OMM’s separation and determining the width of the mitochondria, approximately 400 nm, from the images.

**Figure 4.**
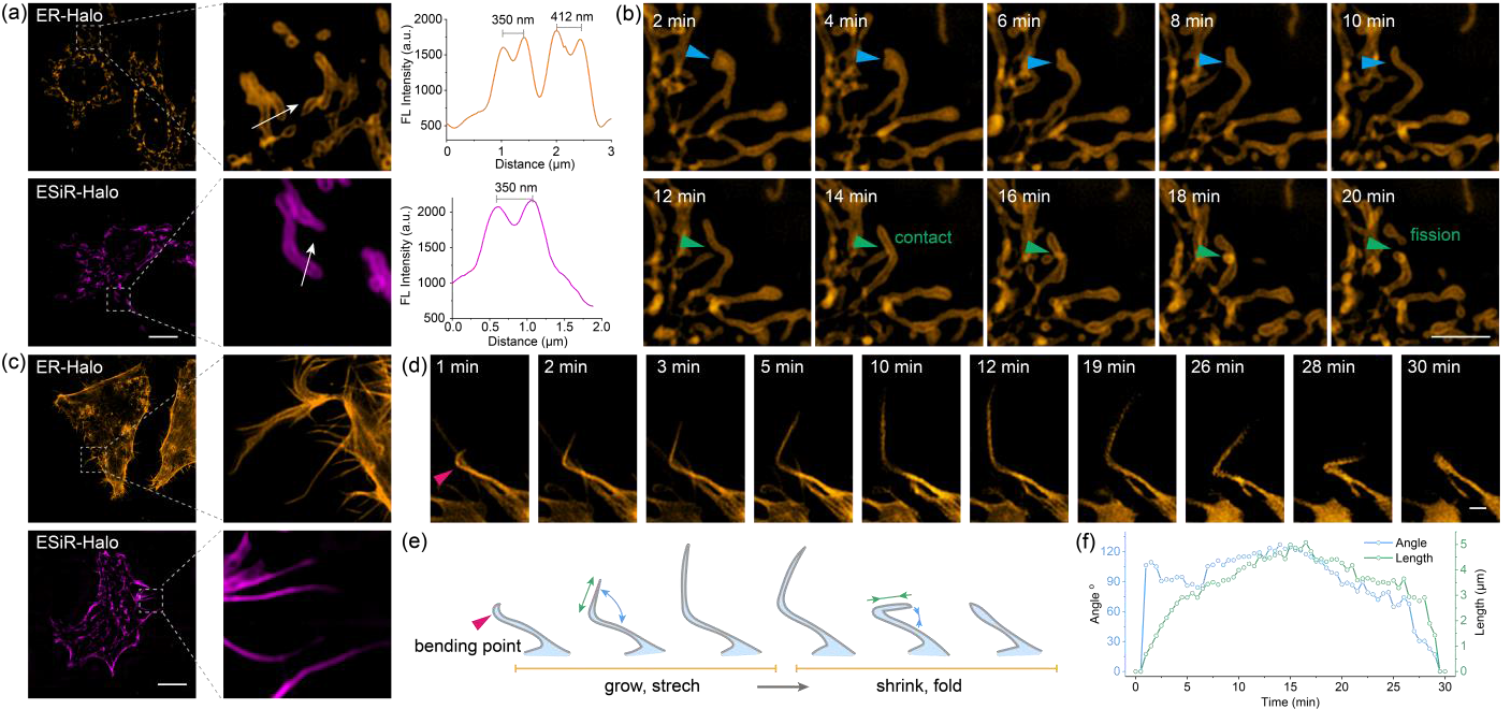
Ether rhodamines-based probes for living-cell SIM imaging. (a) SIM imaging of mitochondria expressing Halo-Tomm20 using **ER-Halo** (top) and **ESiR-Halo** (bottom). Graph: Intensity profile of ROI across cells. (b) Long-term SIM imaging of mitochondria dynamics labeled by **ER-Halo**. (c) SIM imaging of actin expressing Halo-Actin using **ER-Halo** (top) and **ESiR-Halo** (bottom). (d) Long-term SIM imaging of actin growth and shrinkage in 30 minutes. Cells were labeled by **ER-Halo**. (e) Schematic diagram of filopodia growth and shrinking process in (d). (f) Quantitative analysis of the bending angle and the length of filopodia in (d). MCF cells transiently expressing Halo-Tomm20 and Halo-Actin, respectively, were incubated with 100 nM probes for 1 h. Scale bar in (a, c), 10 μm. Scale bar in (b), 5 μm. Scale bar in (d), 1 μm.

We further utilized long-term SIM imaging to track mitochondrial dynamics and observe morphological changes. As shown in Figure 4b, a mitochondrion (blue arrow) initially curved at the apex and gradually stretched into a linear shape over 10 minutes. Additionally, another mitochondrion (green arrow) displayed midsection contact with the apex of a neighboring mitochondrion, which subsequently resulted in fission at the contact site.

To further explore the capabilities of **ER-Halo** and **ESiR-Halo** in super-resolution imaging, MCF cells expressing Halo-Actin were incubated with the probes and subjected to SIM imaging. The results revealed distinct actin structures with high clarity (Figure 4c). Using long-term SIM imaging, we monitored the dynamic process of filopodia growth and retraction in a living cell over 30 minutes. As illustrated in Figure 4d, a rigidly bent filopodium extended gradually from its bending point (magenta arrow), with the bending angle increasing over time. Subsequently, the filopodium began to retract, the bending angle decreased, and eventually, the filopodium disappeared completely. Quantitative analysis of the bending angle and filopodium length (annotated with blue and green arrows, respectively) showed that the bending angle expanded to a maximum of 127° at 14 minutes, while the filopodium reached its most extended length of 5.08 μm at 16.5 minutes. Notably, the maximum angle and length occurred in close temporal proximity. The analysis revealed that filopodium growth correlated with an increase in the bending angle, while retraction coincided with a decrease in the bending angle (Figure 4e-f, Table S10). This correlation suggests that the filopodium growth and shrinkage processes may be synergistic.

We further explored the capabilities of **ER-Halo** and **ESiR-Halo** by transfecting MCF cells with Halo-Tubulin, incubating them with the probes, and visualizing tubulin structures using SIM imaging. The resulting images revealed curly microtubules distributed in a reticular pattern throughout the cells. Fine structural analysis showed clear, filamentous microtubules interlacing with each other, demonstrating the high imaging precision of these probes (Figure S13).

After obtaining excellent SIM imaging results for delicate structures and long-term dynamics of organelles labeled with **ER-Halo** and **ESiR-Halo**, we validated their performance in STED imaging of living cells. First, we examined **ER-Halo** in HeLa cells expressing Halo-Actin. STED imaging provided exceptional resolution, revealing detailed structures not discernible in confocal mode. Specifically, two closely spaced filopodia, indistinguishable in confocal imaging, were resolved under STED imaging. The resolution achieved for the filopodia was 121 nm, with a measured distance of 273 nm between the two filopodia (Figure 5a). We also imaged HeLa cells expressing Halo-TOMM20 and labeled them with **ER-Halo**. The STED images provided a distinct visualization of the outer mitochondrial membrane (OMM), with a more pronounced fluorescence intensity ratio between the OMM and internal mitochondria than SIM imaging (Figure 5b).

**Figure 5.**
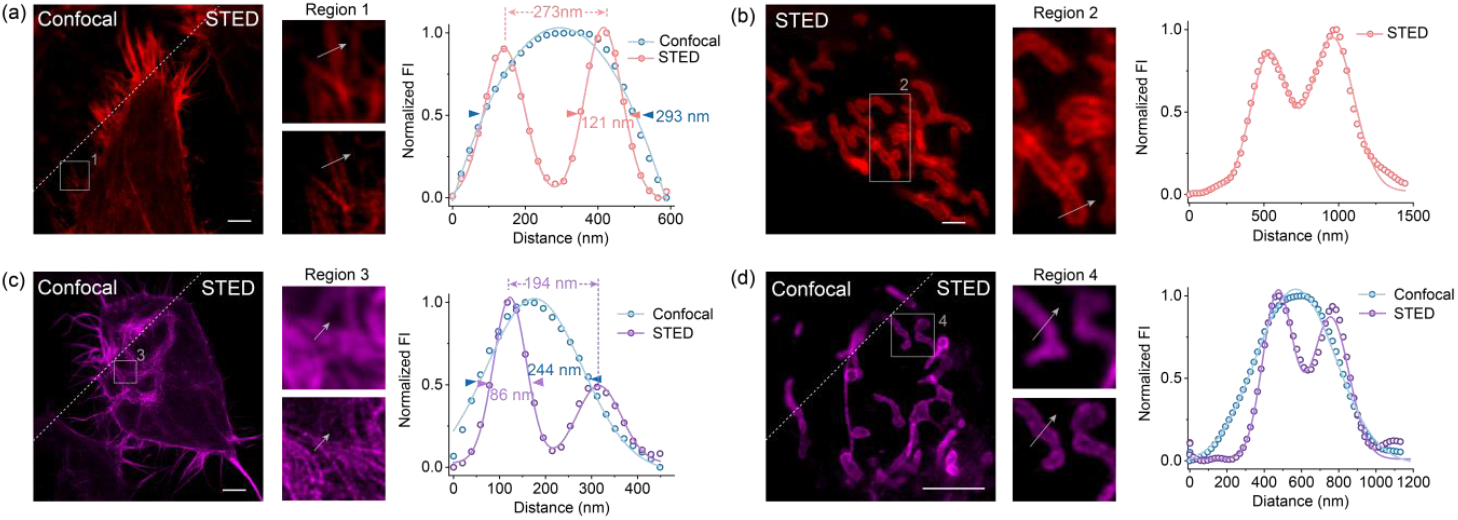
Ether rhodamine-based probes for living-cell STED imaging. (a, b) Confocal and STED imaging of actin (a) and mitochondria using **ER-Halo**. Graph: Intensity profile of ROI across cells. (c, d) Confocal and STED imaging of actin (c) and mitochondria (d) using **ESiR-Halo**. Graph: Intensity profile of ROI across cells. Hela cells transiently expressing Halo-Tomm20 and Halo-Actin, respectively, were incubated with 100 nM probes for 1 h. Scale bar in (a, b, c), 10 μm. Scale bar in (d), 5 μm.

Next, we applied **ESiR-Halo** to STED imaging. Actin bundles, which appeared interwoven and indistinguishable under confocal imaging, were resolved and separated in the STED mode. The resolution for actin structures reached 86 nm, with two actin filaments separated by 194 nm distinctly resolved (Figure 5c). Additionally, **ESiR-Halo** proved effective for OMM imaging under STED, yielding excellent results (Figure 5d).

The strong depletion laser power required for STED microscopy often limits the applicability of many dyes due to photobleaching. However, ether rhodamine probe demonstrated exceptional resilience and performance in STED imaging, attributed to their innate photostability and brightness.

These results confirm the versatility and robustness of **ER-Halo** and **ESiR-Halo** for super-resolution imaging in living cells, making them valuable tools for detailed cellular studies.

#### Application in SMLM imaging

High-brightness fluorophores are essential for SMLM, as they enable precise single-particle tracking with superior spatial and temporal resolution. In addition, fluorophores must exhibit excellent solubility and dispersibility in aqueous environments to ensure effective labeling and accurate localization. Our novel ether rhodamines fulfil these requirements, making them ideal candidates for SIM imaging.

Building on our previous development of a rhodamine-based blinking strategy utilizing intramolecular hydrogen bonding under acidic conditions,^23^ we designed **Lyso-ER**, a lysosome-specific ether rhodamine probe for SMLM imaging (Figure 6a). This probe incorporates an *o*-methylpyridine group to leverage lysosomal acidity for fluorescence activation. **Lyso-ER** remains non-fluorescent at physiological pH due to its spiro-lactam structure. Upon entering lysosomes (pH 4.5–5.5), protons interact with the pyridine nitrogen and carbonyl group, inducing a transition to the fluorescent zwitterion state. This fluorescence activation is reversible, as the zwitterion quickly returns to the spiro-lactam state via a nucleophilic addition reaction between the amide nitrogen and the meso-carbon of the xanthene core. These reversible transitions, mediated by hydrogen bonding, produce blinking fluorescence signals critical for SMLM imaging.

**Figure 6.**
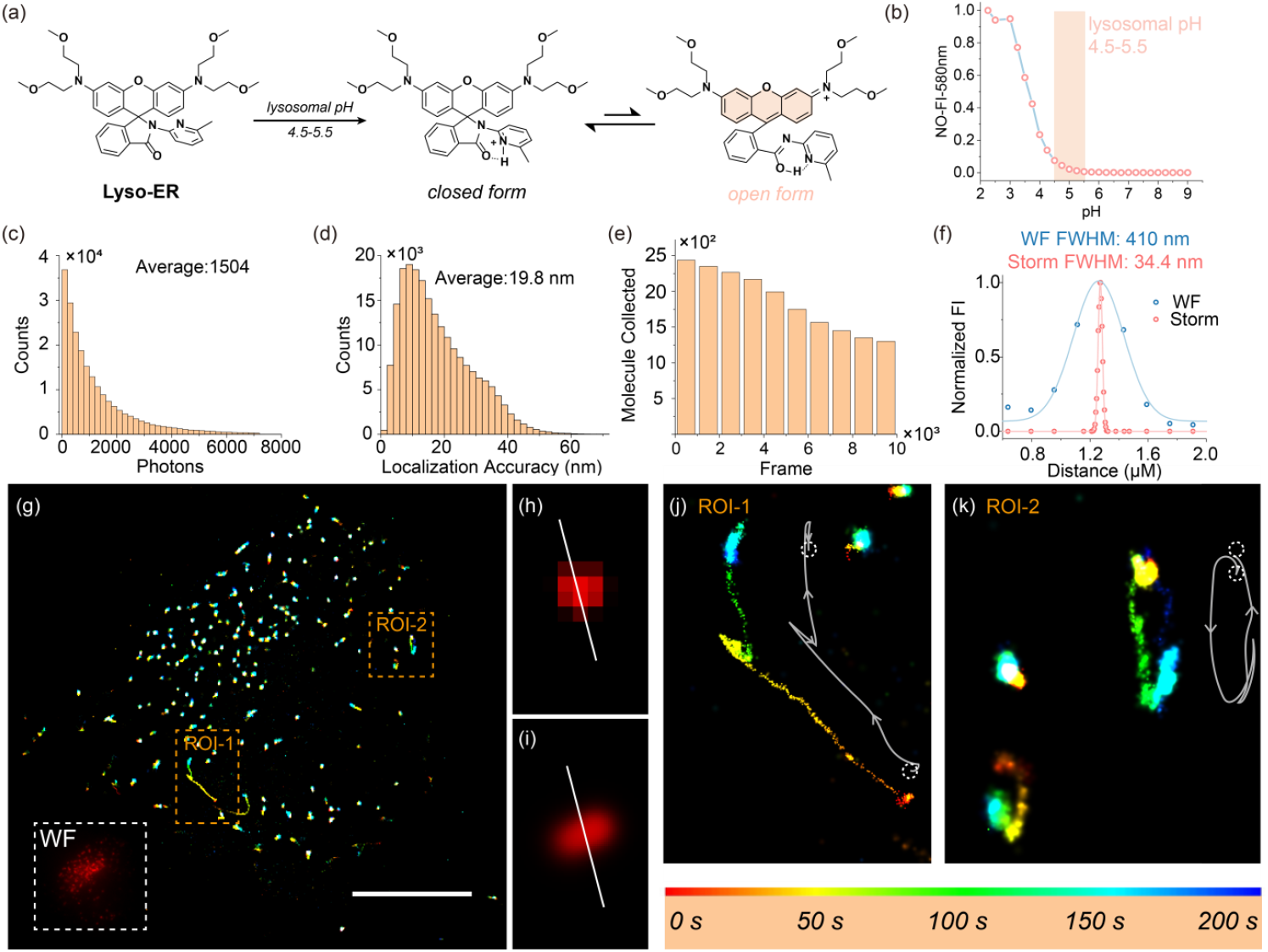
Ether rhodamines-based probe **Lyso-ER** for living-cell lysosome SMLM imaging. (a) Mechanism of **Lyso-ER** spontaneously blinking in lysosomal pH. (b) Fluorescence intensity of **Lyso-ER** at 580 nm as a function of pH in aqueous solution. Histograms of (c) photons per single molecule per frame and (d) localization accuracy. (e) Variation curve of molecules detected during imaging. (f) Intensity distributions across the white lines in (h) and (i). (g) Long-term SMLM imaging of whole-cell lysosomes with **Lyso-ER** in live MCF cells. Inset: Wide-field (WF) imaging. Scale bar: 10 μm. (h, i) WF imaging (h) and SMLM imaging (i) of lysosome in (g). (j, k) Long-term SMLM imaging of the region of interest (ROI) and overlaid time-lapse images of lysosomes.

Compared to traditional rhodamine probes, **Lyso-ER** offers several advantages. Its ether-substituted structure enhances solubility and brightness, improving its ability to label lysosomal targets and provide high photon output. The hydrogen-bond-driven blinking mechanism ensures precise temporal fluorescence control, enabling superior spatial resolution and accurate localization of lysosomal dynamics.

We measured the fluorescence intensity of **Lyso-ER** at 580 nm to determine the proportion of the fluorescent zwitterion state in different pH buffer solutions (Figure 6b, S14, Table S11). At lysosomal pH (4.5–5.5), the proportion of the fluorescent state ranged between 7.5% and 0.6%, while at physiological pH (7.4), it was only 0.3%. This indicates that **Lyso-ER** predominantly exists in the non-fluorescent state at physiological pH, demonstrating its high fluorogenicity for lysosome labeling.

Additionally, cell viability assays revealed that more than 90% of cells remained viable after 24 hours of incubation at the imaging concentration (Figure S15), confirming the biocompatibility of **Lyso-ER** for live-cell imaging applications.

We utilized **Lyso-ER** for lysosomal SMLM imaging in living cells, tracking lysosomal dynamics across 200 seconds with 10,000 frames (Figure 6c-k). From the SMLM images, we extracted quantitative photophysical and localization data. **Lyso-ER** emitted an average of 1,504 photons per molecule (Figure 6c), achieving an impressive localization precision of 19.8 nm on average (Figure 6d). Additionally, more than 50% of the molecules maintained blinking activity throughout the 10,000 frames (Figure 6e), ensuring consistent tracking and signal quality.

Resolution comparison between widefield (WF) and SMLM imaging underscored **Lyso-ER**’s exceptional performance in super-resolution imaging. For the same lysosome, the width measured using SMLM was 34.4 nm, compared to 410 nm in the WF image, reflecting a resolution improvement over an order of magnitude (Figure 6f, h-i).

The ether modification of **Lyso-ER** enhances its solubility and brightness in aqueous environments, making it especially effective in the acidic lysosomal interior. Leveraging these properties, we tracked and analyzed lysosomal dynamics (Figure 6j-k). In ROI-1, a lysosome displayed two distinct instances of oscillatory motion: it advanced a short distance, retreated slightly, and repeated the cycle before eventually progressing further. In ROI-2, two lysosomes exhibited circular trajectories, with one lysosome also showing a brief segment of oscillatory movement. These results highlight the practicality and efficacy of **Lyso-ER** for SMLM imaging in aqueous environments.

## CONCLUSION

We developed a new class of ether-substituted rhodamines by attaching ether chains to the amino groups, significantly improving water solubility and fluorescence brightness. We elucidated the mechanisms underlying these enhancements through a combination of experimental studies and quantum chemical calculations. Specifically, ether chains promote effective dispersion of rhodamines in aqueous environments, reducing aggregation-caused quenching and minimizing intracellular non-specific background fluorescence. Additionally, the ether chains lower the electron-donating capacity and the driving energy for TICT formation. Their flexibility further allows them to form a protective shield around the rhodamine core, suppressing TICT and external conversion, which collectively enhance the brightness of the dyes. Building on these advances, we developed fluorogenic probes **ER-Halo** and **ESiR-Halo** for HaloTag labeling and imaging and a lysosome-specific SMLM probe for tracking dynamic intracellular processes. Compared to traditional N-alkyl rhodamine probes, ether rhodamine probes exhibited superior specificity and brightness in live-cell imaging. Their performance enabled long-term monitoring of organelle dynamics, providing detailed insights into complex cellular processes with high spatial and temporal resolution. These findings demonstrate the exceptional potential of ether rhodamines as advanced fluorophores for live-cell nanoscopy, offering a versatile and practical solution for high-resolution imaging and dynamic tracking in aqueous environments.

## Supporting information

Materials, synthesis and characterization, computational calculations, photophysicaltables and pictures, imaging pictures.

## ASSOCIATED CONTENT

### Supporting Information

Additional materials and instructments, experimental details, synthesis and characterization analysis, computational calculations, photophysical properties tables and pictures, imaging pictures. (file type, PDF)

## AUTHOR INFORMATION

### Author

**Xiangning Fang** - Dalian Institute of Chemical Physics, Chinese Academy of Sciences, Dalian 116023, China; University of Chinese Academy of Sciences, Beijing 100049, China

**Zhifeng Li** - Dalian Institute of Chemical Physics, Chinese Academy of Sciences, Dalian 116023, China

**Hao-Kai Li -** Fluorescence Research Group, Singapore University of Technology and Design, 8 Somapah Road, 487372, Singapore

**Jie Chen** - Dalian Institute of Chemical Physics, Chinese Academy of Sciences, Dalian 116023, China; University of Chinese Academy of Sciences, Beijing 100049, China

**Ning Xu** - Dalian Institute of Chemical Physics, Chinese Academy of Sciences, Dalian 116023, China

**Kai An** - Dalian Institute of Chemical Physics, Chinese Academy of Sciences, Dalian 116023, China; University of Chinese Academy of Sciences, Beijing 100049, China

**Wenchao Jiang** - Dalian Institute of Chemical Physics, Chinese Academy of Sciences, Dalian 116023, China; University of Chinese Academy of Sciences, Beijing 100049, China

**Yi Tao** - Dalian Institute of Chemical Physics, Chinese Academy of Sciences, Dalian 116023, China; University of Chinese Academy of Sciences, Beijing 100049, China

**Pengjun Bao** - Dalian Institute of Chemical Physics, Chinese Academy of Sciences, Dalian 116023, China; University of Chinese Academy of Sciences, Beijing 100049, China

**Zhimin Wu** - Fluorescence Research Group, Singapore University of Technology and Design, 8 Somapah Road, 487372, Singapore

**Yinchan Zhang** - Dalian Institute of Chemical Physics, Chinese Academy of Sciences, Dalian 116023, China; University of Chinese Academy of Sciences, Beijing 100049, China

### Author Contributions

The manuscript was written through the contributions of all authors. All authors have given approval to the final version of the manuscript.

### Notes

The authors declare no competing financial interest.

## ACKNOWLEDGMENT

This work is supported by the National Natural Science Foundation of China (22225806, 22078314, 22278394, 22378385), Dalian Institute of Chemical Physics (DICPI202142, DICPI202436), the Ministry of Education, Singapore (MOE-T2EP10222-0001) and the Singapore University of Technology and Design (SUTD) Kickstarter Initiative (No. SKI 2021_04_09). The authors are grateful for the computing service of SUTD and the National Supercomputing Centre (Singapore).

