## Supplementary material for "Ether Rhodamines with Enhanced Hydrophilicity, Fluorogenicity, and Brightness for Super-Resolution Imaging": Materials, synthesis and characterization, computational calculations, photophysicaltables and pictures, imaging pictures.

### Materials and Instruments

#### Materials

Unless explicitly stated, all reagents were purchased from commercial suppliers (Sigma-Aldrich, *J&K*, Innochem, and Aladdin) and used without further purification. Solvents [dimethyl sulfoxide (DMSO), ethanol (EtOH), and acetonitrile ($CH_3CN$)] were purchased from *J&K* and used without further treatment or distillation. Silica gel (200-300 mesh) was purchased from Innochem.

#### Instruments

$^1$H-NMR spectra were recorded on a Bruker 400 spectrometer and Avance III HD 700 MHz, using TMS as an internal standard. Chemical shifts were given in ppm and coupling constants (J) in Hz.
For UV-vis absorption spectra were collected on an Agilent Cary 60 UV-Vis Spectrophotometer. Fluorescence measurements were performed on an Agilent CARY Eclipse fluorescence spectrophotometer.
Confocal images were performed on Andor Revolution WD, a 100× / NA 1.49 oil objective lens, LU-NV series laser unit (laser combination: 488 nm; 561 nm; 640 nm).
Structured illumination microscopy (SIM) super-resolution images and single-molecule localization super-resolution microscopy (SMLM) images were obtained with Nikon N-SIM/STORM 5.0 Super-Resolution Microscope System with a motorized inverted microscope ECLIPSE Ti2-E, a 100 × / NA 1.49 oil immersion TIRF objective lens (CFI HP), LU-NV series laser unit (405 nm, 488 nm, 561 nm, 647 nm), and an ORCA-Flash 4.0 SCMOS camera (Hamamatsu Photonics K.K.).
Stimulated emission depletion microscopy (STED) images were implemented on Leica TCS SP8 gSTED 3X. The appropriate, 592 and 660 nm, depletion laser was used for **ER-Halo** and **ESiR-Halo** respectively. Fluorescence channels were scanned sequentially and emission was revealed by means of hybrid spectral detectors (HyD SP Leica Microsystems).

### Experimental Procedures

#### Spectral measurements in the different solvents

The probe stock solution (2 mM) was prepared in DMSO. Samples for spectroscopic analysis were prepared by diluting the stock solution into different solvents. The concentration of samples, e.g., **ER1**, **ER2**, **ER1-Ester**, **ER2-Ester**, **ESiR**, **ESiR-Me**, was usually prepared at 2 μM, unless stated otherwise. TFA (0.1% v/v) was added to the sample solutions to test the completely open-formed rhodamines.
**RhoB** was used to obtain relative fluorescence quantum yields of O-bridge rhodamines (fluorescence quantum yields of **RhoB** is 0.35 in PBS)[1]. **TMSiR** was used to obtain relative fluorescence quantum yields of Si-bridge rhodamines (fluorescence quantum yields of **TMSiR** is 0.41 in PBS)[1]. The quantum yield ($\varphi$) was calculated using the following equation.

$$\varphi F_{(X)} = \varphi F_{(S)} \cdot (A_S F_X / A_X F_S)(\eta_X / \eta_S)^2$$

where $\varphi$F, A, and F represent the fluorescence quantum yield, the absorbance at the excitation wavelength, and the area under the corrected emission curve, respectively. And η is the refractive index of the solvent. Subscripts X and S refer to the unknown and the standard.

#### Cell culture

Hela (helacyton gartleri) and MCF cells were purchased from the Cell Bank of Type Culture

Collection of the Chinese Academy of Sciences. HeLa cells were maintained in Dulbecco's modified Eagle's medium (DMEM, Gibco) supplemented with 10% fetal bovine serum (FBS, Hyclone), which were cultured in a humidified atmosphere of 5% $CO_2$/95% air at 37°C. Before the imaging experiments, cells were seeded on a glass bottom cell culture dish (Nest, polystyrene, Φ 15 mm) for 1-2 days to reach 60-80% confluency. The cells were then used for further experiments.

#### Cell transfection

Transfection experiments were performed with Lipofectamine 2000 (Invitrogen) according to the manufacturer's protocol.

Briefly, 2 μL Lipofectamine 2000 (Invitrogen) and appropriate plasmid were firstly diluted in 20 μL DMEM (Dulbecco's modified eagle medium), respectively. 5 min later, the diluted plasmid in 20 μL DMEM was added to the diluted Lipofectamine 2000 (Invitrogen) with homogeneous mixing. Another 10 min later, the mixture was added to the cell-culture dish in 1 mL DMEM. The final concentration of plasmid was controlled at 1000 ng/mL. After incubation for 4 h at 37 °C, the culture medium was changed from DMEM to DMEM with 10% FBS. 24-48 h later, the transfected cells were used for super-resolution imaging.

#### Cytotoxicity

The cytotoxicity of **ER-Halo**, **ESiR-Halo**, and **Lyso-ER** was then inspected using MTT assays. MCF Cells were seeded into 96-well plates with $1 \times 10^4$ cells/well and cultured for 24 h. After being treated with various concentrations (0, 0.1, 0.5, 1.0, 2.0, 5.0, 7.5, and 10 μM) of ER-Halo, ESiR-Halo, and Lyso-ER, respectively, the cells were incubated for 24 h, at 37 °C with a 5% $CO_2$ atmosphere. Then, the medium was replaced with 200 μL fresh medium and 20 μL of 5 mg/mL MTT solution per well. 4 h later, the medium was removed carefully, and 200 μL DMSO was added to each well to dissolve blue formazan. After sufficient shaking for 15 min, the absorption was recorded at 490 nm using a UV-Vis microplate reader and used for analysis of cell viability.

The results in Figure S10 and Figure S13 showed that >90% of HeLa cells survived after 24 h at imaging concentration, demonstrating the low toxicity of these probes toward cultured cell lines.

#### Imaging

##### Confocal imaging of living cells

For imaging of living cells, the experimental group was incubated with probes in DMEM for 60 min at 37°C, then directly used for imaging. Microscopic images of the cells were obtained on a live-cell confocal laser scanning microscope without wash-out steps.

##### SIM imaging of living cells

For imaging of living cells by SIM, the experimental group was incubated with probes in DMEM for 60 min at 37°C, then directly used for imaging.

##### SMLM imaging of living cells

For imaging of living cells by SMLM, the experimental group was incubated with probes (2 μM) in DMEM for 60 min at 37°C, then directly used for imaging.

##### STED imaging of living cells

For imaging of living cells by STED, the experimental group was incubated with probes in DMEM for 60 min at 37°C, then directly used for imaging.

#### Computational Methods

Quantum calculations were performed utilizing Gaussian 16A.[2] All DFT/TD-DFT calculations were performed at the $\omega$B97XD/def2-SVP[3-4] level of theory either in vacuo or in water. The solvent effects (in water) were accounted for using the SMD solvation model.[5] All the structures were optimized and validated by checking positive vibrational frequencies.

The conformational search of **ER1** was done using the CREST software.[6] We subsequently conducted DFT calculations to optimize their molecular structures and determine the corresponding energy values by utilizing a range of conformations generated by CREST.

To calculate the ionization potential, we optimized the molecular structures of both the substituents before and after the loss of an electron. The differences in their Gibbs free energy provide us with the ionization potential.

Avogadro software was used to visualize molecules and their frontier molecular orbitals.[7] When constructing the $S_1$ potential energy surface following the TICT process, we systematically varied two dihedral angles around an amino group from 0° to 90° at 10° intervals. During the subsequent optimization, the dihedral angles for each step were fixed. At the same time, all other parameters were freely optimized using the B3LYP/def2SVP level of theory in a water environment (utilizing the SMD model).

Following this optimization, based on the optimized structures in the $S_1$ state, we computed their respective electronic energies using the CAM-B3LYP/def2-SVP level of theory in water, incorporating the corrected state-specific solvation formalism.

#### Supplementary tables

Table S1. Photophysical properties of **ER1**.

| **Solvent** | $\lambda_{abs}$ (nm) | $\lambda_{em}$ (nm) | $\Delta\lambda$ (nm) | $\varepsilon$ ($M^{-1}$ $cm^{-1}$) | $\Phi$ |
|---|---|---|---|---|---|
| $CHCl_3$ | 550 | 570 | 20 | -- | -- |
| +TFA | 551 | 572 | 21 | 80657 | 0.98 |
| Dioxane | 550 | 563 | 13 | -- | -- |
| +TFA | 558 | 583 | 25 | 22441 | 0.85 |
| EA | 550 | 585 | 24 | -- | -- |
| +TFA | 554 | 578 | 24 | 74559 | 0.90 |
| ACN | 554 | 578 | 24 | -- | -- |
| +TFA | 554 | 579 | 25 | 80890 | 0.63 |
| DMSO | 565 | 57 | 22 | -- | -- |
| +TFA | 563 | 590 | 27 | 75795 | 0.89 |
| EtOH | 542 | 566 | 24 | 45274 | 0.96 |
| +TFA | 543 | 576 | 33 | 81032 | 0.84 |
| MeOH | 543 | 568 | 25 | 72646 | 0.86 |
| +TFA | 550 | 576 | 26 | 80026 | 0.86 |
| $H_2O$ | 545 | 570 | 25 | 69265 | 0.61 |
| +TFA | 548 | 575 | 27 | 71066 | 0.51 |
| Glycol | 552 | 575 | 23 | 57785 | 0.95 |
| +TFA | 558 | 579 | 21 | 77305 | 1.00 |

Table S2. Photophysical properties of **ER2**.

| **Solvent** | $\lambda_{abs}$ (nm) | $\lambda_{em}$ (nm) | $\Delta\lambda$ (nm) | $\varepsilon$ ($M^{-1}$ $cm^{-1}$) | $\Phi$ |
|---|---|---|---|---|---|
| $CHCl_3$ | 555 | 570 | 25 | -- | -- |
| +TFA | 553 | 574 | 21 | 89125 | 0.98 |
| Dioxane | 562 | 580 | 18 | -- | -- |
| +TFA | 559 | 584 | 25 | 33710 | 0.88 |
| EA | 563 | 584 | 21 | -- | -- |
| +TFA | 556 | 579 | 23 | 77475 | 0.84 |
| ACN | 546 | 573 | 27 | -- | -- |
| +TFA | 555 | 580 | 25 | 83320 | 0.64 |
| DMSO | 564 | 588 | 24 | -- | -- |
| +TFA | 564 | 590 | 26 | 72495 | 0.82 |
| EtOH | 542 | 567 | 25 | 41980 | 0.99 |
| +TFA | 554 | 578 | 24 | 80125 | 0.89 |
| MeOH | 544 | 569 | 25 | 71735 | 0.84 |
| +TFA | 553 | 577 | 24 | 84020 | 0.72 |
| $H_2O$ | 545 | 571 | 26 | 65770 | 0.70 |
| +TFA | 550 | 576 | 26 | 69135 | 0.62 |
| Glycol | 552 | 575 | 23 | 87975 | 0.91 |
| +TFA | 559 | 580 | 21 | 83285 | 0.95 |

Table S3. Photophysical properties of **ESiR**.

| Solvent | $\lambda_{abs}$ (nm) | $\lambda_{em}$ (nm) | $\Delta\lambda$ (nm) | $\varepsilon$ ($M^{-1}$ $cm^{-1}$) | $\Phi$ |
|---|---|---|---|---|---|
| $CHCl_3$ | -- | -- | -- | -- | -- |
| +TFA | 653 | 669 | 16 | 39350 | 0.52 |
| Dioxane | -- | -- | -- | -- | -- |
| +TFA | -- | -- | -- | -- | -- |
| EA | -- | -- | -- | -- | -- |
| +TFA | 656 | 671 | 15 | 6540 | 0.72 |
| ACN | -- | -- | -- | -- | -- |
| +TFA | 653 | 672 | 19 | 143975 | 0.68 |
| DMSO | -- | -- | -- | -- | -- |
| +TFA | 664 | 682 | 18 | 17290 | 0.75 |
| EtOH | -- | -- | -- | -- | -- |
| +TFA | 653 | 671 | 18 | 153155 | 0.54 |
| MeOH | -- | -- | -- | -- | -- |
| +TFA | 651 | 672 | 21 | 153970 | 0.50 |
| $H_2O$ | 643 | 664 | 21 | 28055 | 0.41 |
| +TFA | 649 | 669 | 20 | 57225 | 0.34 |

Table S4. Photophysical properties of **ER1-Ester**.

| Solvent | $\lambda_{abs}$ (nm) | $\lambda_{em}$ (nm) | $\Delta\lambda$ (nm) | $\varepsilon$ ($M^{-1}$ $cm^{-1}$) | $\Phi$ |
|---|---|---|---|---|---|
| $CHCl_3$ | 556 | 576 | 20 | 81851 | 0.93 |
| Dioxane | 552 | 588 | 26 | 43486 | 0.46 |
| EA | 568 | 581 | 13 | 74204 | 0.59 |
| ACN | 555 | 580 | 25 | 57962 | 0.61 |
| DMSO | 565 | 591 | 26 | 72036 | 0.84 |
| EtOH | 556 | 579 | 23 | 84578 | 0.84 |
| MeOH | 555 | 577 | 22 | 78626 | 0.68 |
| $H_2O$ | 550 | 577 | 27 | 72413 | 0.50 |
| Glycol | 561 | 582 | 21 | 79450 | 0.92 |

Table S5. Photophysical properties of **ER2-Ester**.

| Solvent | $\lambda_{abs}$ (nm) | $\lambda_{em}$ (nm) | $\Delta\lambda$ (nm) | $\varepsilon$ ($M^{-1}$ $cm^{-1}$) | $\Phi$ |
|---|---|---|---|---|---|
| $CHCl_3$ | 555 | 574 | 19 | 82575 | 0.99 |
| Dioxane | 562 | 586 | 24 | 59270 | 0.69 |
| EA | 556 | 580 | 24 | 73055 | 0.73 |
| ACN | 555 | 580 | 25 | 75745 | 0.51 |
| DMSO | 562 | 590 | 28 | 73655 | 0.78 |
| EtOH | 554 | 579 | 25 | 80400 | 0.73 |
| MeOH | 553 | 578 | 25 | 78405 | 0.65 |
| $H_2O$ | 551 | 577 | 26 | 74565 | 0.56 |
| Glycol | 557 | 576 | 21 | 72040 | 0.96 |

Table S6. Photophysical properties of **ESiR-Me**.

| **Solvent** | $\lambda_{abs}$ (nm) | $\lambda_{em}$ (nm) | $\Delta\lambda$ (nm) | $\varepsilon$ ($M^{-1}$ $cm^{-1}$) | $\Phi$ |
|---|---|---|---|---|---|
| $CHCl_3$ | 655 | 667 | 12 | 135555 | 0.57 |
| Dioxane | 655 | 669 | 14 | 60790 | 0.33 |
| EA | 656 | 671 | 15 | 117850 | 0.37 |
| ACN | 655 | 670 | 15 | 124790 | 0.51 |
| DMSO | 666 | 683 | 17 | 121675 | 0.55 |
| EtOH | 656 | 671 | 15 | 138560 | 0.52 |
| MeOH | 655 | 670 | 15 | 134435 | 0.47 |
| $H_2O$ | 648 | 665 | 17 | 112410 | 0.33 |

Table S7. Quantum yield of open-formed rhodamines at different concentrations in PBS (10 mM).

| Concentration (μM) | **ER1-Ester** | **ER2-Ester** | **ESiR-Me** | **RhoB-Ester** | **JF549-Ester** | **TMSiR-Me** | **JF646-Me** |
|---|---|---|---|---|---|---|---|
| 1 | 0.5174 | 0.5561 | 0.3179 | 0.1729 | 0.8945 | 0.3680 | 0.4700 |
| 2 | 0.4900 | 0.5458 | 0.3300 | 0.1655 | 0.8769 | 0.3319 | 0.41004 |
| 5 | 0.4420 | 0.4601 | 0.2892 | 0.1368 | 0.6899 | 0.2654 | 0.32933 |
| 10 | 0.3746 | 0.3713 | 0.2184 | 0.1132 | 0.5780 | 0.2000 | 0.23225 |

Table S8. Photophysical properties of **ER-Halo** and **ESiR-Halo**.

| | **ER-Halo** | | | | | **ESiR-Halo** | | | | |
|---|---|---|---|---|---|---|---|---|---|---|
| **Solvent** | $\lambda_{abs}$ (nm) | $\lambda_{em}$ (nm) | $\Delta\lambda$ (nm) | $\varepsilon$ ($M^{-1}$ $cm^{-1}$) | $\Phi$ | $\lambda_{abs}$ (nm) | $\lambda_{em}$ (nm) | $\Delta\lambda$ (nm) | $\varepsilon$ ($M^{-1}$ $cm^{-1}$) | $\Phi$ |
| ACN | -- | -- | -- | -- | -- | -- | -- | -- | -- | -- |
| +TFA | 556 | 582 | 26 | 99800 | 0.59 | 657 | 676 | 19 | 128775 | 0.34 |
| DMSO | -- | -- | -- | -- | -- | -- | -- | -- | -- | -- |
| +TFA | 565 | 592 | 27 | 93220 | 0.80 | 657 | 672 | 15 | 96545 | 0.46 |
| $CHCl_3$ | -- | -- | -- | -- | -- | -- | -- | -- | -- | -- |
| +TFA | 555 | 577 | 22 | 92240 | 1.00 | 666 | 684 | 18 | 31540 | 0.41 |
| EA | -- | -- | -- | -- | -- | -- | -- | -- | -- | -- |
| +TFA | 556 | 580 | 24 | 96130 | 0.82 | 660 | 675 | 15 | 6955 | 0.61 |
| EtOH | 545 | 570 | 25 | 71670 | 0.99 | -- | -- | -- | -- | -- |
| +TFA | 557 | 579 | 22 | 98190 | 0.83 | 656 | 673 | 17 | 127895 | 0.37 |
| $H_2O$ | 551 | 577 | 26 | 69910 | 0.58 | 649 | 669 | 20 | 27235 | 0.33 |
| +TFA | 554 | 581 | 27 | 65530 | 0.52 | 654 | 672 | 18 | 127895 | 0.25 |
| MeOH | 547 | 574 | 27 | 91090 | 0.80 | -- | -- | -- | -- | -- |
| +TFA | 555 | 582 | 27 | 103810 | 0.68 | 656 | 674 | 18 | 128330 | 0.49 |
| PBS | 551 | 575 | 24 | 69010 | 0.51 | 649 | 669 | 20 | 24565 | 0.27 |

Table S9. Plasmids for HaloTag expression in different organelles.

| Name | Addgene# | Location |
|---|---|---|
| LZ10 PBREBAC-H2BHalo | 91564 | Histone H2B in chromatin |
| pSEMS-Tom20-Halo7Tag | 111135 | Outer mitochondrial membrane Tomm20 |
| LifeAct-HaloTag | 176105 | F-actin |
| TUBB5-Halo | 64691 | Beta-tubulin |

Table S10. The bending angle and length of newly grown filopodia in long-term SIM imaging.

| | | | | | | | | | | | |
|---|---|---|---|---|---|---|---|---|---|---|---|
| Time (min) | 0 | 0.5 | 1 | 1.5 | 2 | 2.5 | 3 | 3.5 | 4 | 4.5 | 5 |
| Bending angle (°) | 0 | 0 | 106.61 | 109.51 | 105.06 | 90.71 | 92.25 | 91.57 | 94.44 | 94.66 | 86.01 |
| Length (μm) | 0 | 0 | 0.68 | 1 | 1.41 | 1.8 | 2.11 | 2.41 | 2.68 | 2.92 | 2.91 |
| Time (min) | 5.5 | 6 | 6.5 | 7 | 7.5 | 8 | 8.5 | 9 | 9.5 | 10 | 10.5 |
| Bending angle (°) | 87.57 | 84.2 | 85.85 | 105.51 | 106.88 | 110.96 | 106.75 | 111.5 | 111.46 | 114.72 | 115.14 |
| Length (μm) | 3.08 | 2.91 | 3.26 | 3.44 | 3.51 | 3.42 | 3.58 | 3.61 | 3.71 | 3.99 | 4.09 |
| Time (min) | 11 | 11.5 | 12 | 12.5 | 13 | 13.5 | 14 | 14.5 | 15 | 15.5 | 16 |
| Bending angle (°) | 115.17 | 118.26 | 119.22 | 113.46 | 123.66 | 120.8 | 127.12 | 123.03 | 124.73 | 121.98 | 121.35 |
| Length (μm) | 4.24 | 4.15 | 4.53 | 4.41 | 4.47 | 4.58 | 4.67 | 4.43 | 4.99 | 4.87 | 4.81 |
| Time (min) | 16.5 | 17 | 17.5 | 18 | 18 | 18.5 | 19 | 19.5 | 20 | 20.5 | 21 |
| Bending angle (°) | 117.51 | 114.91 | 110.3 | 109.69 | 109.69 | 103.79 | 99.15 | 98.01 | 87.23 | 88.72 | 86.72 |
| Length (μm) | 5.08 | 4.73 | 4.4 | 4.43 | 4.43 | 4.32 | 4.41 | 4.41 | 4.27 | 4 | 4.18 |
| Time (min) | 21.5 | 22 | 22.5 | 23 | 23.5 | 24 | 24.5 | 25 | 25.5 | 26 | 26.5 |
| Bending angle (°) | 80.39 | 90.13 | 79.25 | 77.35 | 72.03 | 78.22 | 79.41 | 64.92 | 66.68 | 74.18 | 67.73 |
| Length (μm) | 3.59 | 3.66 | 3.67 | 3.61 | 3.48 | 3.52 | 3.47 | 3.58 | 3.29 | 3.65 | 2.94 |
| Time (min) | 27 | 27.5 | 28 | 28.5 | 29 | 29.5 | 30 | | | | |
| Bending angle (°) | 40.48 | 30.66 | 30.62 | 22.92 | 17.02 | 0 | 0 | | | | |
| Length (μm) | 2.91 | 2.78 | 2.91 | 1.89 | 1.43 | 0 | 0 | | | | |

Table S11. The percent of the "on" state of **Lyso-ER** at different pH.

| pH | 2.25 | 2.5 | 3 | 3.25 | 3.5 | 3.75 | 4 | 4.25 | 4.5 | 4.75 | 5 |
|---|---|---|---|---|---|---|---|---|---|---|---|
| FI Intensity | 713.496 | 670.913 | 676.660 | 550.579 | 418.355 | 302.659 | 167.135 | 98.8707 | 53.8358 | 32.1307 | 15.3146 |
| percent of "on" state | 1 | 0.9403 | 0.9484 | 0.7717 | 0.5863 | 0.4242 | 0.2342 | 0.1386 | 0.0755 | 0.045 | 0.0215 |

| pH | 5 | 5.25 | 5.5 | 5.75 | 6 | 6.25 | 6.5 | 6.75 | 7 | 7.25 | 7.4 |
|---|---|---|---|---|---|---|---|---|---|---|---|
| FI Intensity | 15.3146 | 8.7560 | 4.7037 | 3.0470 | 2.2819 | 0.9554 | 1.0446 | 0.0613 | 1.2655 | 0.6773 | 0.2150 |
| percent of "on" state | 0.0215 | 0.0123 | 0.0066 | 0.0043 | 0.0032 | 0.0013 | 0.0015 | 0.0001 | 0.0018 | 0.0009 | 0.0003 |

### Supplementary figures

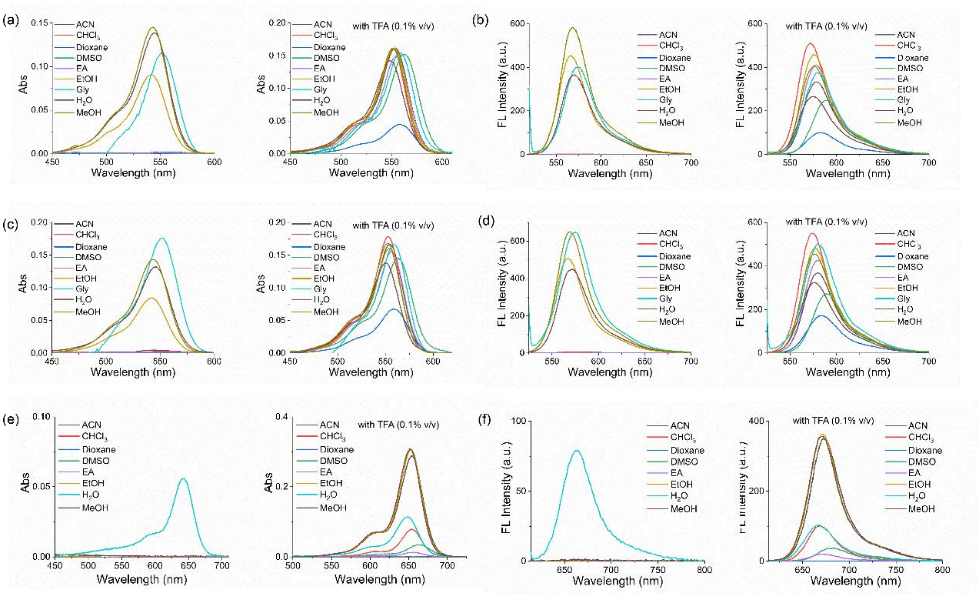

**Figure S1**. (a,b) UV-Vis absorption (a) and fluorescence spectra (b) of **ER1** in different solvents. (c, d) UV-Vis absorption (c) and fluorescence spectra (d) of **ER2** in different solvents. (e, f) UV-Vis absorption (e) and fluorescence spectra (f) of **ESiR** in different solvents. The concentration of the sample is 2 μM.

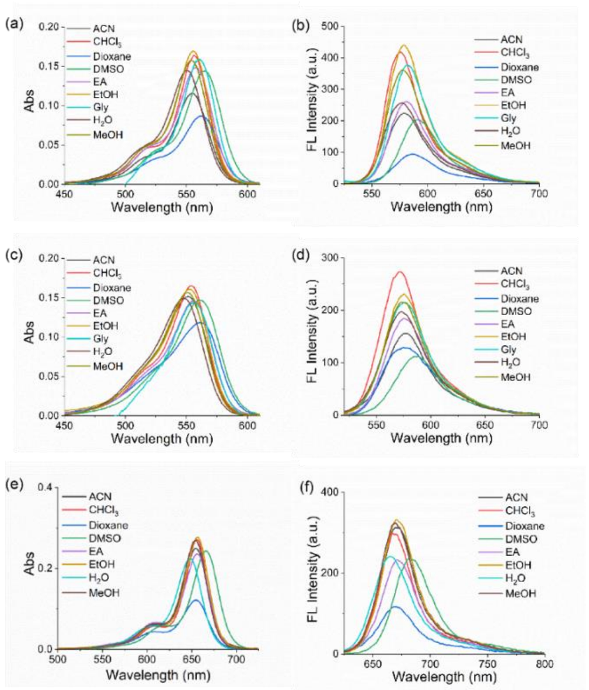

**Figure S2**. (a, b) UV-Vis absorption (a) and fluorescence spectra (b) of **ER1-Ester** in different solvents. (c, d) UV-Vis absorption (c) and fluorescence spectra (d) of **ER2-Ester** in different solvents. (e, f) UV-Vis absorption (e) and fluorescence spectra (f) of **ESiR-Me** in different solvents. The concentration of the sample is 2 μM.

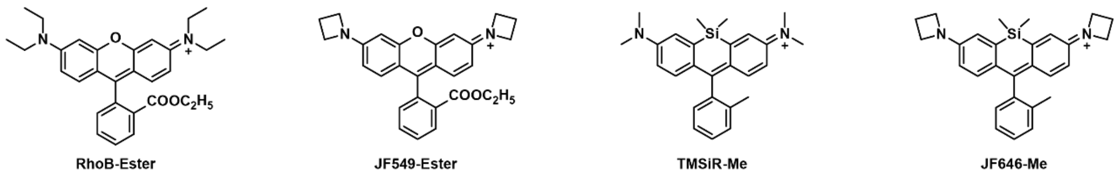

**Figure S3**. Structure of open-formed N-alkyl and azetidine rhodamines

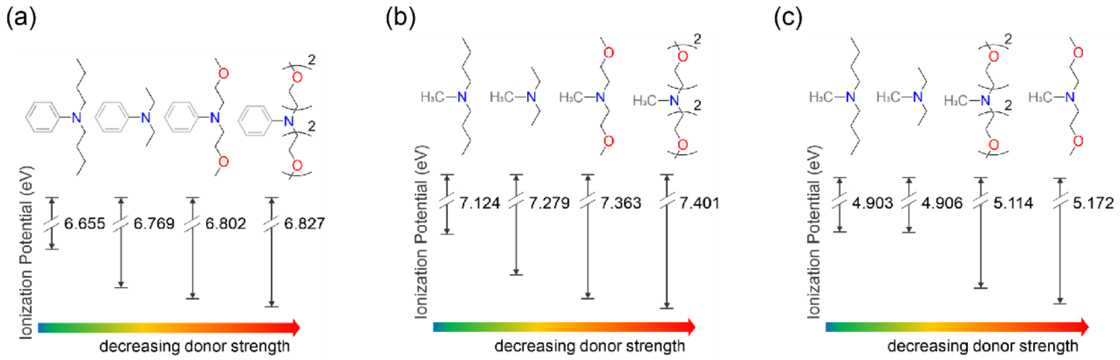

**Figure S4**. (a) Calculated ionization potential of various amino groups (passivated with a phenyl ring) in vacuo. (b) Calculated ionization potential of various amino groups (passivated with a methyl group) in vacuo. (c) Calculated ionization potential of various amino groups (passivated with a methyl group) in water.

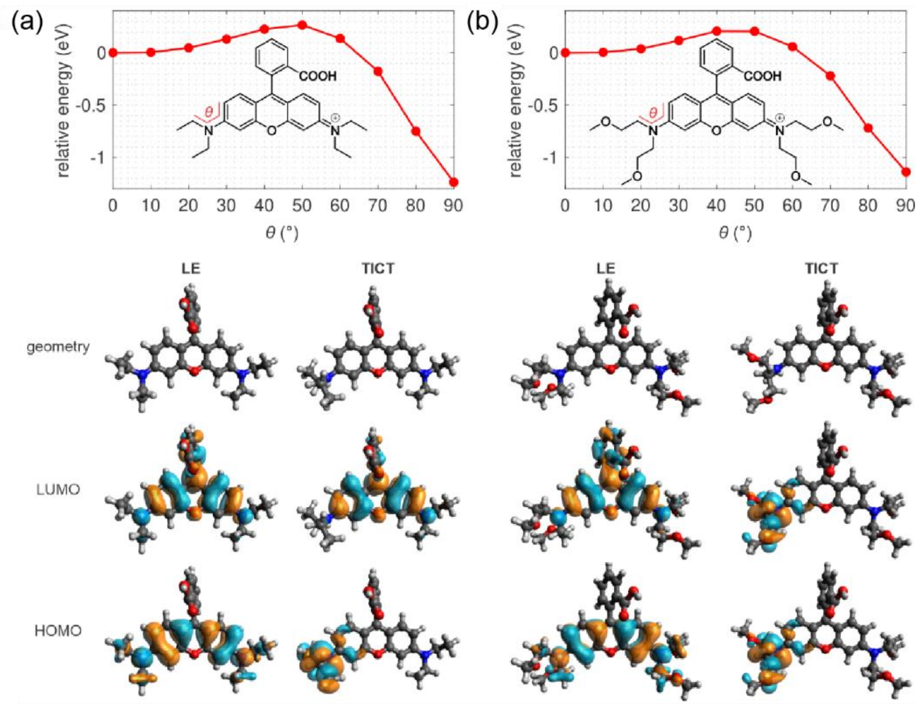

**Figure S5.** Relative electronic energy of the $S_1$ potential energy surface of (a) **RhoB** and (b) **ER1** as a function of $\theta$ in water. The bottom panel shows the geometries, HOMO, and LUMO of these dyes in the locally excited (LE; $\theta = 0°$) and TICT states ($\theta = 90°$), respectively.

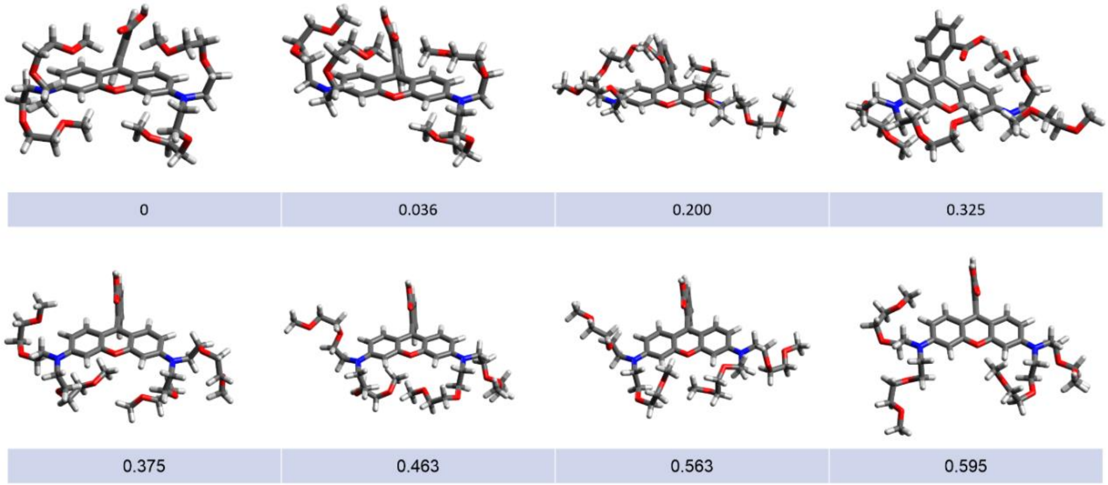

**Figure S6**. Various conformations of **ER2** and their relative electronic energy (eV) in water.

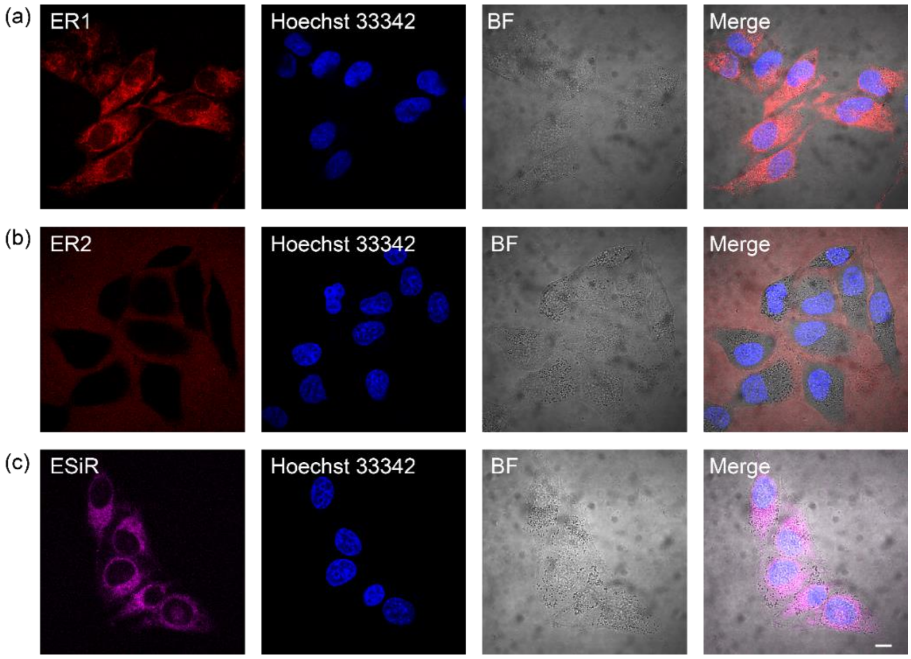

**Figure S7**. Confocal imaging of **ER1**, **ER2** and **ESiR** in living cells. Scale bar: 10 μm.

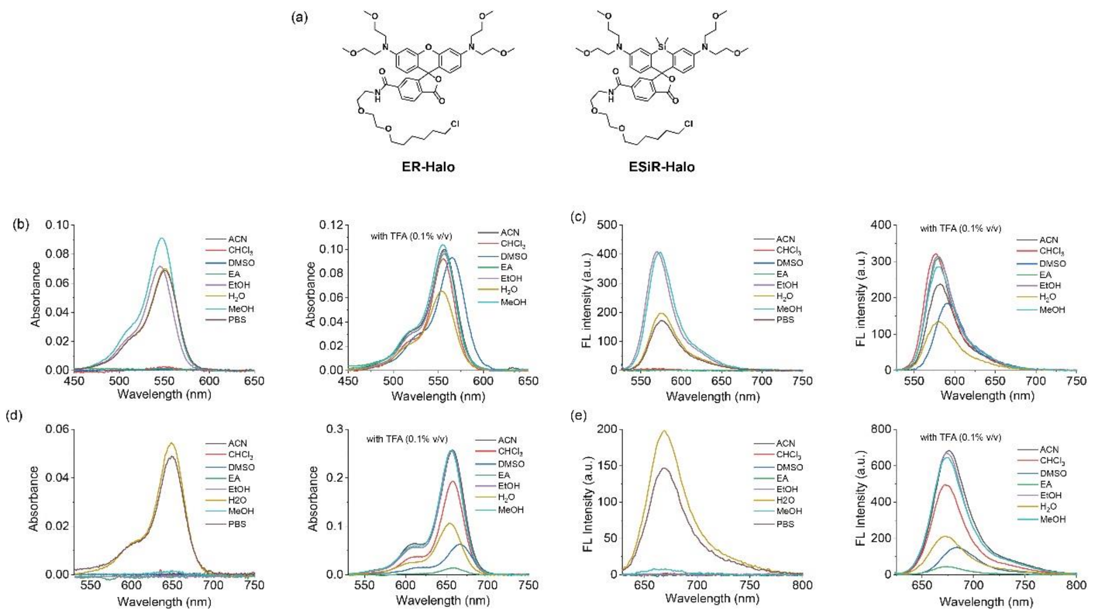

**Figure S8**. (a) Structure of ether chains decorated rhodamine-based probes **ER-Halo** and **ESiR-Halo**. (b, c) UV-Vis absorption (b) and fluorescence spectra (c) of **ER-Halo** in different solvents. (d, e) UV-Vis absorption (d) and fluorescence spectra (e) of **ESiR-Halo** in different solvents. The concentration of **ER-Halo** is 1 μM, and the concentration of **ESiR-Halo** is 2 μM.

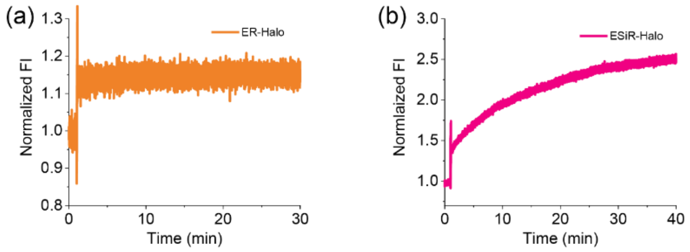

**Figure S9**. Time course of normalized fluorescence intensity of 1 μM probe in the presence of 2 μM Halo-tag.

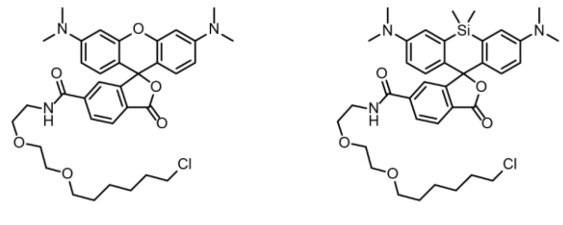

TMR-Halo TMSiR-Halo

**Figure S10**. Structure of N-methyl rhodamine-based probes **TMR-Halo** and **TMSiR-Halo**.

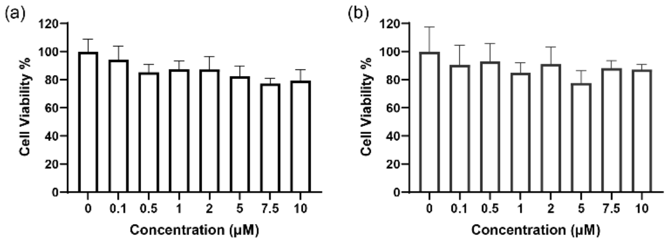

**Figure S11**. Cytotoxicity of **ER-Halo** (a) and **ESiR-Halo** (b) in MCF cells.

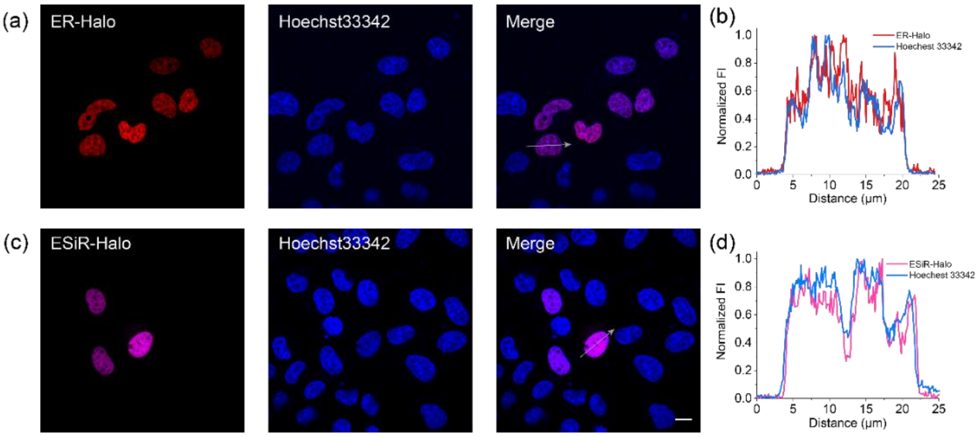

**Figure S12**. (a) Confocal imaging of nuclei using **ER-Halo** and Hoechst33342. (b) Intensity profile of regions of interest (ROI) across cells in (a). (c) Confocal imaging of nuclei using **ESiR-Halo** and Hoechst33342. (d) Intensity profile of ROI across cells in (c). HeLa cells transiently expressing Halo-H2B were incubated with 3 μM Hoechst and 100 nM ether chains decorated rhodamines-based probes for 1 h. Scale bar: 10 μm.

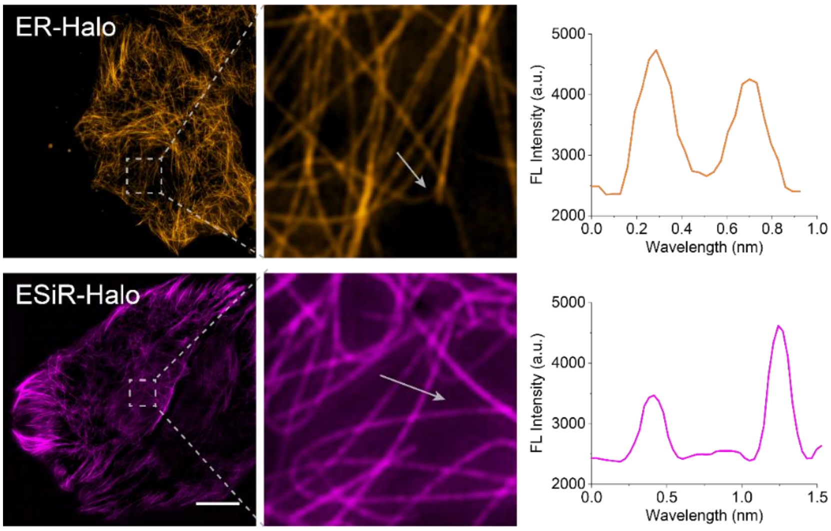

**Figure S13**. SIM imaging of tubulin in MCF cells expressed Halo-Tubulin using **ER-Halo** and **ESiR-Halo**. Graph: Intensity profile of ROI across cells. MCF cells transiently expressing Halo-Tubulin were incubated with 10 μM docetaxel and 100 nM probes for 1 h. Scale bar, 10 μm.

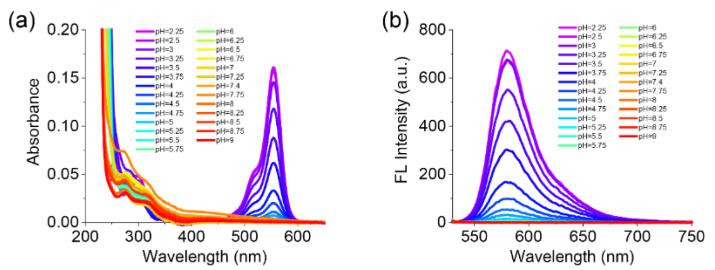

**Figure S14**. UV-Vis absorption (a) and fluorescence spectra (b) of **Lyso-ER** at different pH. [**Lyso-ER**] = 2 μM.

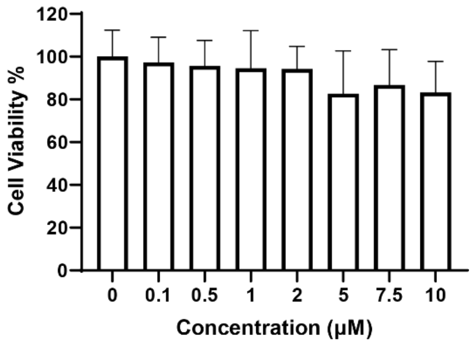

**Figure S15**. Cytotoxicity of **Lyso-ER** in MCF cells.

### Synthesis

Synthesis and characterization of **O-DAEA-1**

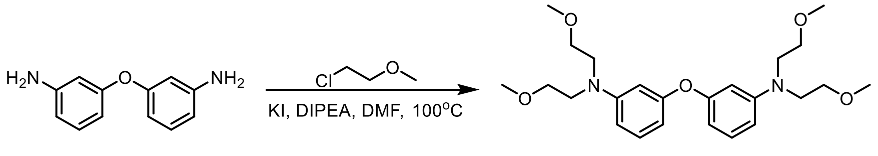

O-DAEA O-DAEA-1

**O-DAEA** (500 mg, 2.5 mmol), 2-methoxyethyl chloride (2.36 g, 25 mmol), DIPEA (731 mg, 25 mmol), KI (4.15 g, 25 mmol) were added in 8 mL dry DMF. The reaction mixture was stirred at 100 °C overnight. After the reaction was completed, the residue was poured into 20 ml of water. The organic phase was collected and dried with anhydrous sodium sulfate. The solvent was removed by evaporation. The crude product was purified by silica gel chromatography (PE/EA = 5:1; V/V) to afforded **O-DAEA-1** as a colorless viscous oily liquid 324 mg, yield 30%.

$^1$H NMR (400 MHz, $CDCl_3$) δ 7.10 (t, J = 8.1 Hz, 2H), 6.45–6.38 (m, 4H), 6.31–6.27 (m, 2H), 3.52 (s, 16H), 3.32 (s, 12H). $^{13}$C NMR (101 MHz, $CDCl_3$) δ 158.55, 149.44, 130.05, 106.77, 106.29, 102.70, 70.10, 58.97, 51.05. HRMS (ESI) calcd for $C_{24}H_{37}N_2O_5$ $[M+H]^+$ 433.2697, found 433.2676.

Synthesis and characterization of **O-DAEA-2**

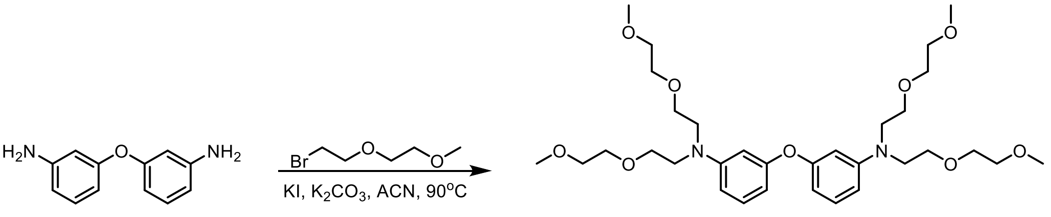

O-DAEA O-DAEA-2

**O-DAEA** (162 mg, 0.81 mmol), m-PEG-2-Br (1.18 g, 6.48 mmol), $K_2CO_3$ (900 mg, 6.48 mmol), KI (134 mg, 0.81 mmol) were placed in a 100 mL two-necked bottle. Then 10 mL of dry anhydrous $CH_3CN$ was added. The reaction was stirred at 90 °C for overnight. After the reaction was completed, the residue was poured into 20 ml of EA. The solvent was removed by evaporation. The crude product was purified by silica gel chromatography (PE/EA = 1:4; V/V) to afforded **O-DAEA-2** as a dark red viscous oily liquid 148 mg, yield 30%.

$^1$H NMR (400 MHz, $CDCl_3$) δ 7.10 (t, *J* = 8.2 Hz, 2H), 6.45 – 6.38 (m, 4H), 6.27 (dd, *J* = 8.0, 1.7 Hz, 2H), 3.66 – 3.47 (m, 32H), 3.38 (s, 12H). $^{13}$C NMR (101 MHz, $CDCl_3$) δ 158.52, 149.33, 130.00, 106.77, 106.21, 102.67, 71.94, 70.55, 68.38, 59.10, 51.03.

Synthesis and characterization of **ER1**

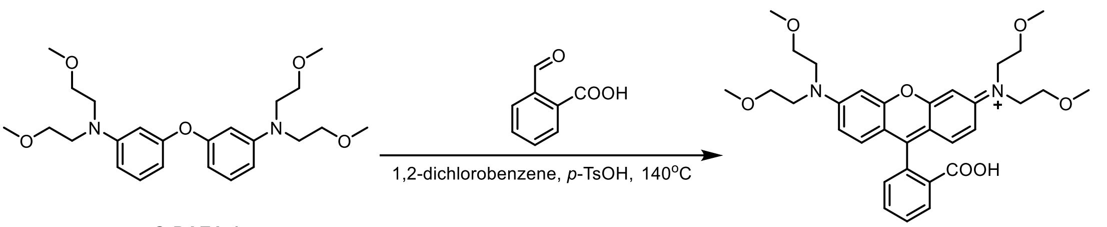

O-DAEA-1 ER1

To a 30 mL sealable pressure tube charged with a magnetic stir bar were added the intermediate **O-DAEA-1** (281 mg, 0.65 mmol), 2-Carboxybenzaldehyde (488 mg, 3.25 mmol), p-toluenesulfonic acid (112 mg, 0.65 mmol) and 1,2-dichlorobenzene 1 mL. The tube was sealed tightly and heated at 140 °C for 24 h. [Caution: To avoid potential danger, all the reaction tubes were placed behind a blast shield.] After cooling to room temperature, the crude product was purified by silica gel chromatography (DCM/MeOH = 10:1; V/V) to afforded **ER1** as a purple-red solid 207 mg, yield 56%.

$^{1}$H NMR (400 MHz, $CDCl_3$) δ 8.04 (d, J = 7.4 Hz, 1H), 7.62 (dt, J = 14.0, 7.3 Hz, 2H), 7.19 (d, J = 7.4 Hz, 1H), 6.63 (d, J = 8.9 Hz, 2H), 6.53 (d, J = 2.1 Hz, 2H), 6.45 (d, J = 8.9 Hz, 2H), 3.59 (d, J = 4.7 Hz, 8H), 3.56 (d, J = 4.5 Hz, 8H), 3.35 (s, 12H). HRMS (ESI) calcd for $C_{32}H_{39}N_2O_7$ $[M+H]^+$ 563.2752, found 563.2720.

Synthesis and characterization of **ER2**

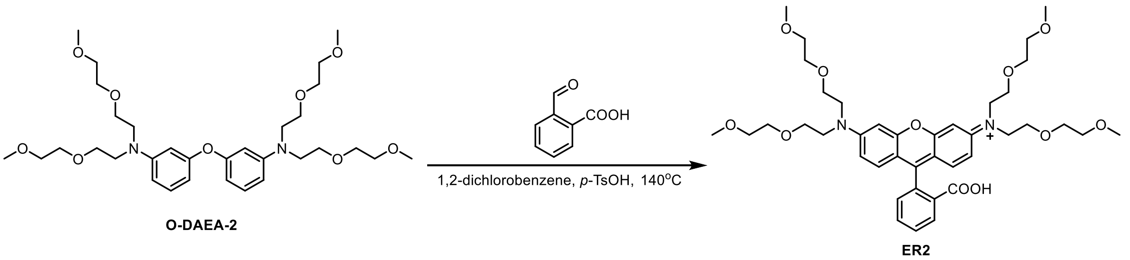

To a 30 mL sealable pressure tube charged with a magnetic stir bar were added the intermediate **O-DAEA-2** (37 mg, 0.05 mmol), 2-Carboxybenzaldehyde (40 mg, 0.27 mmol), p-toluenesulfonic acid (9 mg, 0.05 mmol) and 1,2-dichlorobenzene 1 mL. The tube was sealed tightly and heated at 140 °C for 24 h. [Caution: To avoid potential danger, all the reaction tubes were placed behind a blast shield.] After cooling to room temperature, the crude product was purified by silica gel chromatography (DCM/MeOH = 10:1; V/V) to afforded **ER2** as a purple-red solid 18 mg, yield 45%.

$^{1}$H NMR (400 MHz, CDCl3) δ 8.02 (d, J = 7.4 Hz, 1H), 7.62 (dt, J = 22.5, 7.3 Hz, 2H), 7.19 (d, J = 7.4 Hz, 1H), 6.58 (d, J = 8.9 Hz, 2H), 6.51 (d, J = 1.7 Hz, 2H), 6.41 (d, J = 8.8 Hz, 2H), 3.74–3.56 (m, 24H), 3.57–3.47 (m, 8H), 3.37 (s, 12H). $^{13}$C NMR (101 MHz, CDCl3) δ 169.58, 153.45, 150.05, 134.35, 129.34, 129.12, 128.01, 125.22, 124.40, 108.56, 107.19, 97.99, 71.95, 70.65, 68.29, 59.10, 51.11, 29.70. HRMS (ESI) calcd for $C_{40}H_{55}N_2O_{11}$ $[M+H]^+$ 739.3800, found 739.3830.

Synthesis and characterization of **ER1-Ester**

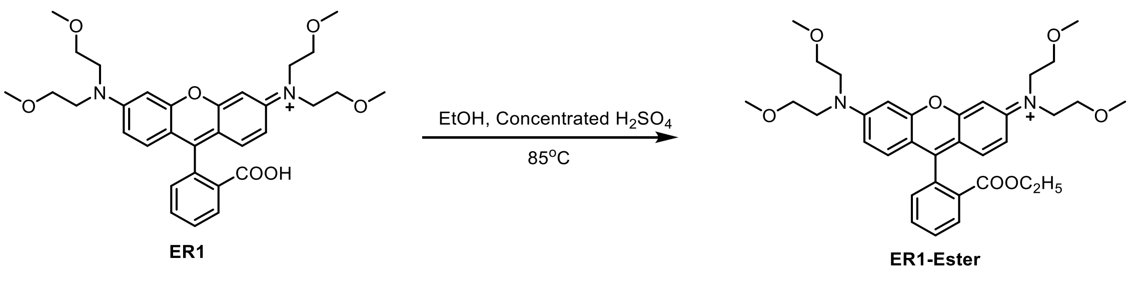

Concentrated sulfuric acid (1 mL, 1.84 g/cm$^3$) was added dropwise to the solution of **ER1** (125 mg, 0.22 mmol) in 10 mL ethanol. The reaction was heated tightly at 85 °C for 9 h. After the reaction was completed, the residue was slowly poured into 100 ml of ice-water mixture. Then it was slowly added saturated sodium carbonate solution to pH neutral. Then it was filtered and washed three times with DCM. The lower organic phase is collected and dried with anhydrous sodium sulfate. The solvent was removed by evaporation. The crude product was purified by silica gel chromatography (DCM/EA/MeOH = 4:4:1; V/V/V) to afforded **ER1-Ester** as a purple solid 96 mg, yield 73%.

$^1$H NMR (700 MHz, MeOD) δ 8.30 (d, J = 7.9 Hz, 1H), 7.87 (t, J = 7.4 Hz, 1H), 7.81 (t, J = 7.7 Hz, 1H), 7.44 (d, J = 7.5 Hz, 1H), 7.18–7.10 (m, 6H), 4.02 (q, J = 7.1 Hz, 2H), 3.89 (t, J = 5.1 Hz, 8H), 3.69 (s, 8H), 3.34 (s, 12H), 0.97 (t, J = 7.1 Hz, 3H). $^{13}$C NMR (176 MHz, MeOD) δ 165.17, 159.38, 157.71, 157.26, 133.38, 132.74, 130.86, 130.59, 130.39, 130.24, 130.21, 114.98, 113.83, 96.94, 69.87, 63.70, 63.68, 63.66, 61.17, 58.07, 58.06, 51.42, 14.05, 12.69. HRMS (ESI) calcd for $C_{34}H_{43}N_2O_7$ $[M]^+$ 591.3065, found 591.3024.

Synthesis and characterization of **ER2-Ester**

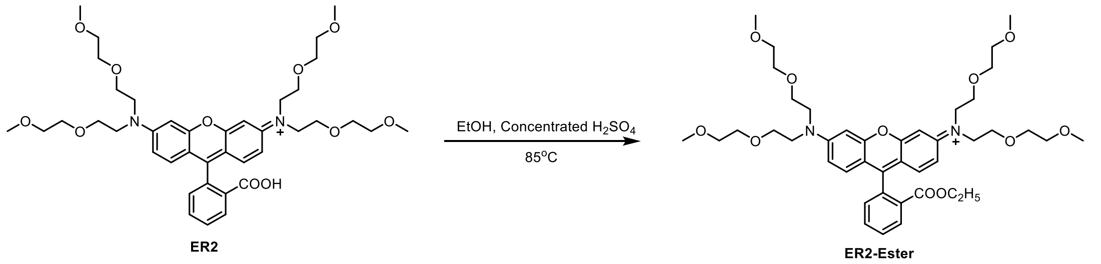

Concentrated sulfuric acid (1 mL, 1.84 g/cm$^3$) was added dropwise to the solution of **ER2** (36 mg, 0.048 mmol) in 10 mL methanol. The reaction was heated tightly at 85 °C for 9 h. After the reaction was completed, the residue was slowly poured into 100 ml of ice-water mixture. Then it was slowly added saturated sodium carbonate solution to pH neutral. Then it was filtered and washed three times with DCM. The lower organic phase is collected and dried with anhydrous sodium sulfate. The solvent was removed by evaporation. The crude product was purified by silica gel chromatography (DCM/EA/MeOH = 3:3:1; V/V/V) to afforded **ER2-Ester** as a purple solid 28 mg, yield 75%.

$^1$H NMR (400 MHz, MeOD) δ 8.31 (d, J = 7.3 Hz, 1H), 7.85 (dt, J = 24.9, 7.3 Hz, 2H), 7.72 (d, J = 8.0 Hz, 1H), 7.45 (d, J = 7.2 Hz, 1H), 7.25 (d, J = 7.9 Hz, 1H), 7.14 (dd, J = 19.4, 7.9 Hz, 4H), 4.07–4.07 (m, 2H), 3.92 (d, J = 4.9 Hz, 8H), 3.79 (s, 8H), 3.61 (d, J = 4.5 Hz, 8H), 3.51 (d, J = 4.4 Hz, 8H), 3.31 (s, 12H), 1.29 (s, 3H). HRMS (ESI) calcd for $C_{42}H_{59}N_2O_{11}$ $[M]^+$ 767.4113, found 767.4093.

Synthesis and characterization of compound **1**

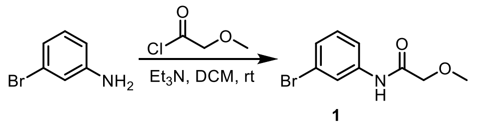

3-Bromoaniline (2.0 g, 11.6 mmol) was added dropwise to the solution of triethylamine (3.53 g, 34.8 mmol) in 8 mL DCM. It was added slowly Methoxyacetyl chloride (1.89 g, 17.4 mmol) dissolved in DCM at 0 °C. And the reaction was stirred at room temperature for 5 h, while monitored by TLC. After the reaction was completed, the mixture was added water for quenching reaction. Then it was washed by saturated sodium bicarbonate solution and water, respectively. Then it was filtered and washed three times with DCM. The lower organic phase is collected and dried with anhydrous sodium sulfate. The solvent was removed by evaporation. The crude product was purified by silica gel chromatography (pure DCM) to afforded compound **1** as a yellow viscous oily liquid 2.8 g, yield 99%.

$^1$H NMR (700 MHz, $CDCl_3$) δ 8.19 (s, 1H), 7.76 (t, *J* = 1.9 Hz, 1H), 7.42 (ddd, *J* = 8.1, 1.9, 0.8 Hz, 1H), 7.18 (ddd, *J* = 8.0, 1.8, 1.0 Hz, 1H), 7.12 (t, *J* = 8.0 Hz, 1H), 3.94 (s, 2H), 3.43 (s, 3H).

Synthesis and characterization of compound **2**

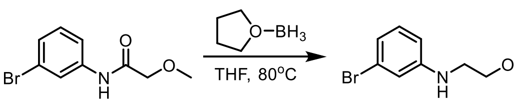

**1** **2**

Compound **1** (1.27 g 5.21 mmol) was placed in a 100 mL double-mouth bottle and vacuumed by nitrogen. It was added 8 mL anhydrous tetrahydrofuran in a nitrogen-protected atmosphere. Then it was added borane tetrahydrofuran complex solution (0.86 g 10.42 mmol) was added slowly at 0 °C. The reaction was heated and refluxed at 80 °C for overnight. After the reaction was completed, the mixture was added methanol for quenching reaction. The reaction solvent was removed by evaporation. The residue was diluted by 20 mL EA. Then it was washed by 10 mL 3 M $HCl_{(aq)}$. It was slowly added saturated sodium carbonate solution to pH neutral. Then it was filtered and washed three times with EA. The upper organic phase is collected and dried with anhydrous sodium sulfate. The solvent was removed by evaporation. The crude product was purified by silica gel chromatography (pure DCM) to afforded compound **2** as a yellow viscous oily liquid 1.19 g, yield 99%.

$^1$H NMR (700 MHz, $CDCl_3$) δ 6.93 (t, *J* = 8.0 Hz, 1H), 6.74 (d, *J* = 7.8 Hz, 1H), 6.68 (s, 1H), 6.45 (dd, *J* = 8.2, 1.8 Hz, 1H), 4.04 (s, 1H), 3.52 (t, *J* = 5.2 Hz, 2H), 3.31 (s, 3H), 3.18 (t, *J* = 5.0 Hz, 2H).

Synthesis and characterization of compound **3**

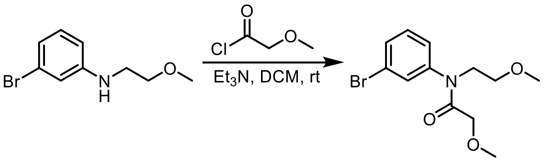

**2** **3**

Compound **2** (1.19 g, 5.2 mmol) was added dropwise to the solution of triethylamine (1.57 g, 15.5 mmol) in 5 mL DCM. It was added slowly Methoxyacetyl chloride (0.84 g, 7.78 mmol) dissolved in DCM at 0 °C. And the reaction was stirred at room temperature for 5 h, while monitored by TLC. After the reaction was completed, the mixture was added water for quenching reaction. Then it was washed by saturated sodium bicarbonate solution and water, respectively. Then it was filtered and washed three times with DCM. The lower organic phase is collected and dried with anhydrous sodium sulfate. The solvent was removed by evaporation. The crude product was purified by silica gel chromatography (PE/EA = 10:1; V/V) to afforded compound **3** as a yellow viscous oily liquid 1.5 g, yield 96%.

$^1$H NMR (700 MHz, $CDCl_3$) δ 7.43 (d, *J* = 7.7 Hz, 1H), 7.37 (s, 1H), 7.24 (t, *J* = 8.0 Hz, 1H), 7.15 (d, *J* = 7.9 Hz, 1H), 3.78 (t, *J* = 5.5 Hz, 2H), 3.69 (s, 2H), 3.45 (s, 2H), 3.23 (d, *J* = 21.4 Hz, 6H).

Synthesis and characterization of compound **4**

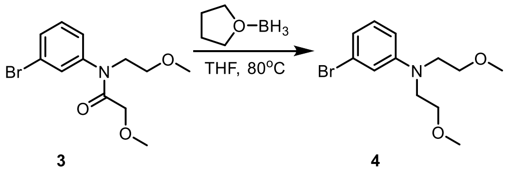

**3** **4**

Compound **3** (1.4 g 4.6 mmol) was placed in a 100 mL double-mouth bottle and vacuumed by nitrogen. It was added 8 mL anhydrous tetrahydrofuran in a nitrogen-protected atmosphere. Then it was added borane tetrahydrofuran complex solution (0.80 g 9.3 mmol) was added slowly at 0 °C. The reaction was heated and refluxed at 80 °C for overnight. After the reaction was completed, the mixture was added methanol for quenching reaction. The reaction solvent was removed by evaporation. The residue was diluted by 20 mL EA. Then it was washed by 10 mL 3 M $HCl_{(aq)}$. It was slowly added saturated sodium carbonate solution to pH neutral. Then it was filtered and washed three times with EA. The upper organic phase is collected and dried with anhydrous sodium sulfate. The solvent was removed by evaporation. The crude product was purified by silica gel chromatography (PE/EA = 20:1; V/V) to afforded compound **4** as a yellow viscous oily liquid 1.05 g, yield 79%.

$^1$H NMR (700 MHz, $CDCl_3$) δ 6.95 (t, *J* = 8.1 Hz, 1H), 6.74 (s, 1H), 6.69 (d, *J* = 7.8 Hz, 1H), 6.54 (dd, *J* = 8.4, 1.9 Hz, 1H), 3.45 (s, 8H), 3.27 (s, 6H).

Synthesis and characterization of **Si-DAEA**

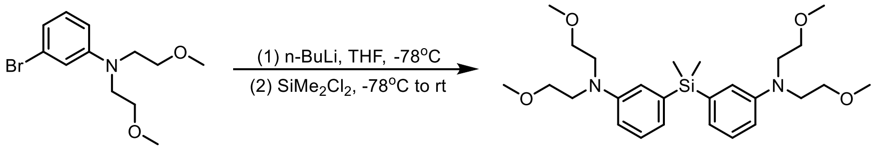

Compound **4** (1.28 g 4.45 mmol) was placed in a 50 mL schlenk and vacuumed by nitrogen. It was added 10 mL anhydrous tetrahydrofuran in a nitrogen-protected atmosphere. Then it was added n-Butyllithium (0.28 g 4.45 mmol) was added slowly at -78 °C. The reaction was performed for 0.5 h at -78 °C. Then Dichlorodimethylsilane (0.28 g, 2.23 mmol) was added dropwise to the reaction system. The reaction was performed at room temperature for overnight. After the reaction was completed, the mixture was added saturated $NH4Cl_{(aq)}$ for quenching reaction. The residue was diluted by 20 mL water. Then it was filtered and washed three times with EA. The upper organic phase is collected and dried with anhydrous sodium sulfate. The solvent was removed by evaporation. The crude product was purified by silica gel chromatography (PE/EA = 8:1; V/V) to afforded **Si-DAEA** as a yellow viscous oily liquid 422 g, yield 20%.

$^1$H NMR (400 MHz, $CDCl_3$) δ 7.10 – 7.03 (m, 2H), 6.73 (t, *J* = 4.4 Hz, 4H), 6.58 (dd, *J* = 8.6, 2.5 Hz, 2H), 3.43 – 3.34 (m, 16H), 3.19 (s, 12H), 0.38 (s, 6H). HRMS (ESI) calcd for $C_{26}H_{43}N_2O_4Si$ $[M+H]^+$ 475.2992, found 475.2998.

Synthesis and characterization of **ESiR**

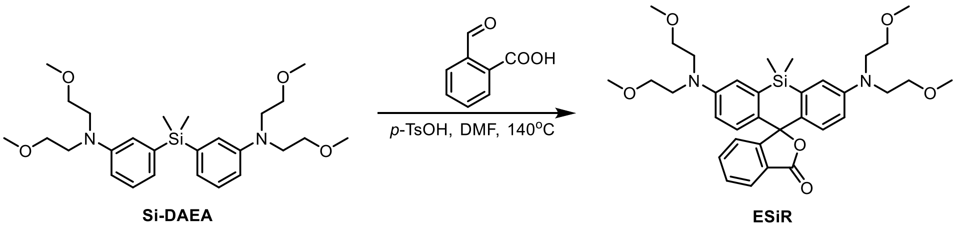

To a 30 mL sealable pressure tube charged with a magnetic stir bar were added the intermediate **Si-DAEA** (50 mg, 0.10 mmol), 2-Carboxybenzaldehyde (80 mg, 0.52 mmol), p-toluenesulfonic acid (20 mg, 0.10 mmol) and DMF 200 μL. The tube was sealed tightly and heated at 140 °C for 8 h. [Caution: To avoid potential danger, all the reaction tubes were placed behind a blast shield.] After cooling to room temperature, the mixture was diluted with 2 mL MeOH, then added chloranil (49 mg, 0.2 mmol) and stirred for 2 h. After filtration and removal of the solvent, the crude product was purified by silica gel chromatography (PE/EA = 2:1; V/V) to afforded **ESiR** as a white solid 10 mg, yield 16%.

$^1$H NMR (400 MHz, $CDCl_3$) δ 7.96 (d, *J* = 7.6 Hz, 1H), 7.65 (t, *J* = 7.1 Hz, 1H), 7.55 (t, *J* = 7.4 Hz, 1H), 7.32 (d, *J* = 7.6 Hz, 1H), 6.99 (d, *J* = 2.8 Hz, 2H), 6.73 (d, *J* = 8.9 Hz, 2H), 6.52 (dd, J = 9.0, 2.9 Hz, 2H), 3.54 (dd, J = 12.6, 4.5 Hz, 16H), 3.34 (s, 12H), 0.60 (d, J = 6.3 Hz, 6H). $^{13}$C NMR (101 MHz, CDCl3) δ 148.47, 138.99, 135.26, 133.37, 130.42, 130.09, 127.35, 126.46, 117.95, 114.31, 71.81, 60.75, 52.54, 31.42, 2.05. HRMS (ESI) calcd for $C_{34}H_{45}N_2O_6Si$ $[M+H]^+$ 605.3047, found 605.3067.

Synthesis and characterization of **ESiR-Me**

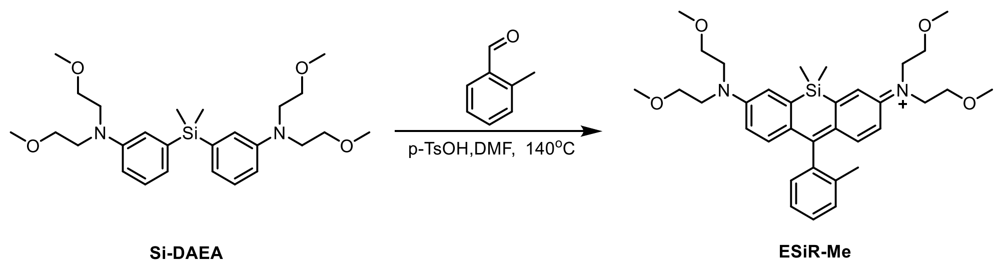

To a 30 mL sealable pressure tube charged with a magnetic stir bar were added the intermediate **Si-DAEA** (50 mg, 0.10 mmol), o-Tolualdehyde (63 mg, 0.52 mmol), p-toluenesulfonic acid (20 mg, 0.10 mmol) and DMF 200 μL. The tube was sealed tightly and heated at 140 °C for 8 h. After cooling to room temperature, the mixture was diluted with 2 mL MeOH, then added chloranil (49 mg, 0.2 mmol) and stirred for 2 h. After filtration and removal of the solvent, the crude product was purified by silica gel chromatography (DCM/MeOH = 10:1; V/V) to afforded **ESiR-Me** as a blue solid 18 mg, yield 30%.

$^1$H NMR (400 MHz, MeOD) δ 7.49 (d, *J* = 2.8 Hz, 2H), 7.47 – 7.43 (m, 1H), 7.38 (dd, *J* = 16.1, 8.0 Hz, 2H), 7.13 (d, *J* = 7.2 Hz, 1H), 7.07 (d, *J* = 9.7 Hz, 2H), 6.85 (dd, *J* = 9.7, 2.8 Hz, 2H), 3.92 (d, *J* = 4.6 Hz, 8H), 3.66 (t, *J* = 5.0 Hz, 8H), 3.36 (d, *J* = 14.7 Hz, 12H), 0.60 (s, 3H), 0.58 (s, 3H).$^{13}$C NMR (101 MHz, MeOD) δ 172.53, 157.24, 150.64, 143.96, 141.52, 138.50, 132.86, 131.66, 130.43, 128.31, 124.87, 117.35, 73.00, 60.92, 54.20, 21.08, 0.22. HRMS (ESI) calcd for $C_{34}H_{45}N_2O_6Si$ $[M]^+$ 575.3300, found 575.3311.

Synthesis and characterization of **ER-COOH**

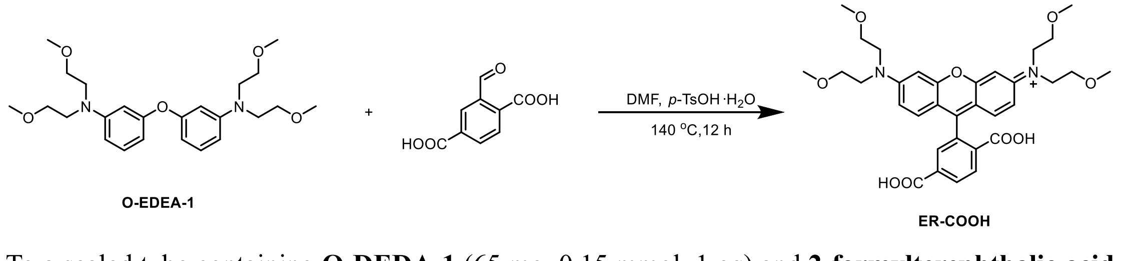

To a sealed tube containing **O-DEDA-1** (65 mg, 0.15 mmol, 1 eq) and **2-formylterephthalic acid** (146 mg, 0.75 mmol, 5 eq) and ***p*-TsOH·H₂O** (29mg, 0.15 mmol, 1 eq), DMF (5mL) was added, and the mixture was stirred at 140 °C for 12 h. The crude material was purified by silica gel flash column chromatography (0-50% MeOH/$CH_2Cl_2$) affording compound **ER-COOH** as purple solid (50 mg, 56%).

$^1$H NMR (400 MHz, MeOD) δ 8.24 (dd, *J* = 18.6, 7.9 Hz, 2H), 7.86 (s, 1H), 7.15 (d, *J* = 9.3 Hz, 2H), 7.10 – 7.01 (m, 4H), 3.84 (t, *J* = 5.2 Hz, 8H), 3.64 (t, *J* = 5.2 Hz, 8H), 3.30 (d, *J* = 7.6 Hz, 12H). HRMS calcd for $C_{33}H_{39}N_2O_9^+$ $[M]^+$ 607.2650, found 607.2676.

Synthesis and characterization of **ER-Halo**

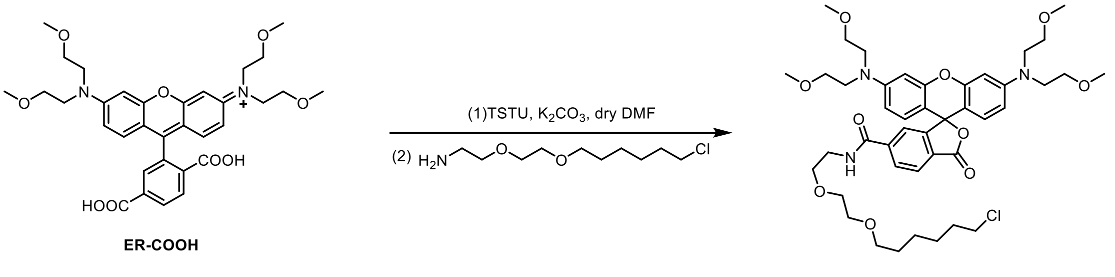

To a schlenk flask containing **TSTU** (12 mg, 0.04 mmol, 1 eq), **$K_2CO_3$** (28 mg, 0.2 mmol, 5 eq) and compound **ER-COOH** (25 mg, 0.04 mmol, 1 eq), the atmosphere was purged with $N_2$ by three cycles of vacuum/backfilling. Dry DMF (5mL) was added, the mixture was stirred at room temperature for 30 min. Then **2-(2-((6-chlorohexyl)oxy)ethoxy)ethan-1-amine** (9 mg, 0.04 mmol, 1 eq) was added, and the reaction mixture was stirred at room temperature overnight. The crude material was purified by silica gel flash column chromatography (0-10% MeOH/$CH_2Cl_2$) affording **ER-Halo** as purple solid (12 mg, 36%).

$^1$H NMR (400 MHz, MeOD) δ 8.10 (d, $J$ = 7.9 Hz, 1H), 8.01 (dd, $J$ = 8.1, 1.6 Hz, 1H), 7.62 (d, $J$ = 1.6 Hz, 1H), 7.12 (dd, $J$ = 6.9, 4.8 Hz, 2H), 6.98 (dd, $J$ = 13.0, 6.0 Hz, 4H), 3.77 (t, $J$ = 5.2 Hz, 8H), 3.58 – 3.46 (m, 16H), 3.41 (dd, $J$ = 9.3, 4.0 Hz, 2H), 3.32 (t, $J$ = 6.5 Hz, 2H), 3.21 (dt, $J$ = 3.2, 1.6 Hz, 12H), 1.93 (dd, $J$ = 12.0, 6.3 Hz, 2H), 1.63 – 1.58 (m, 2H), 1.53 – 1.47 (m, 2H), 1.40 – 1.37 (m, 2H). HRMS calcd for $C_{43}H_{59}ClN_3O_{10}^+$ $[M]^+$ 812.3883, found 812.3884.

Synthesis and characterization of **ESiR-COOH**

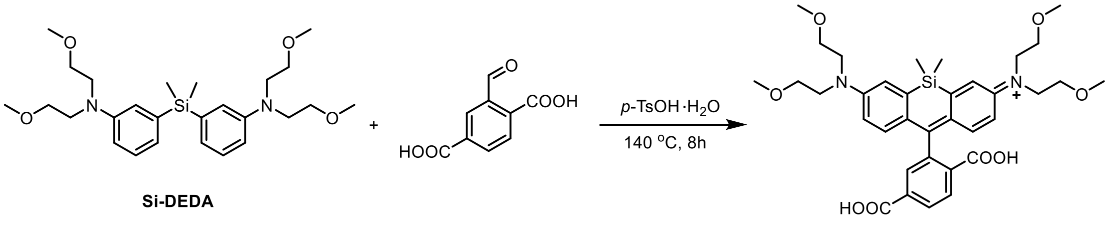

To a sealed tube containing **Si-DEDA** (350 mg, 0.738 mmol, 1 eq), 2-formylterephthalic acid (716 mg, 3.69 mmol, 5 eq) and p-TsOH·$H_2O$ (140mg, 0.738 mmol, 1 eq), DMF (1mL) was added, and the mixture was stirred at 140 °C for 5 h. The crude material was purified by silica gel flash column chromatography (0-2.5% MeOH/DCM) affording **ESiR-COOH** as bule solid (110 mg).

HRMS calcd for $C_{35}H_{35}N_2O_8Si^+$ $[M]^+$ 649.2940, found 649.2955.

Synthesis and characterization of **ESiR-NHS**

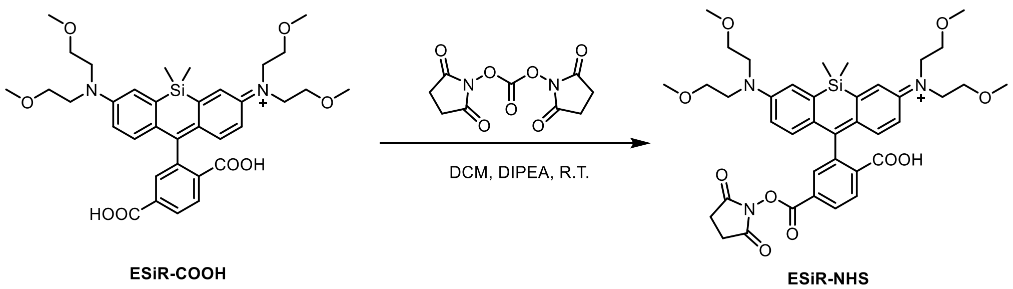

To a round-bottomed flask containing **ESiR-COOH** (110mg, 0.17 mmol, 1eq), DIPEA (109mg, 0.85 mmol, 5eq) and DCM (10mL), DSC (217 mg, 0.85 mmol, 5eq) was added. The reaction mixture was stirred at room temperature overnight. The solvent was removed by evaporation. The crude product was purified by silica gel chromatography (0-50%, EA/DCM) to afforded **ESiR-NHS** as a yellow solid (25 mg, 20%).

$^1$H NMR (400 MHz, $CDCl_3$) δ 8.29 (dd, $J$ = 8.0, 1.1 Hz, 1H), 8.08 (d, $J$ = 8.0 Hz, 1H), 8.02 (s, 1H), 6.99 (d, $J$ = 2.8 Hz, 2H), 6.71 (d, $J$ = 8.9 Hz, 2H), 6.56 (dd, $J$ = 9.0, 2.8 Hz, 2H), 3.55 (dt, $J$ = 10.4, 5.3 Hz, 16H), 3.35 (s, 12H), 2.90 (s, 4H), 0.62 (s, 3H), 0.57 (s, 3H). $^{13}$C NMR (101 MHz, $CDCl_3$) δ 170.70, 170.32, 162.47, 156.59, 148.40, 138.24, 133.39, 132.05, 131.67, 131.33, 129.71, 128.18, 127.55, 117.71, 114.41, 93.72, 71.52, 60.48, 52.27, 27.08, 1.58, 0.00. HRMS calcd for $C_{39}H_{48}N_3O_{10}Si^+$ $[M]^+$ 746.3103, found 746.3111.

Synthesis and characterization of **ESiR-Halo**

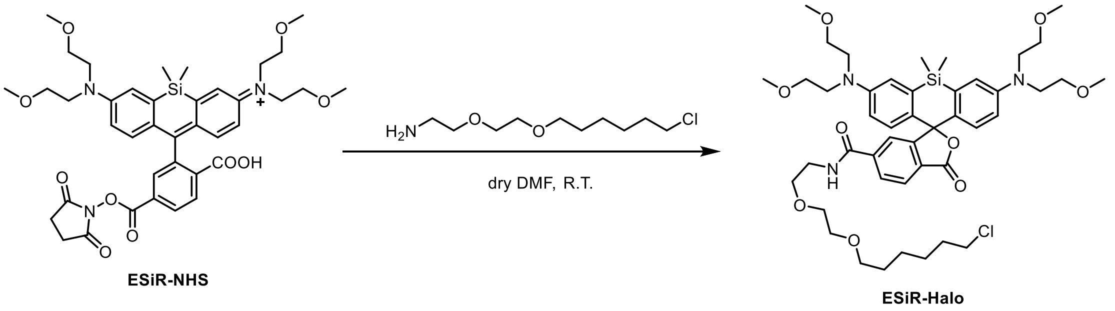

To a schlenk flask containing **ESiR-NHS** (25 mg, 0.033 mmol, 1 eq) and **2-(2-((6-chlorohexyl)oxy)ethoxy)ethan-1-amine** (22.5 mg, 0.1 mmol, 3 eq), the atmosphere was purged with $N_2$ by three cycles of vacuum/backfilling. Dry DMF (5mL) was added, the mixture was stirred at room temperature overnight. The solvent was removed by evaporation. The crude product was purified by silica gel chromatography (0-2%, MeOH/DCM) to afforded **ESiR-Halo** as a colorless liquid (15 mg, 53%).

$^1$H NMR (400 MHz, $CDCl_3$) δ 7.99 (d, J = 7.9 Hz, 1H), 7.90 (d, J = 7.9 Hz, 1H), 7.71 (s, 1H), 6.98 (d, J = 2.8 Hz, 2H), 6.73 (d, J = 8.9 Hz, 2H), 6.54 (dd, J = 9.0, 2.8 Hz, 2H), 3.63 (dd, J = 6.6, 3.9 Hz, 8H), 3.56 (d, J = 4.7 Hz, 8H), 3.52 (dd, J = 9.7, 5.7 Hz, 10H), 3.41 (t, J = 6.7 Hz, 2H), 3.34 (s, 12H), 2.01 (dd, J = 12.4, 6.4 Hz, 2H), 1.74 (dd, J = 14.1, 7.2 Hz, 2H), 1.58 – 1.49 (m, 2H), 1.40 (dd, J = 15.3, 7.5 Hz, 2H), 0.63 (s, 3H), 0.57 (s, 3H). $^{13}$C NMR (101 MHz, $CDCl_3$) δ 171.35, 167.67, 156.56, 148.29, 141.16, 138.28, 132.32, 130.60, 129.80, 128.76, 127.19, 124.98, 117.65, 114.32, 93.49, 72.67, 72.67, 71.71, 71.55, 71.44, 70.99, 70.99, 60.49, 52.28, 46.43, 41.44, 33.92, 30.82, 28.07, 26.81, 1.62, 0.11 . HRMS calcd for $C_{45}H_{65}ClN_3O_9Si^+$ $[M]^+$ 854.4173, found 854.4200.

Synthesis and characterization of **Lyso-ER**

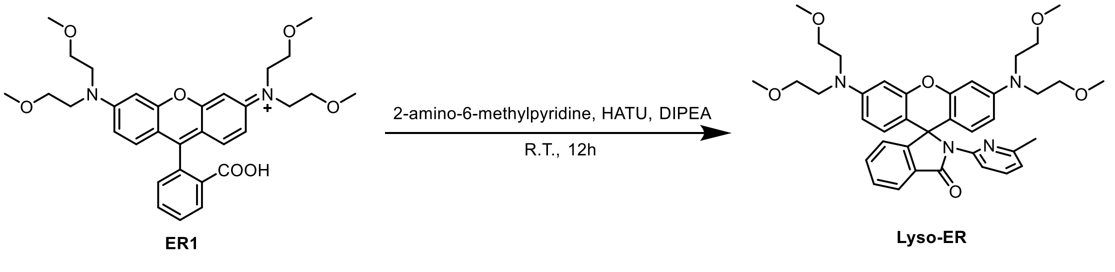

To a schlenk flask containing **ER1** (35 mg, 0.062 mmol, 1 eq), **HATU** (28mg, 0.075mmol, 1.2 eq) and **2-amino-6-methylpyridine** (23 mg, 0.217 mmol, 3.5 eq), the atmosphere was purged with $N_2$ by three cycles of vacuum/backfilling. Dry DMF (5mL) and DIPEA (40mg, 0.31mmol, 5eq) was added, the mixture was stirred at room temperature overnight. The solvent was removed by evaporation. The crude product was purified by silica gel chromatography (0-3%, MeOH/DCM) to afforded **Lyso-ER** as a colorless liquid (15 mg, 37%).

$^{1}$H NMR (400 MHz, $CDCl_3$) δ 8.27 (d, *J* = 8.4 Hz, 1H), 8.00 (dd, *J* = 6.1, 2.0 Hz, 1H), 7.54 – 7.47 (m, 2H), 7.42 – 7.36 (m, 1H), 7.13 (dd, *J* = 6.0, 1.6 Hz, 1H), 6.62 (d, *J* = 7.4 Hz, 1H), 6.43 (d, *J* = 2.5 Hz, 2H), 6.39 (d, *J* = 8.8 Hz, 2H), 6.15 (dd, *J* = 8.8, 2.6 Hz, 2H), 3.59 – 3.44 (m, 16H), 3.33 (s, 12H), 2.21 (s, 3H). $^{13}$C NMR (101 MHz, $CDCl_3$) δ 167.97, 155.79, 153.88, 153.34, 149.57, 148.50 , 137.07, 133.44, 131.06, 128.20, 127.95, 124.47, 123.13, 118.06, 111.79, 109.86, 106.98, 97.84, 70.06, 65.95, 58.99, 50.92, 23.09. HRMS calcd for $C_{38}H_{45}N_4O_6[M+H]^+$ 653.3334, found 653.3344.

### NMR and HRMS Spectra

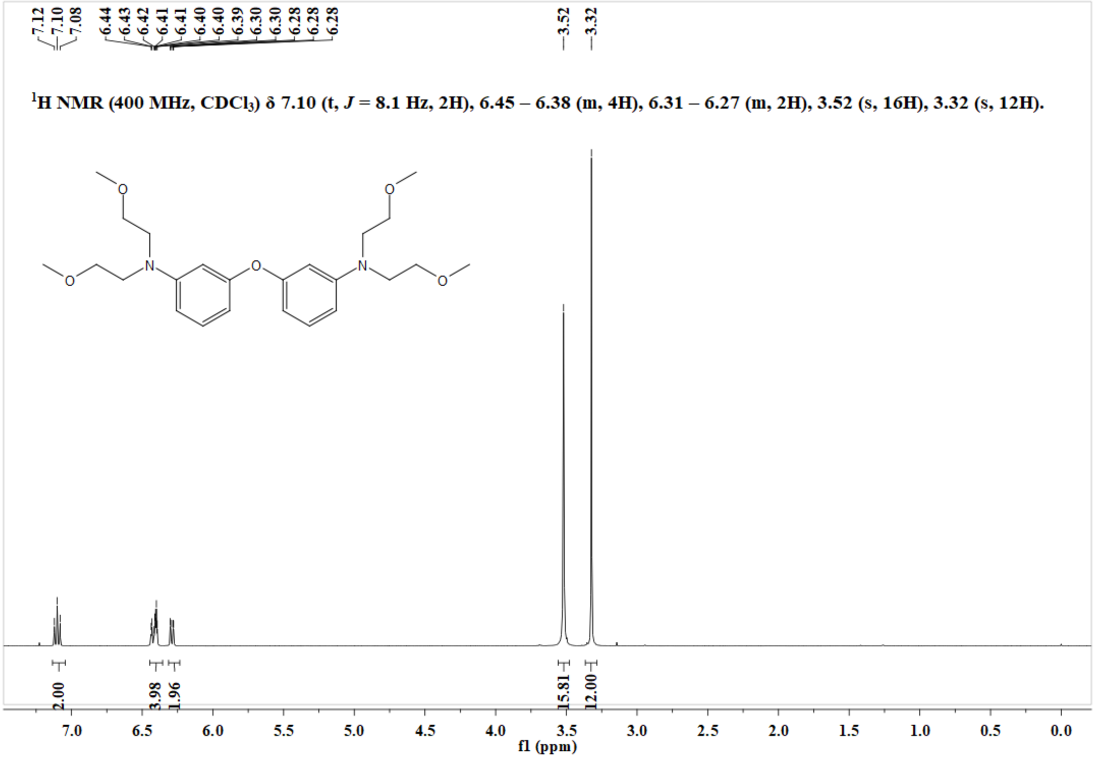

**Figure S16**. $^{1}$H NMR spectrum of **O-DAEA-1** in $CDCl_3$.

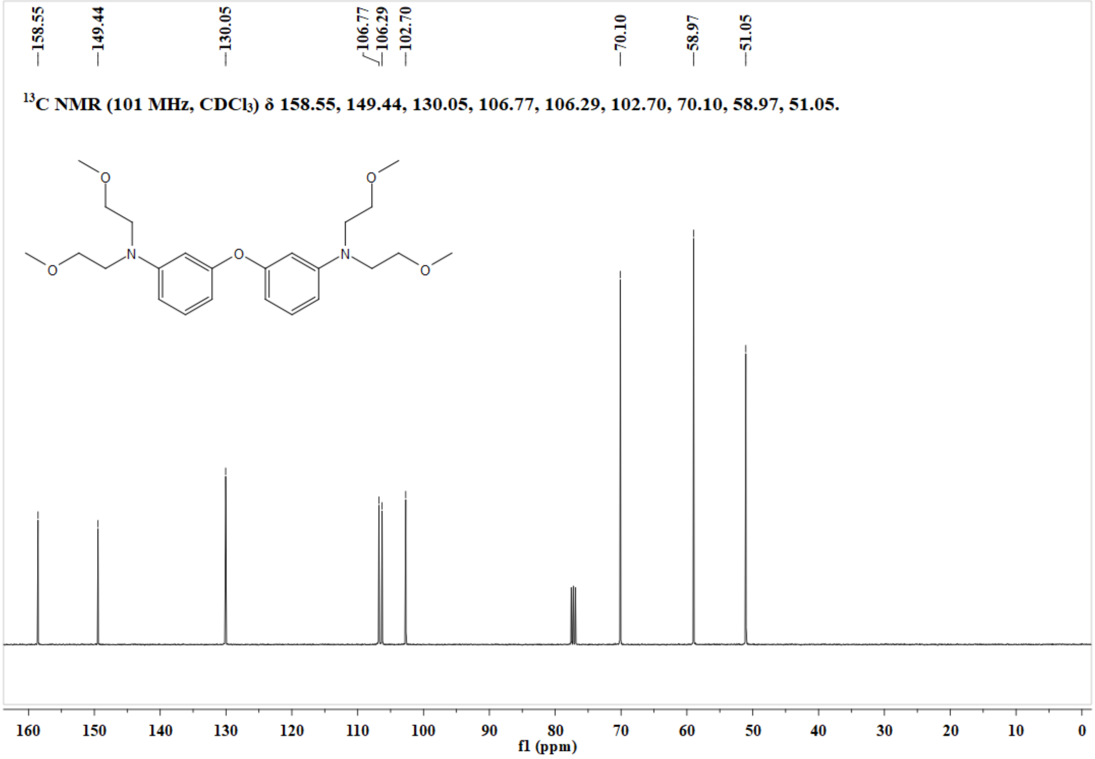

**Figure S17**. $^{13}$C NMR spectrum of **O-DAEA-1** in $CDCl_3$.

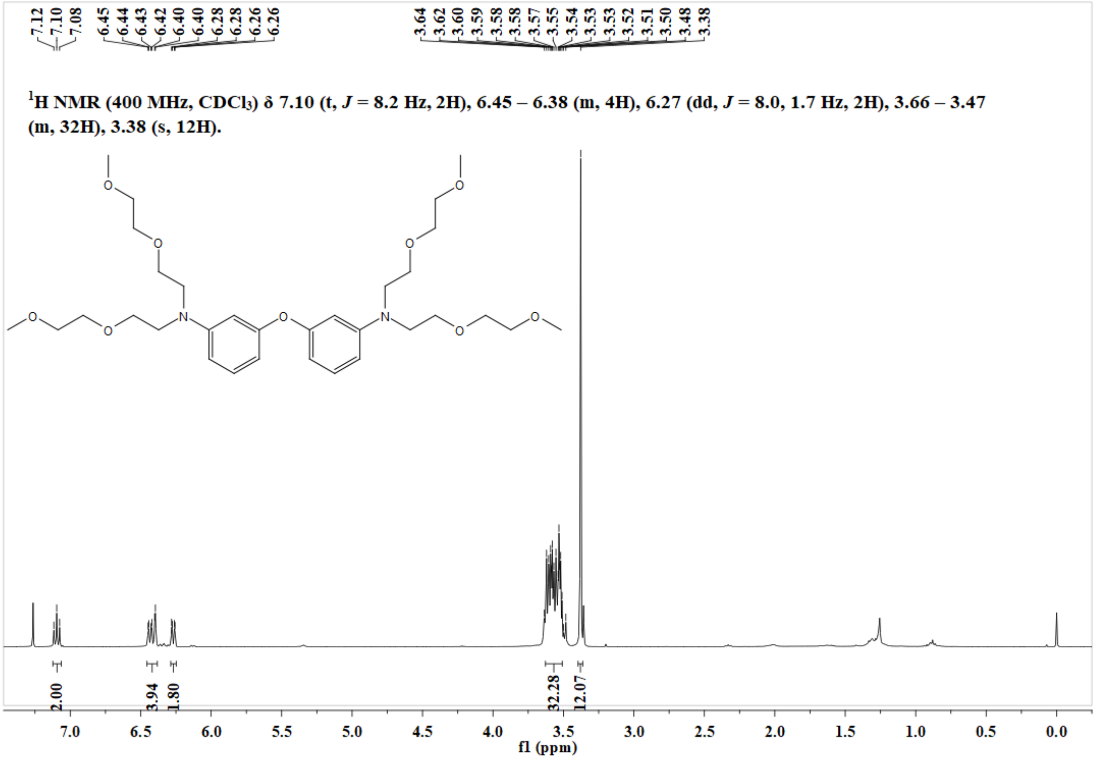

**Figure S18**. $^{1}H$ NMR spectrum of **O-DAEA-2** in $CDCl_3$.

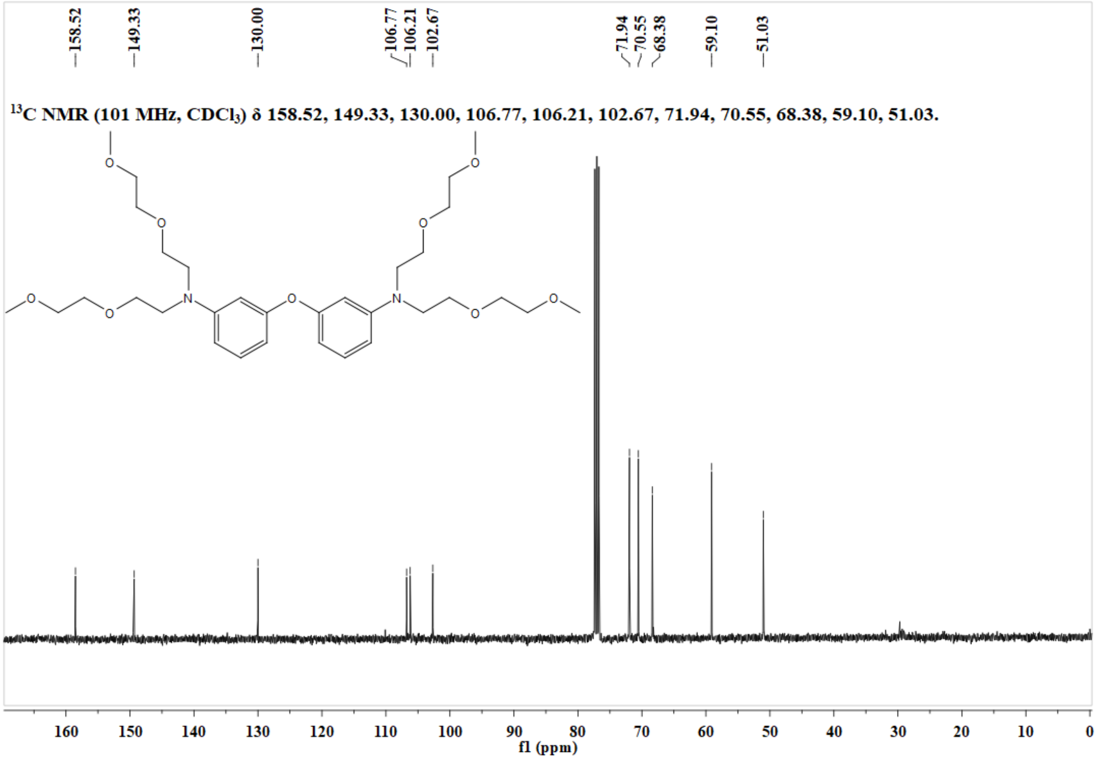

**Figure S19**. $^{13}C$ NMR spectrum of **O-DAEA-2** in $CDCl_3$.

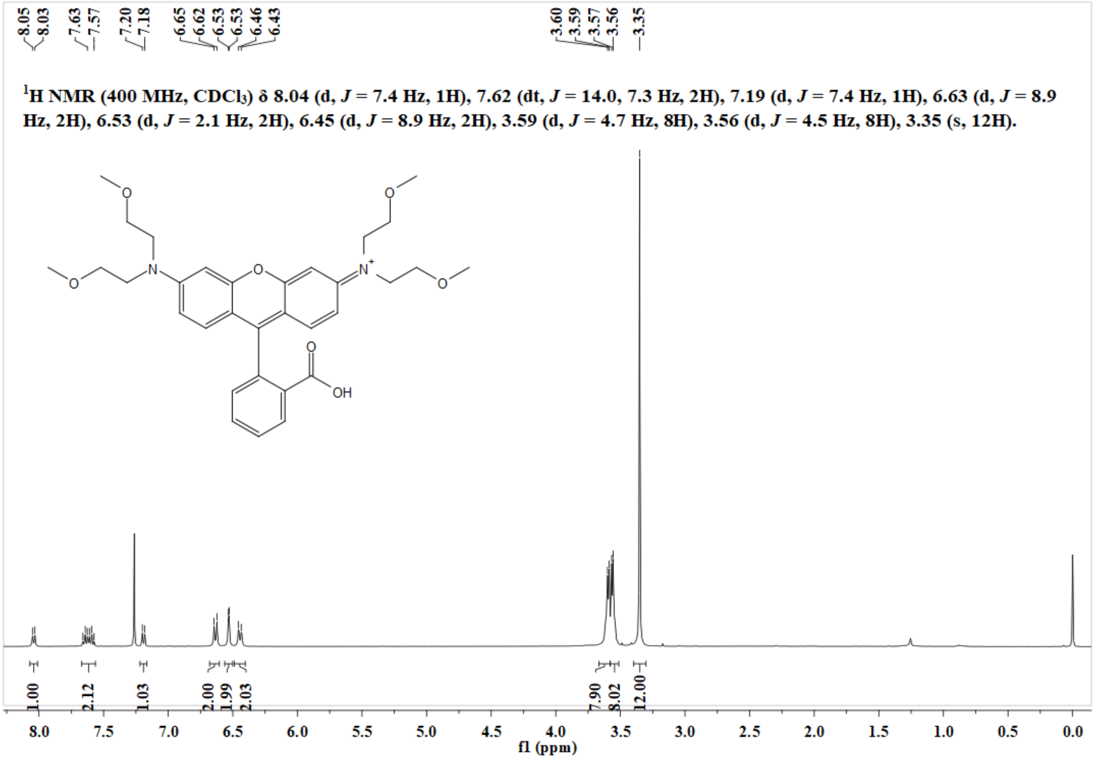

**Figure S20**. $^{1}$H NMR spectrum of **ER1** in $CDCl_3$.

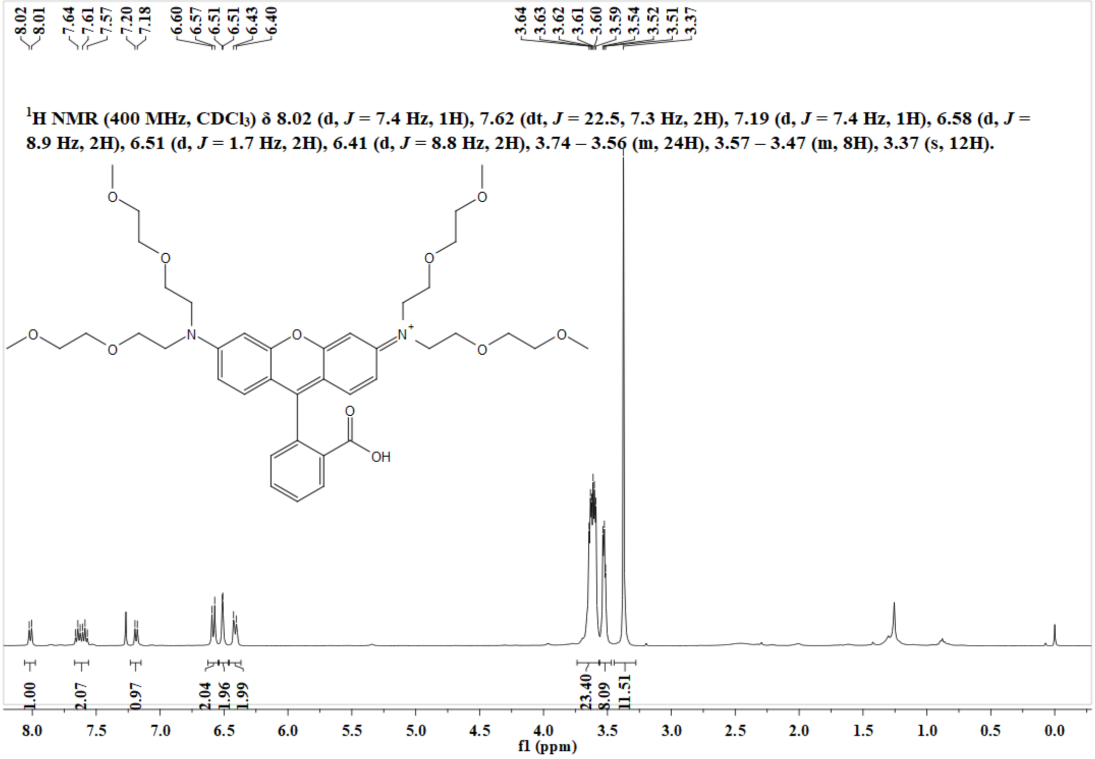

**Figure S21**. $^{1}$H NMR spectrum of **ER2** in $CDCl_3$.

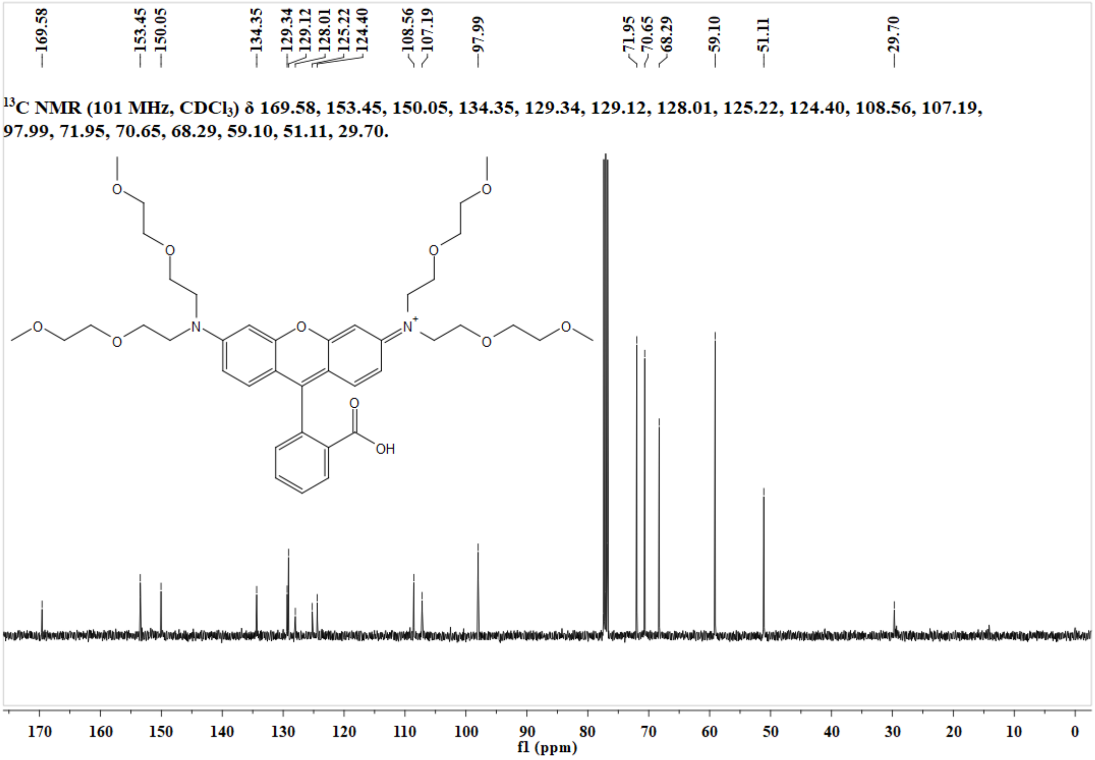

**Figure S22**. $^{13}C$ NMR spectrum of **ER2** in $CDCl_3$.

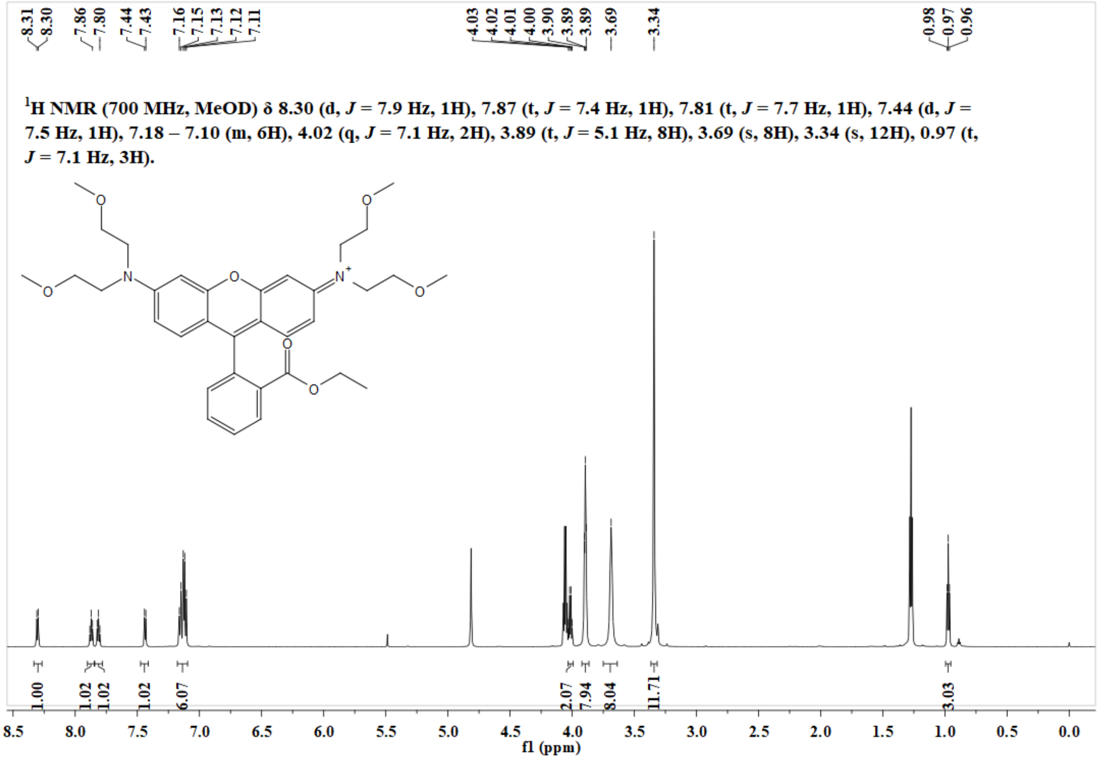

**Figure S23**. $^{1}H$ NMR spectrum of **ER1-Ester** in MeOD.

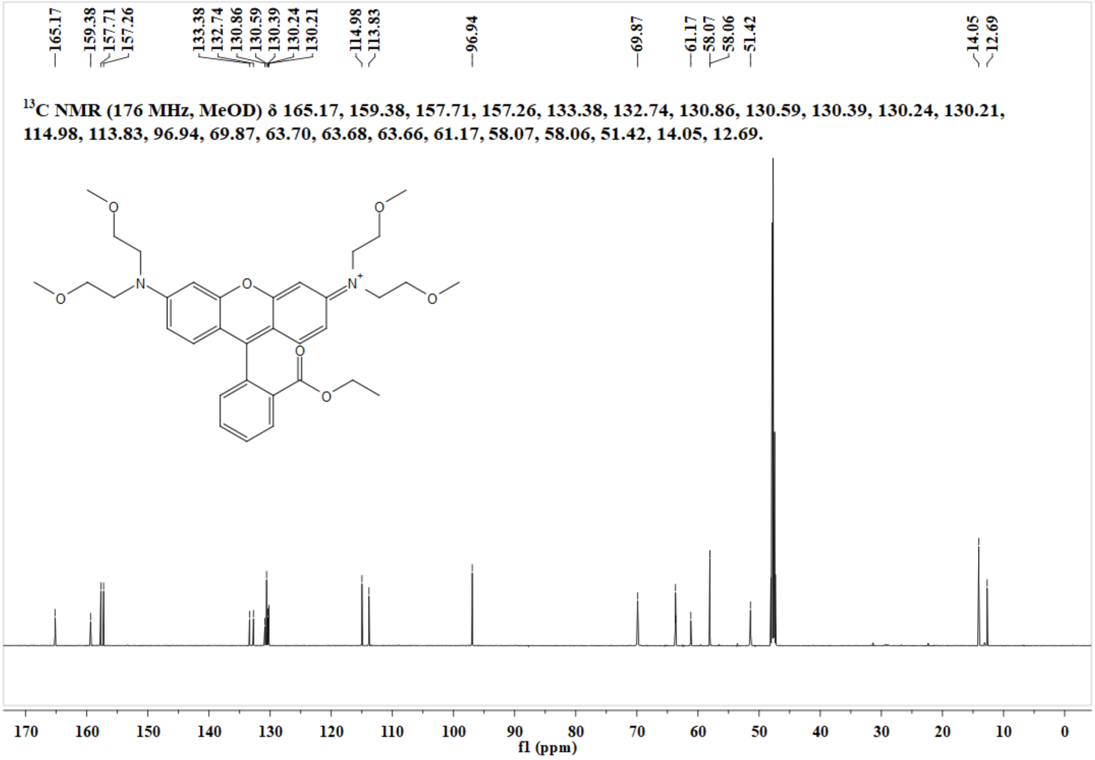

**Figure S24**. $^{13}$C NMR spectrum of **ER1-Ester** in MeOD.

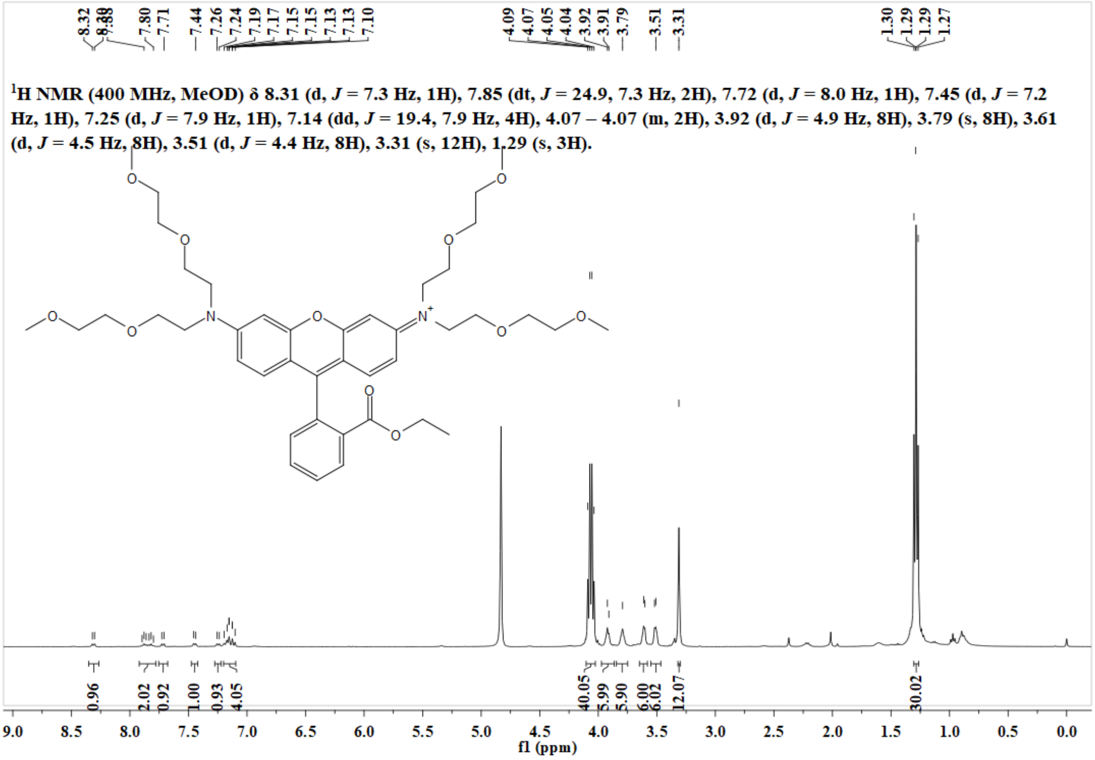

**Figure S25**. $^{1}$H NMR spectrum of **ER2-Ester** in MeOD.

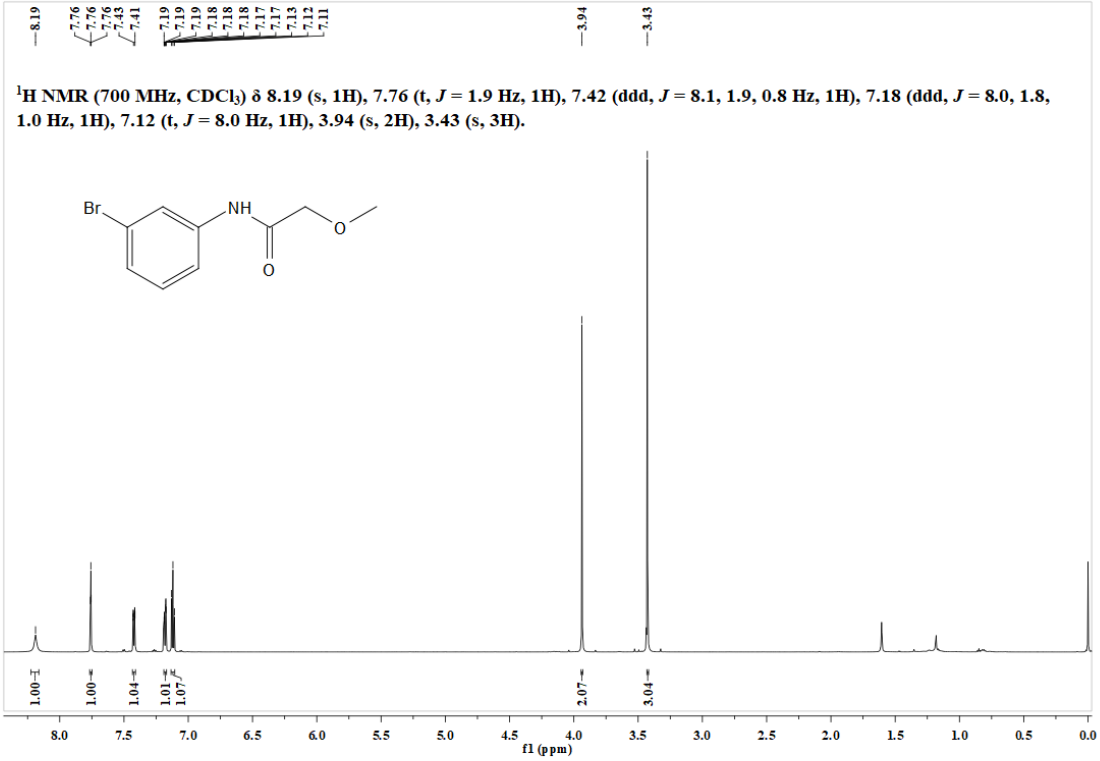

**Figure S26.** $^{1}$H NMR spectrum of compound **1** in $CDCl_3$.

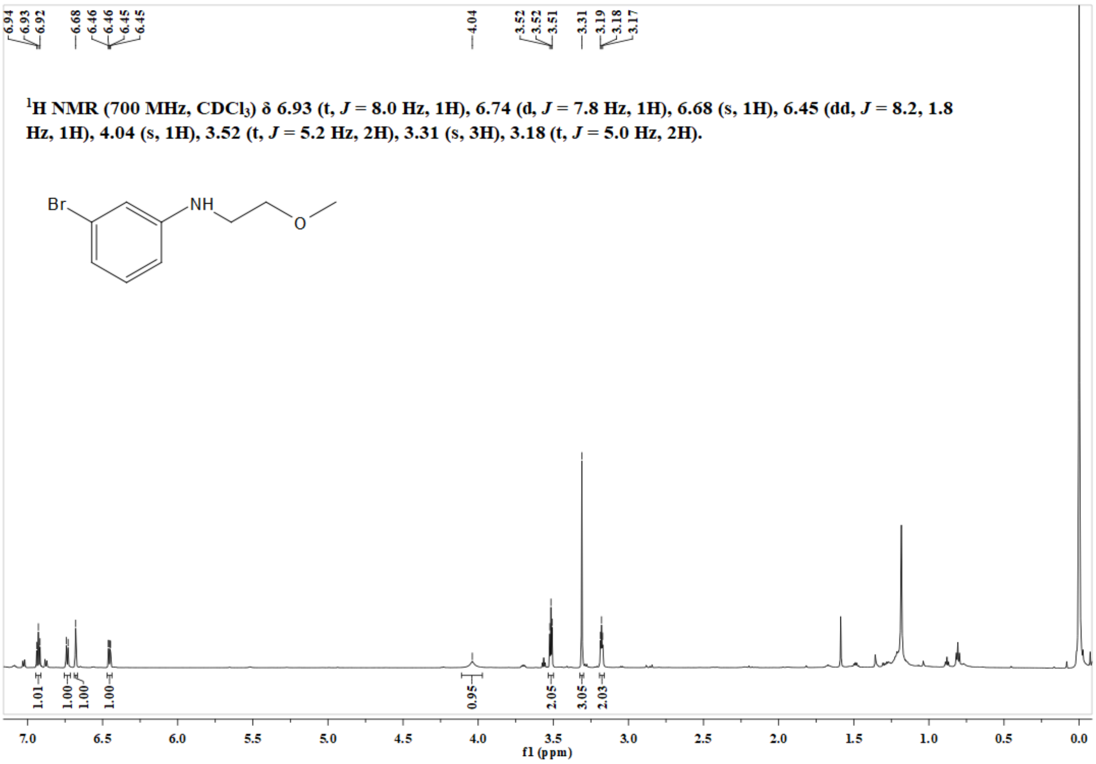

**Figure S27.** $^{1}$H NMR spectrum of compound **2** in $CDCl_3$.

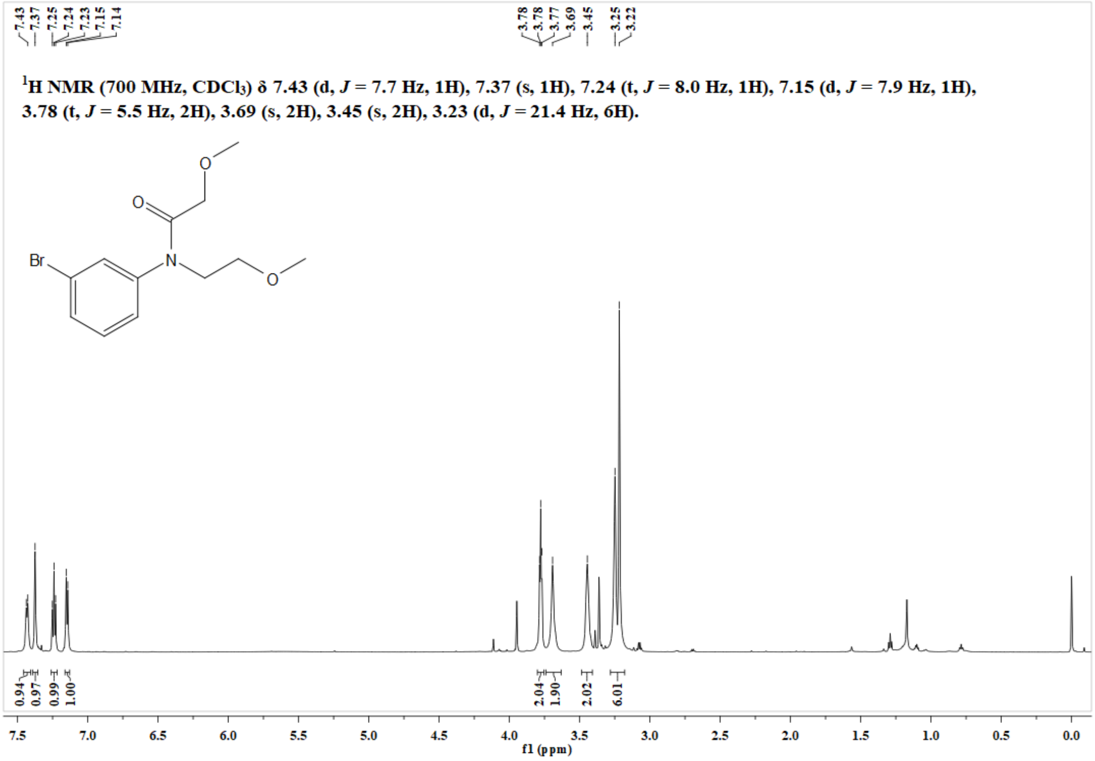

**Figure S28.** $^1$H NMR spectrum of compound **3** in $CDCl_3$.

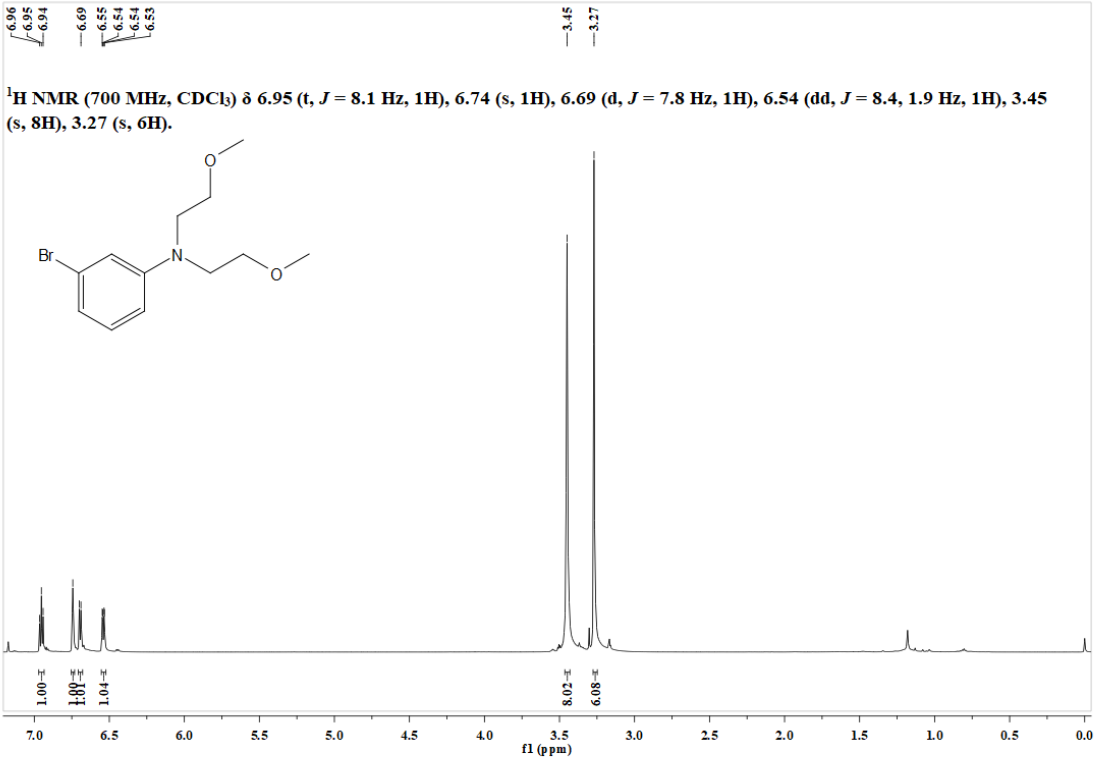

**Figure S29.** $^1$H NMR spectrum of compound **4** in $CDCl_3$.

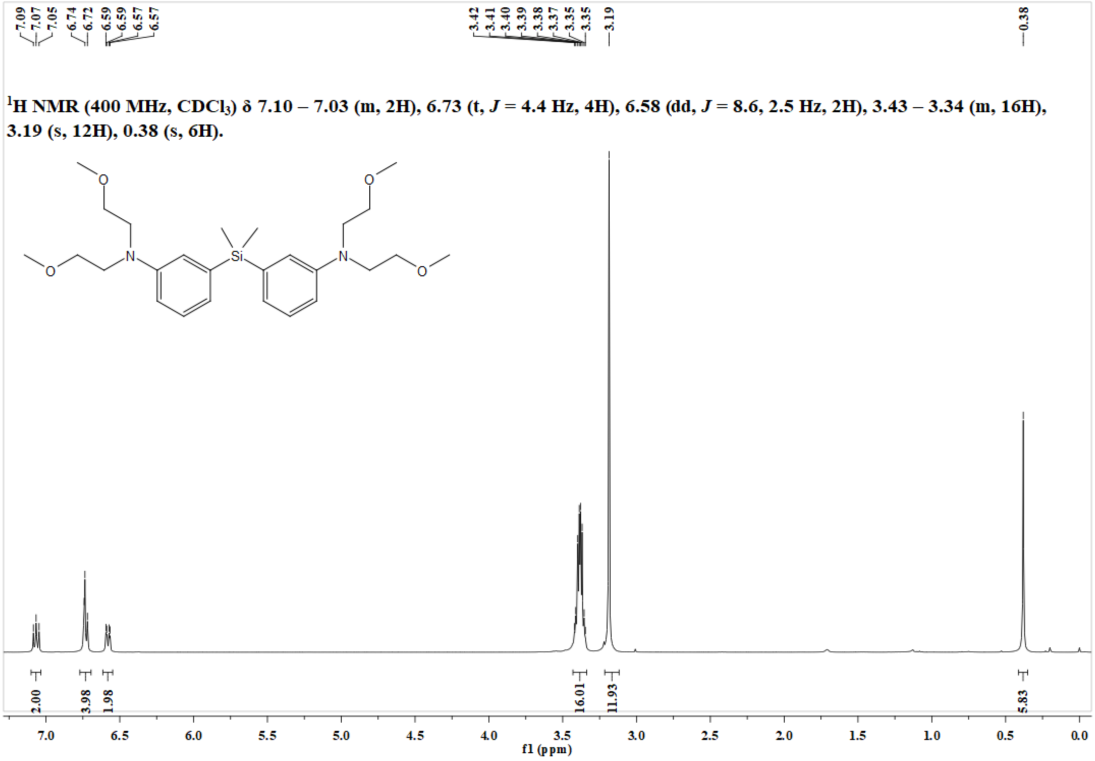

**Figure S30.** $^{1}$H NMR spectrum of **Si-DAEA** in $CDCl_3$.

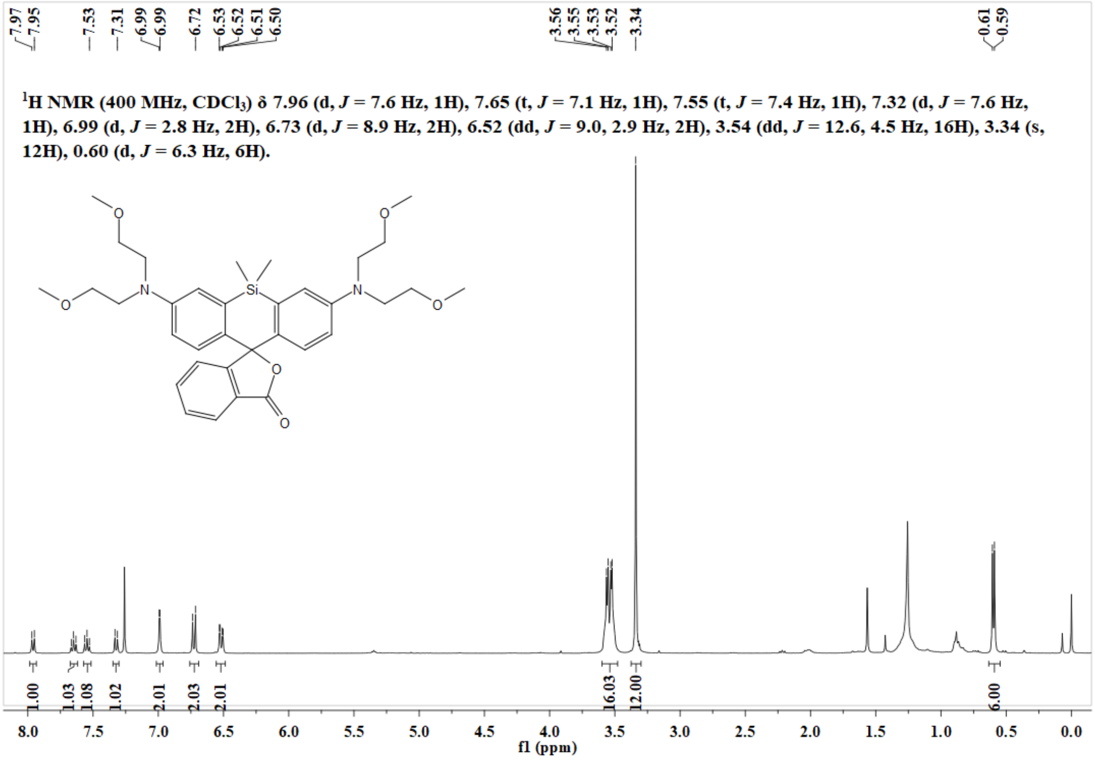

**Figure S31.** $^{1}$H NMR spectrum of **ESiR** in $CDCl_3$.

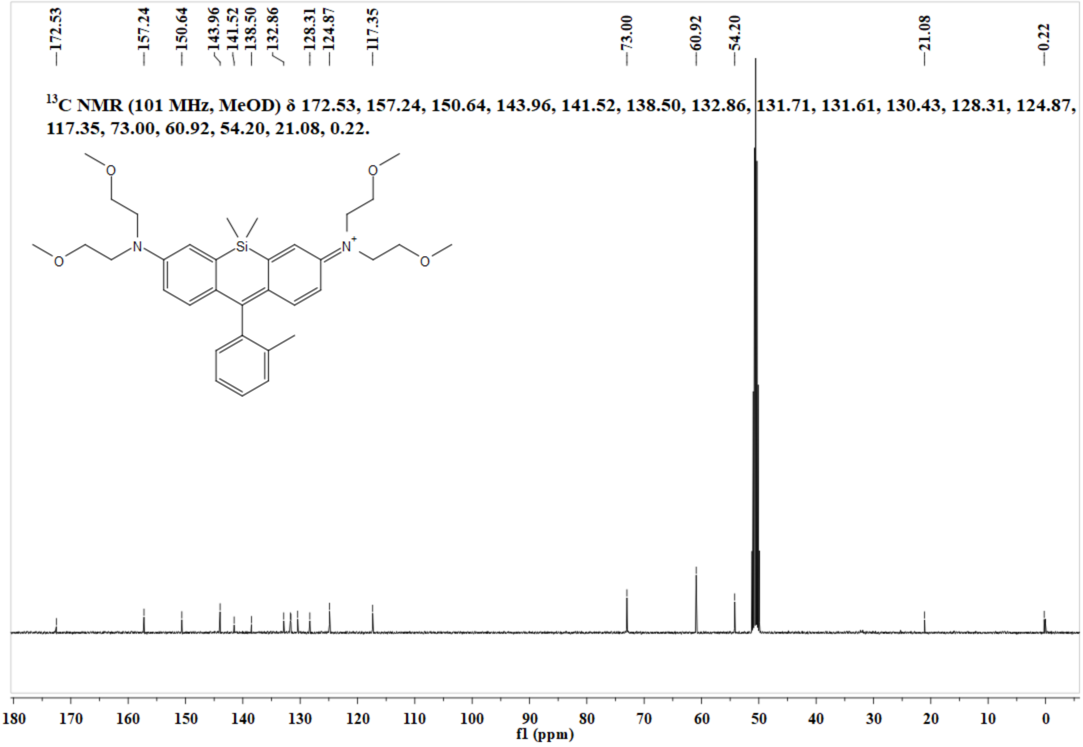

**Figure S32.** $^{13}$C NMR spectrum of **ESiR** in $CDCl_3$.

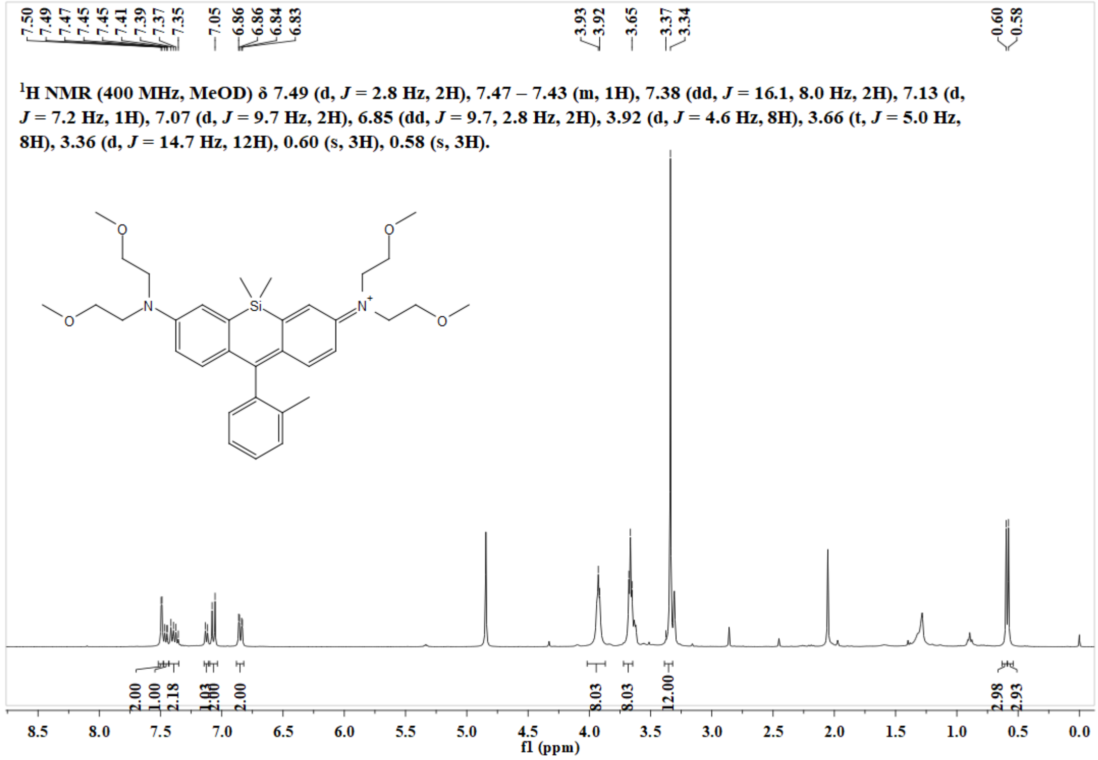

**Figure S33.** $^{1}$H NMR spectrum of **ESiR-Me** in MeOD.

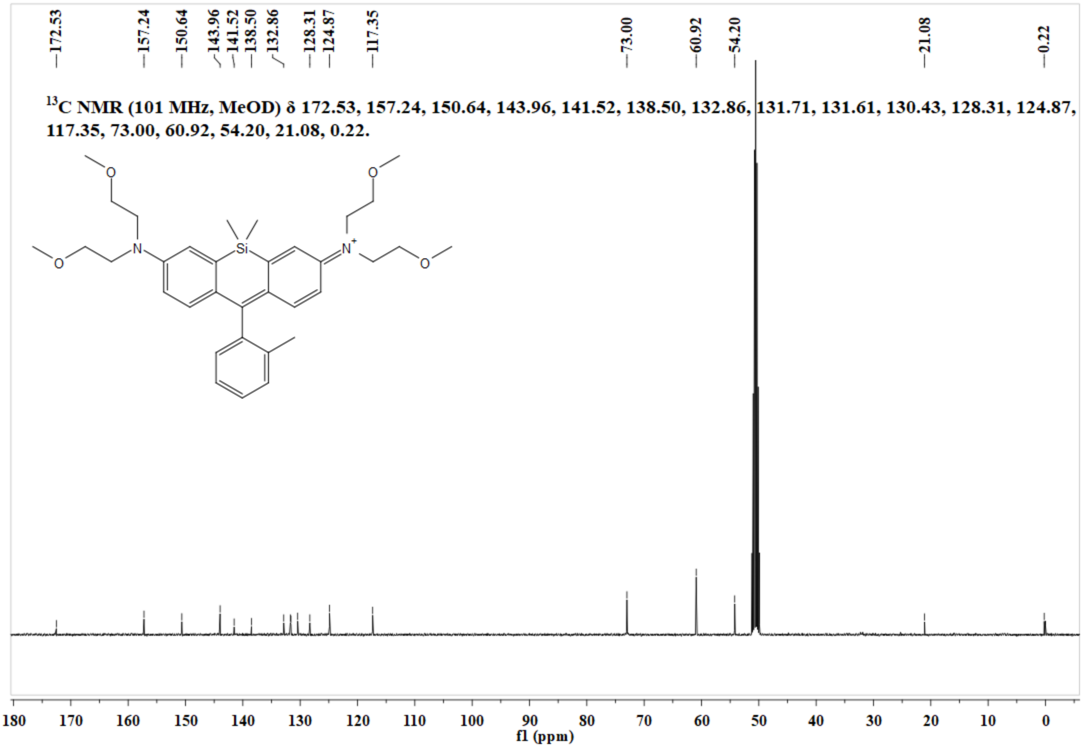

**Figure S34.** $^{13}C$ NMR spectrum of **ESiR-Me** in MeOD.

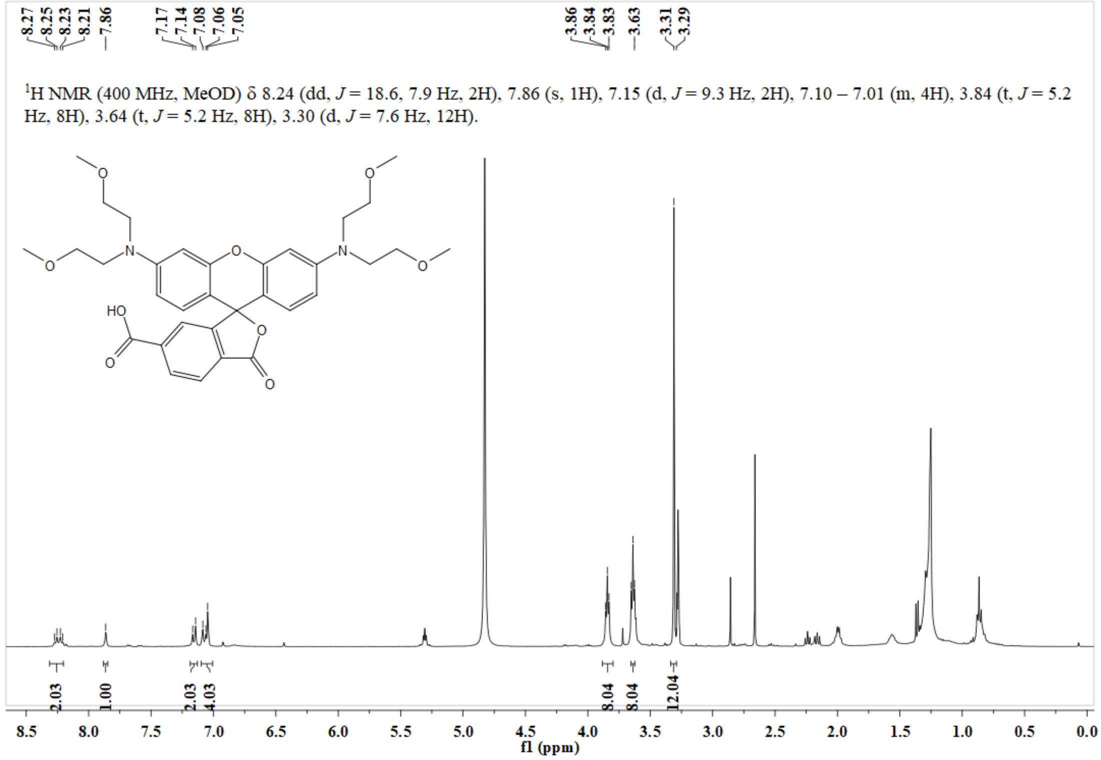

**Figure S35.** $^{1}H$ NMR spectrum of **ER-COOH** in MeOD..

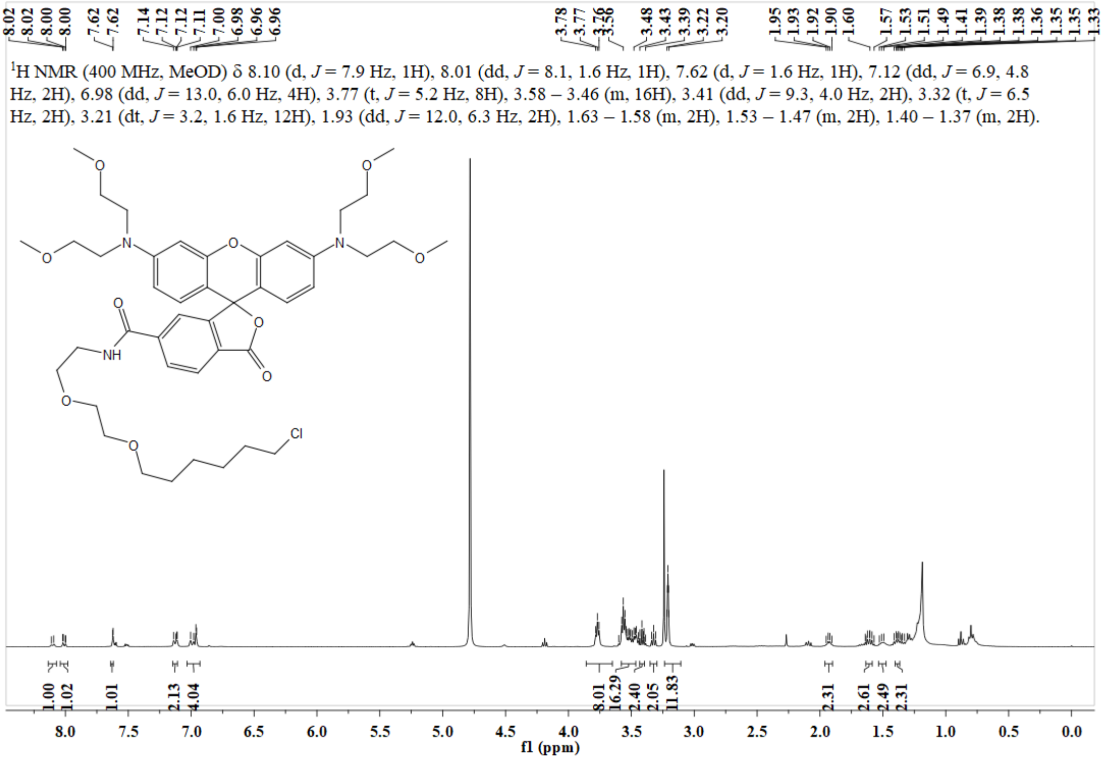

**Figure S36.** $^{1}$H NMR spectrum of **ER-Halo** in MeOD..

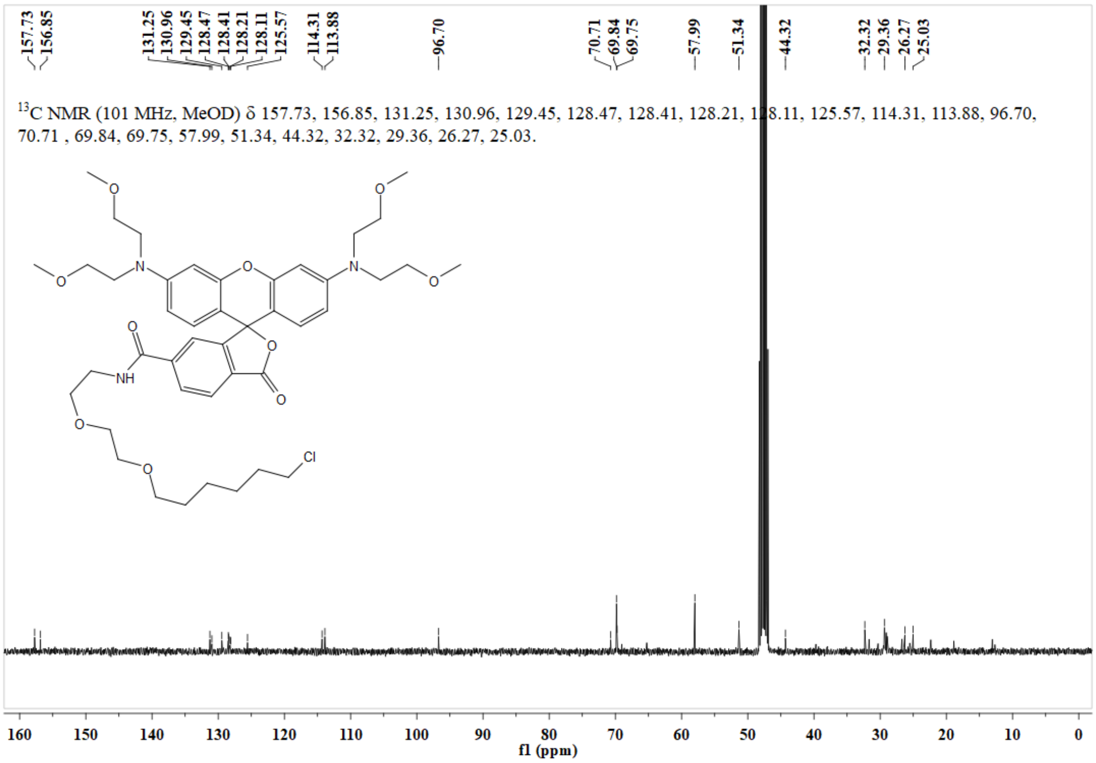

**Figure S37.** $^{13}$C NMR spectrum of **ER-Halo** in MeOD..

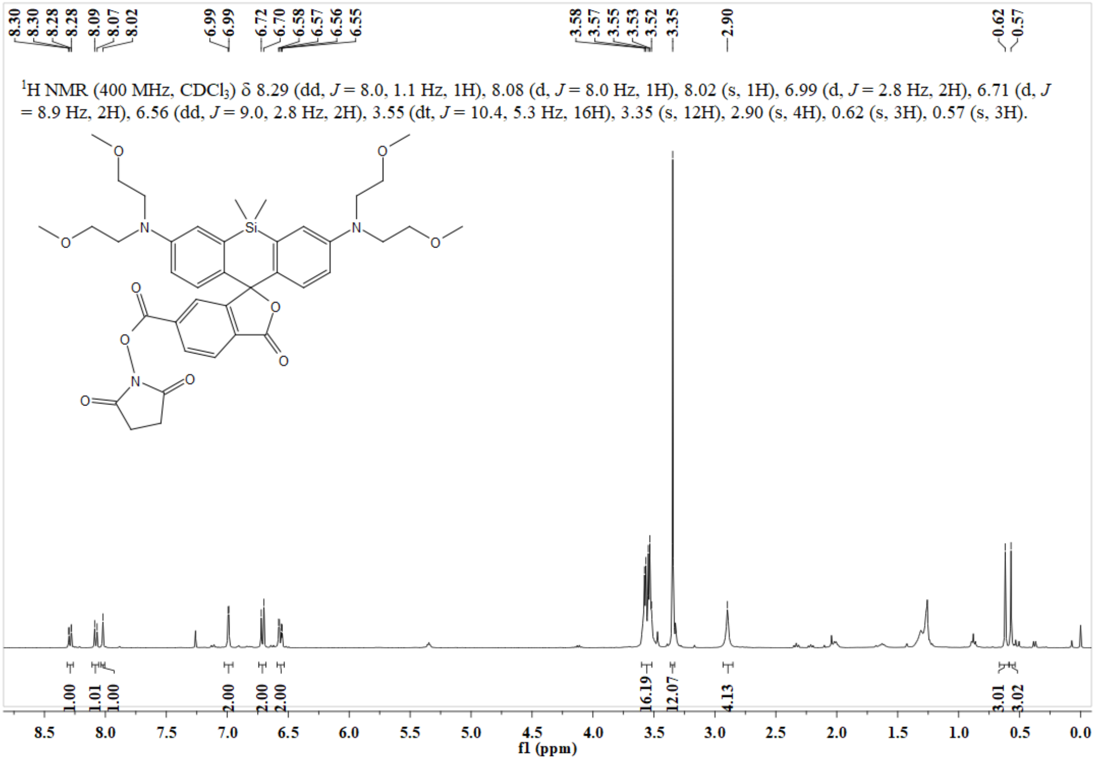

**Figure S38.** $^{1}$H NMR spectrum of **ESiR-NHS** in $CDCl_3$.

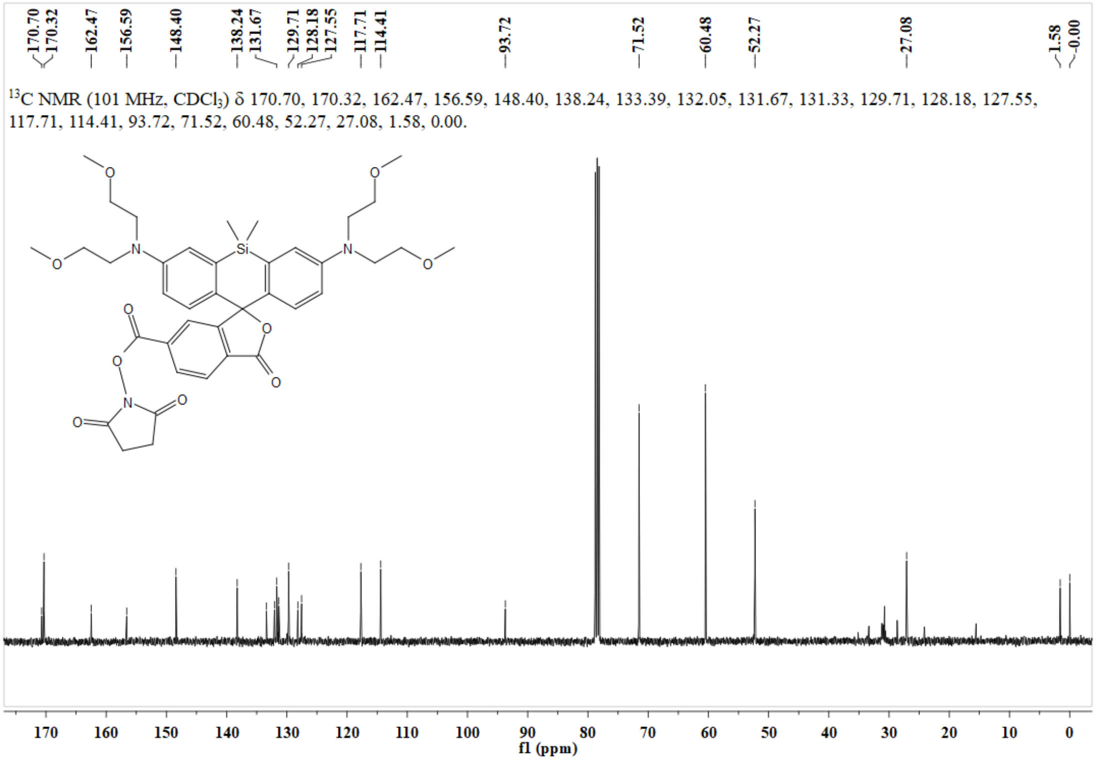

**Figure S39.** $^{13}$C NMR spectrum of **ESiR-NHS** in $CDCl_3$.

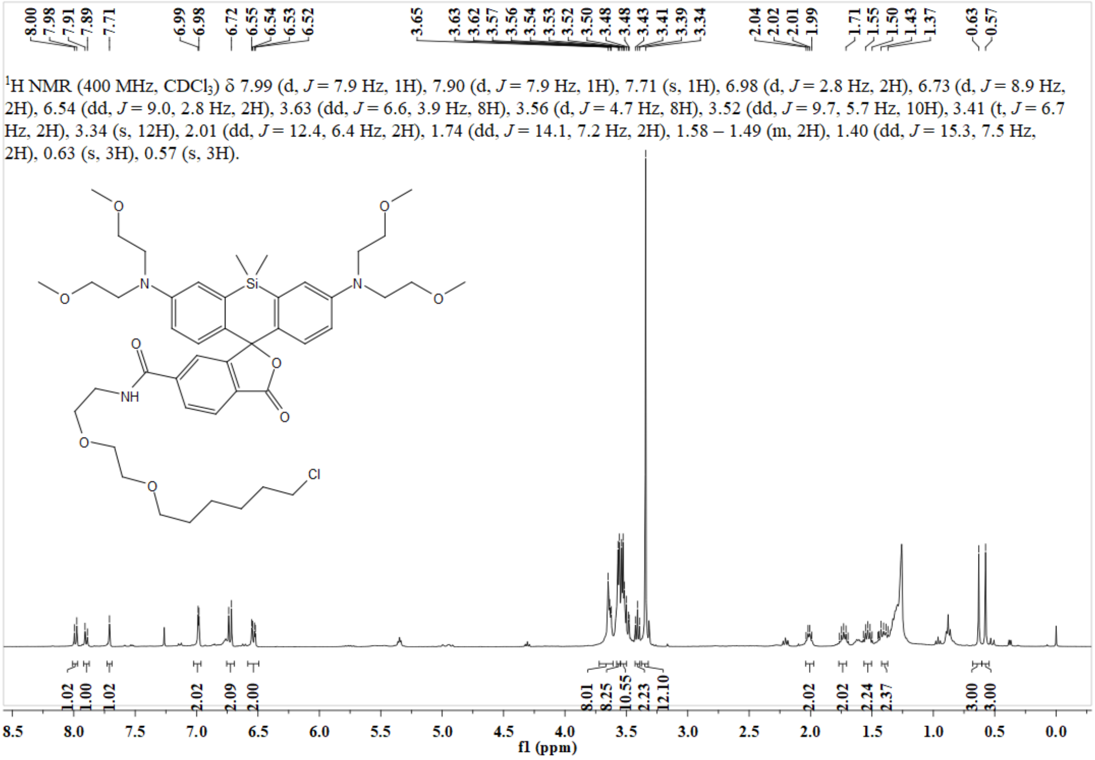

**Figure S40.** $^{1}$H NMR spectrum of **ESiR-Halo** in $CDCl_3$.

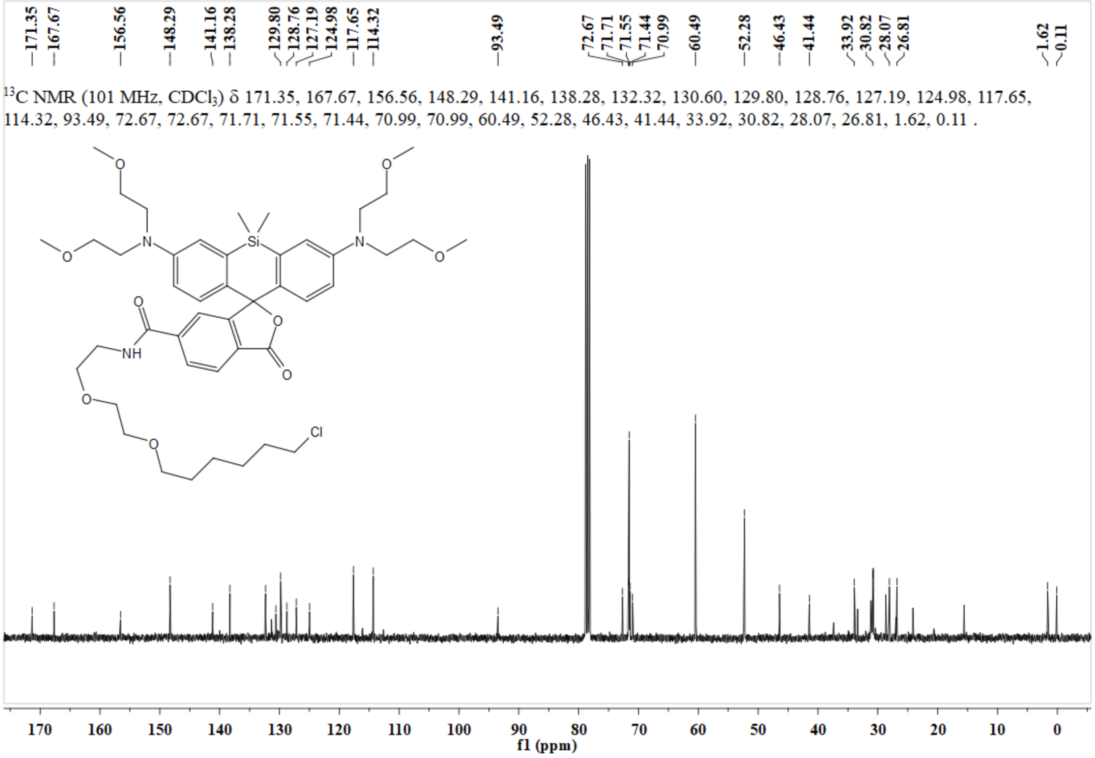

**Figure S41.** $^{13}$C NMR spectrum of **ESiR-Halo** in $CDCl_3$.

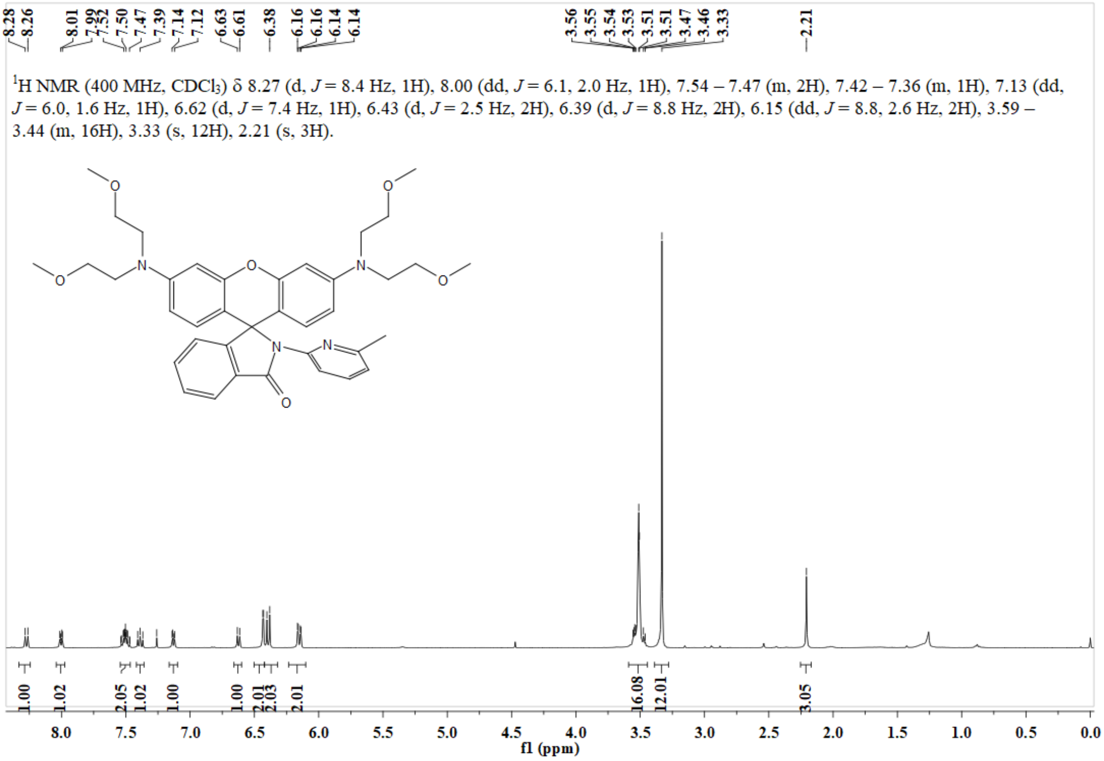

**Figure S42.** $^{1}H$ NMR spectrum of **Lyso-ER** in $CDCl_3$**.**

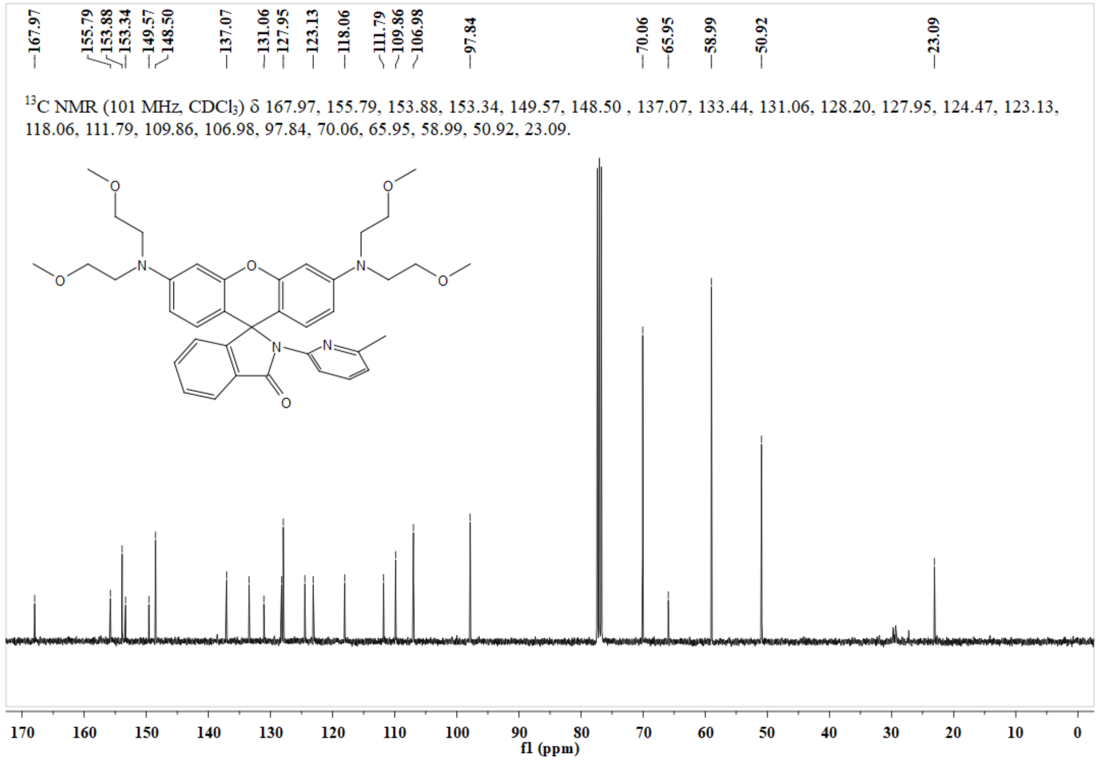

**Figure S43.** $^{13}C$ NMR spectrum of **Lyso-ER** in $CDCl_3$**.**

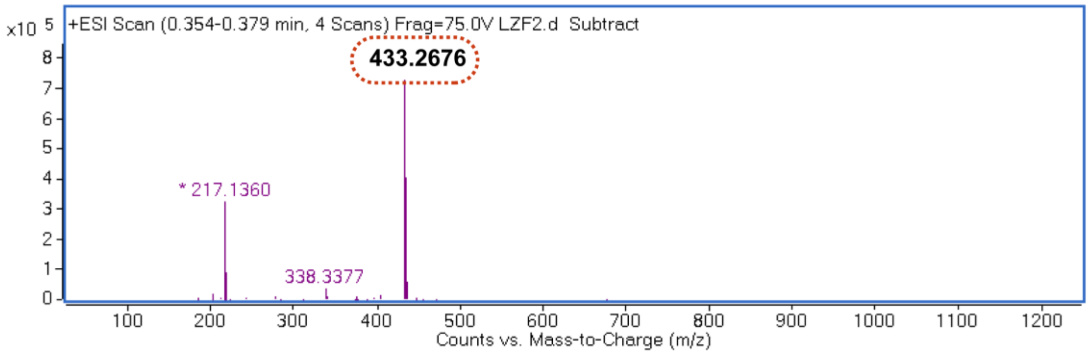

**Figure S44.** HRMS spectrum of **O-DAEA-1**.

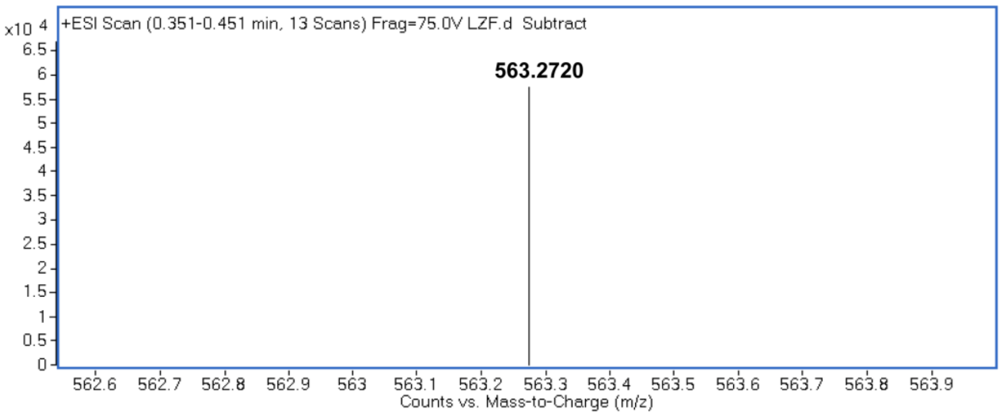

**Figure S45.** HRMS spectrum of **ER-1**.

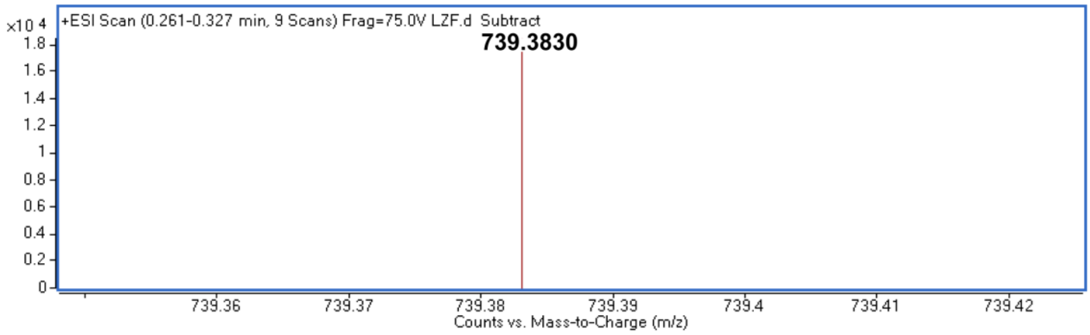

**Figure S46.** HRMS spectrum of **ER-2**.

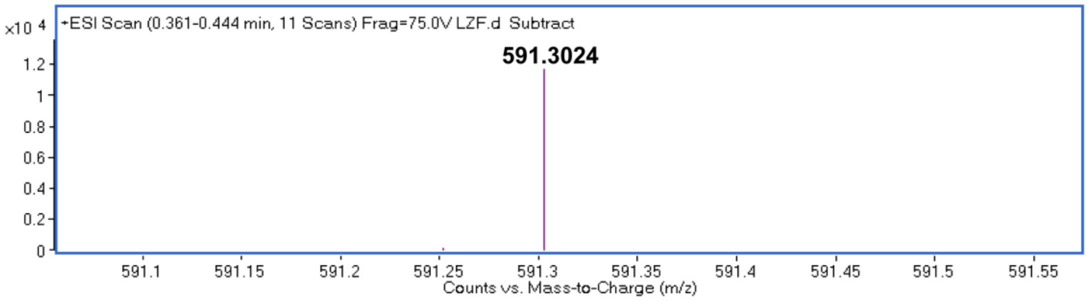

**Figure S47.** HRMS spectrum of **ER1-Ester**.

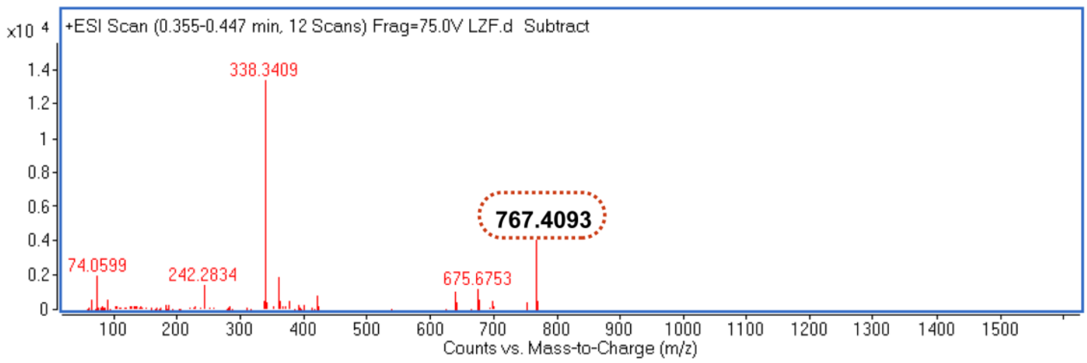

**Figure S48.** HRMS spectrum of **ER2-Ester**.

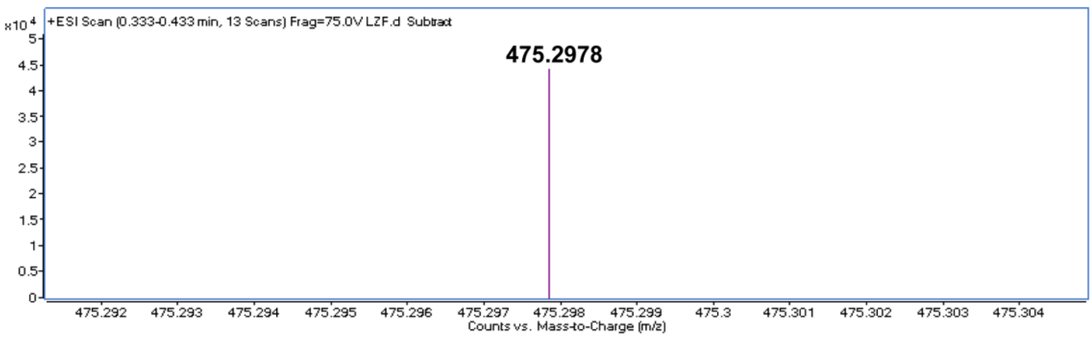

**Figure S49.** HRMS spectrum of **Si-DAEA**.

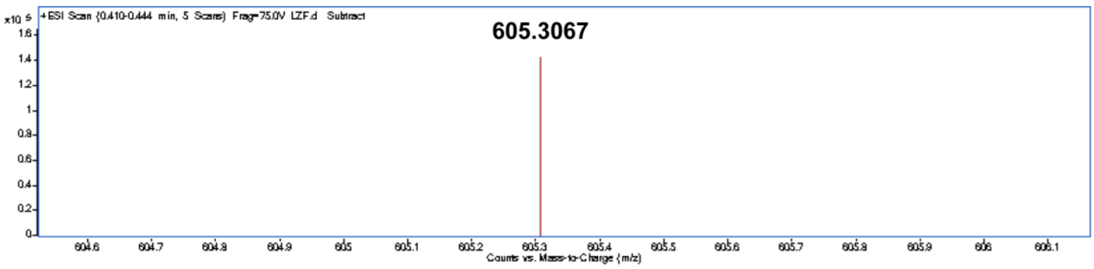

**Figure S50.** HRMS spectrum of **ESiR**.

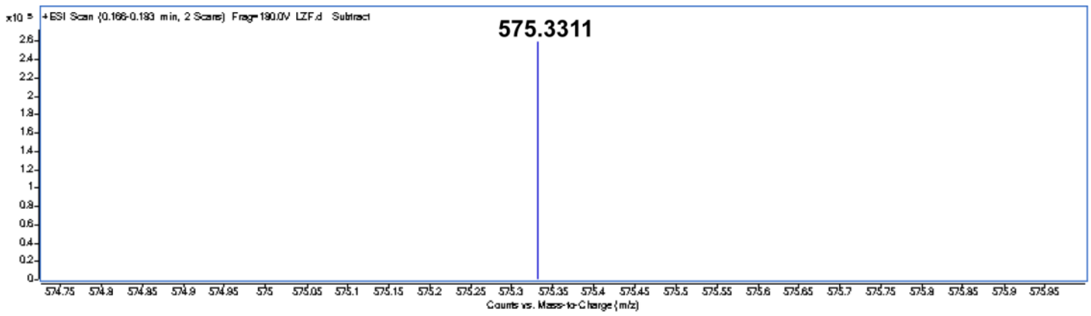

**Figure S51.** HRMS spectrum of **ESiR-Me**.

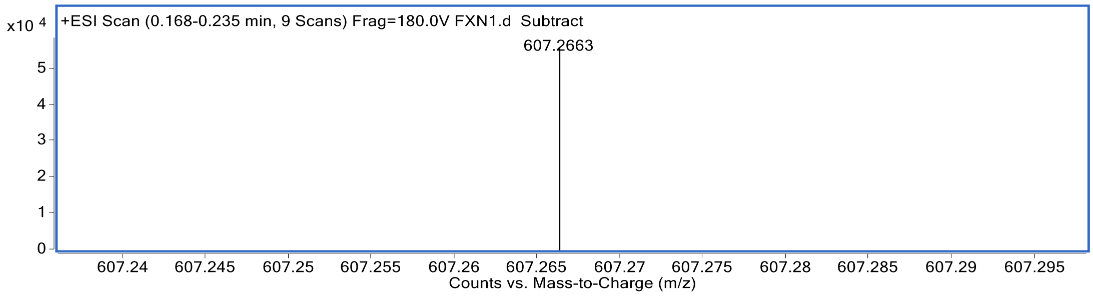

**Figure S52.** HRMS spectrum of **ER-COOH**

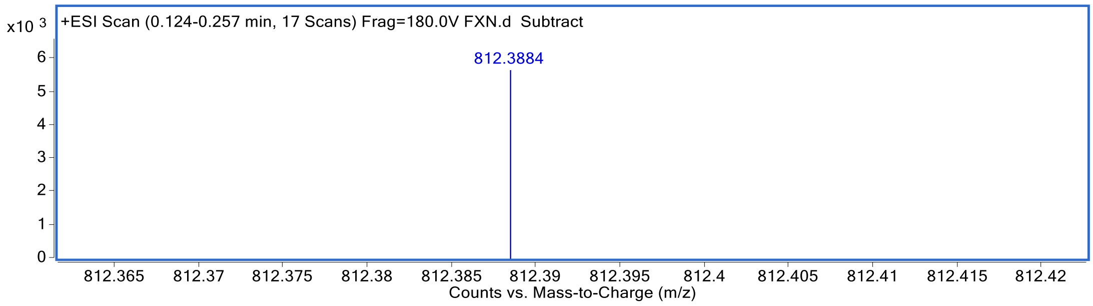

**Figure S53.** HRMS spectrum of **ER-Halo**.

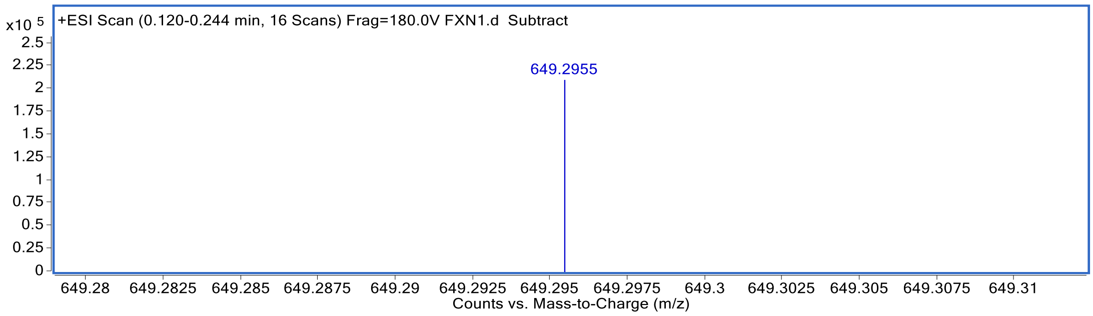

**Figure S54.** HRMS spectrum of **ESiR-COOH**.

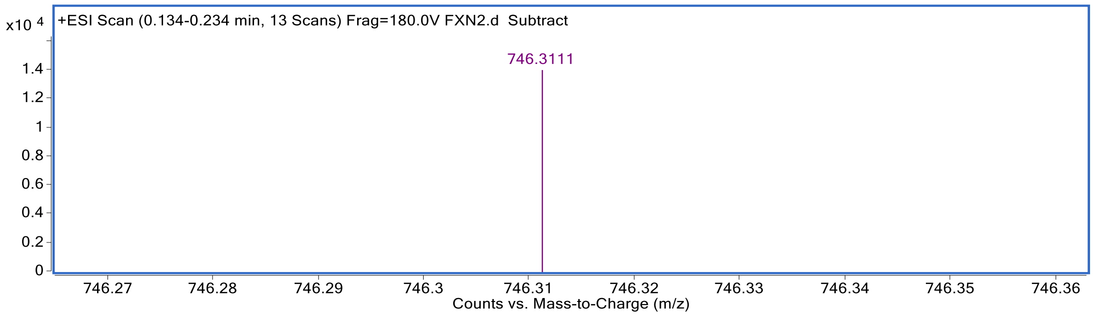

**Figure S55.** HRMS spectrum of **ESiR-NHS**.

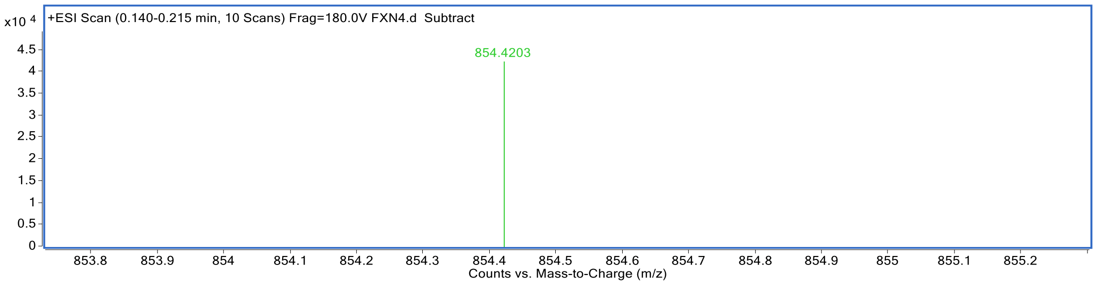

**Figure S56.** HRMS spectrum of **ESiR-Halo**.

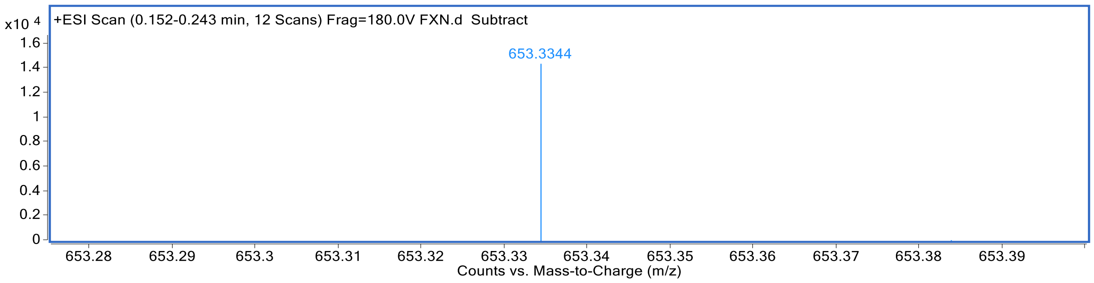

**Figure S57.** HRMS spectrum of **Lyso-ER**.
